# Hiding in plain sight: structure and sequence analysis reveals the importance of the antibody DE loop for antibody-antigen binding

**DOI:** 10.1101/2020.02.12.946350

**Authors:** Simon P. Kelow, Jared Adolf-Bryfogle, Roland L. Dunbrack

## Abstract

Antibody variable domains contain “complementarity determining regions” (CDRs), the loops that form the antigen binding site. CDRs1-3 are recognized as the canonical CDRs. However, a fourth loop sits adjacent to CDR1 and CDR2 and joins the D and E strands on the antibody v-type fold. This “DE loop” is usually treated as a framework region, even though mutations in the loop affect the conformation of the CDRs and residues in the DE loop occasionally contact antigen. We analyzed the length, structure, and sequence features of all DE loops in the Protein Data Bank, as well as millions of sequences from HIV-1 infected and naïve patients. We refer to the DE loop as H4 and L4 in the heavy and light chain respectively. Clustering the backbone conformations of the most common length of L4 (6 residues) reveals four conformations: two κ-only clusters, one λ-only cluster, and one mixed κ/λ cluster. The vast majority of H4 loops are length-8 and exist primarily in one conformation; a secondary conformation represents a small fraction of H4-8 structures. H4 sequence variability exceeds that of the antibody framework in naïve human high-throughput sequences, and both L4 and H4 sequence variability from λ and heavy germline sequences exceed that of germline framework regions. Finally, we identified dozens of structures in the PDB with insertions in the DE loop, all related to broadly neutralizing HIV-1 antibodies, as well as antibody sequences from high-throughput sequencing studies of HIV-infected individuals, illuminating a possible role in humoral immunity to HIV-1.

## Introduction

Antibodies utilize three hypervariable loops on each variable domain to bind antigens. These three loops are referred to as complementarity determining regions or CDRs, and were first identified by their high sequence variation relative to the rest of the variable domain sequence (1). However, there is a fourth loop, structurally adjacent to CDR1 and CDR2 and referred to as the DE loop, which joins strands D and E in the immunoglobulin v-type fold (2,3) (Figure 1). In the linear sequence, the DE loop sits between CDRs 2 and 3 and is encoded by V-region gene segments (4). The DE loop has been traditionally considered part of the antibody framework, so studies addressing the ability of specific DE loop residues to affect antibody binding (5–8) have addressed these residues as framework residues, and not part of a CDR-like loop. However, mutations in the DE loop can affect antigen binding and in some structures it directly contacts antigen.

**Figure 1:**
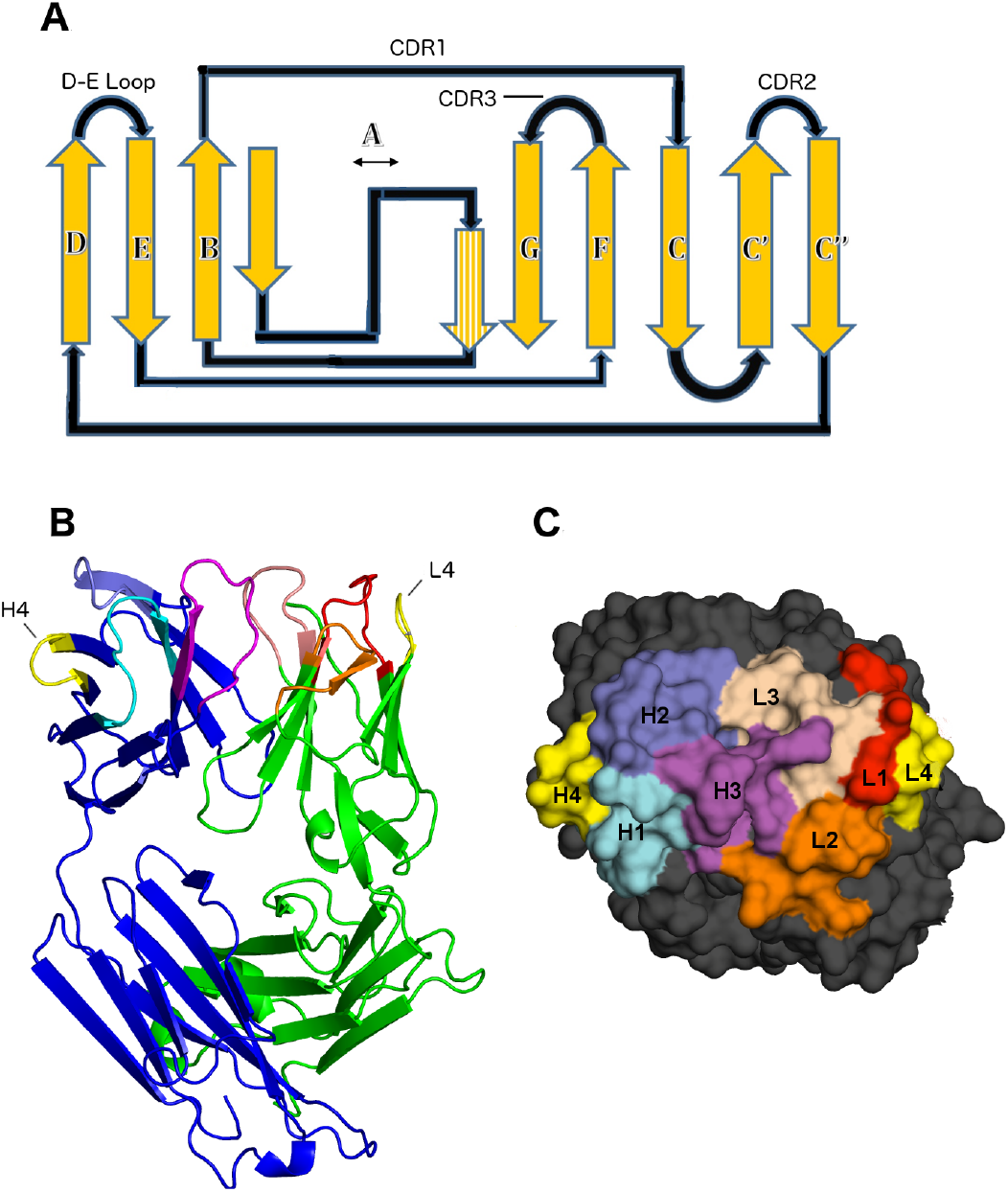
Position of the DE loop in antibody structures. **A.** V-type fold according to Bork et. al. The DE loop and CDRs are indicated. Strand A forms beta-strand pairing interactions with both strand B and strand G. **B.** Example of antibody Fab fragment (light chain in green, heavy chain in blue). **C.** Top-down view of antibody combining site. The canonical CDRs and the DE loop are marked in panel C and are represented in the same colors in panel B.

Chothia and Lesk first noted hydrophobic packing interactions of the DE loop with L1, in particular that VL residue 87 (IMGT numbering; Chothia residue 71) packs against L1, and is typically either Phe or Tyr (5). Foote and Winter demonstrated that some antibodies lose binding affinity to target antigen upon mutation of this residue from Tyr to Phe, noting that this interaction mediates interaction of L1 with target antigen though a hydrogen bond between Tyr87 and Asn37 (7). Al-Lazikani et al. observed a switch in conformation of CDR L1 of length 11 when Tyr87 changes to Phe87 (8). Tramontano et al. noted that an Arg residue at IMGT VH residue 80 in the heavy chain (Chothia VH residue 71) makes hydrogen bonds to H1 and H2 and packs against side-chain residues of H1 and H2, stabilizing specific H2 conformations and bringing H1 and H2 into closer contact with each other (6). Several studies since these initial observations have considered various mutations of DE loop residues, with particular focus on VH residue 80, and successfully engineered significant differences in both antibody stability or antibody-antigen affinity (9–13). However, the effects (or lack thereof) of these mutations are unpredictable, and appear to vary with the germline construct of the antibody.

Previously we demonstrated the importance of the DE loop in redesigning an unstable anti-EGFR antibody, C10, and its affinity-matured form P2224 (14). The VL region of C10 appeared to be a fusion of λ3 and λ1 V-region gene loci, introduced most likely through PCR amplification. We redesigned the antibody framework in an attempt to stabilize the antibody and prevent antibody aggregation by grafting the sequences of the λ antibody L1, L2, and L3 CDRs onto a κ framework. We observed that the λ DE loop was different in structure and sequence from a typical κ DE loop in antibodies. Grafting the DE loop along with L1, L2, and L3 from the P2224 λ antibody onto a κ framework produced an antibody with significantly increased thermostability, while also retaining P2224’s binding affinity. As a control, grafting L1, L2, and L3 while keeping the host κ DE loop sequence produced an antibody with lower stability and significantly reduced affinity.

In this paper, we analyze the structures and sequences of the DE loops of heavy and light chain variable domains in the Protein Data Bank (PDB), along with a large set of sequences from multiple high-throughput antibody sequencing studies. We first define the DE loop (which we refer to as L4 on the light chain and H4 on the heavy chain) as IMGT residues 80-87 based on the structural variability observed in Ramachandran maps of residues encompassing the D and E strands of the heavy and light chains in the PDB. With these definitions, we expand on the observations presented by Lehmann et al. by clustering the backbone conformations of L4 and H4 loops in the structures of antibodies to address their structural contribution to antigen binding. The vast majority of L4 loops are of length 6, the exceptions being human λ5 and λ6, mouse λ4-λ8, rat λ2 and λ3, and rabbit λ5 and λ6 DE loops, which are length 8. All human and mouse germline H4 loops are of length 8. Some rabbit and llama H4 sequences are of length 7.

From a clustering of the conformations of L4 and H4 with validated electron density (15), we demonstrate that L4 loops of length 6 exist in four dominant conformations, two of which only contain κ antibodies, one of which contains only λ antibodies, and one of which contains both κ and λ antibodies. H4 length-8 structures have one primary conformation, as well as a secondary conformation that represents a small fraction of the H4 length-8 structures. The primary heavy chain H4 cluster and L4 length-8 structures have very similar backbone conformations. In addition to classifying the structures of L4 of lengths 6 and 8 and H4 of lengths 6, 7, and 8, we also calculate all hydrogen bond interactions between the DE loop and the CDRs that influence the conformation of these CDRs. We also correlate the structural features with antibody germline identity as defined by the IMGT database (4).

Finally, we examine 125 structures in the PDB that have insertions in the heavy or light chain DE loops compared to their germlines. With only one exception, these all turn out to be structures of broadly neutralizing antibodies (bnAbs) of HIV-1 in human patients (16–52). From these structures, we identify insertions in the DE loops on both the light and heavy chains that contact the antigen gp120, and compare the binding contribution of the DE loops with insertions to the rest of the CDRs in these antibodies. From sequencing studies of HIV-1 infected individuals, we identify insertions and deletions, hypersomatic mutation, and frameshift mutations in and around the DE loop region for the light and heavy chain. The insertion and frameshift mutations are not observed in a large set of antibodies from uninfected individuals, and thus may represent a mechanism in humoral immunity to HIV-1.

## Results

### Clustering of canonical length L4 and H4 structures

To define the regions of structural variability in the vicinity of both H4 and L4, we plotted the ϕ and ψ dihedrals of the D and E strands and the residues in between for all heavy and light chains of antibodies in the PDB with the most common L4 or H4 lengths of 6 and 8 residues respectively (Figure 2). We found that IMGT residues 77-79 and 86-90 uniformly occupy the beta region of the Ramachandran map, while there is some variability in residues 80-82 and 85 of light chains and residues 81-85 of heavy chains. So that the starting and ending residues are opposite each other in the beta strands and because residues 80 and 87 contact CDR1 as noted above, we define the DE loop as IMGT residues 80-87. Kabat and Chothia number the H4 region as residues 71-78 and L4 loops of length 6 as residues 66-71. L4 loops of length 8 would require insertion codes, such as 68A, 68B. In the rest of this paper, we number the residues in the DE loops from 1 to N for DE loops of length N, such that L4 loops of length 6 are numbered 1-6, and L4 and H4 loops of length 8 are numbered 1-8. A mapping of our residue numbering to those of IMGT, Kabat, Chothia, and Honegger-Plückthun is presented in Table 1.

**Figure 2.**
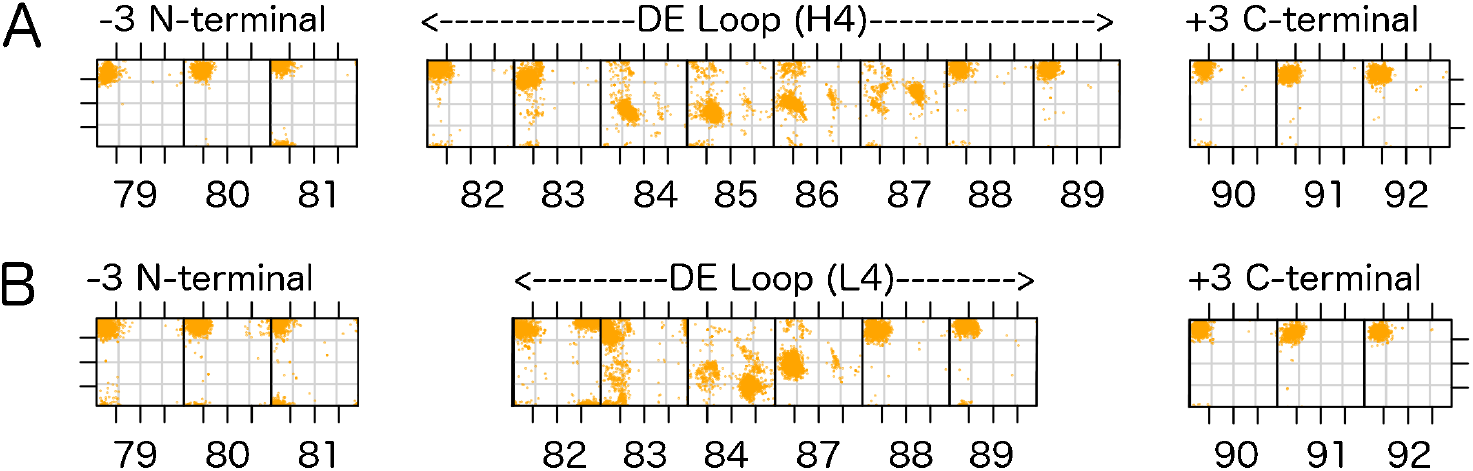
Ramachandran plots for part of the D strand, the DE loop, and part of the E strand (IMGT residues 77-90) for the most common DE loop lengths. **A.** ϕ (x-axis) and ψ (y-axis) for residues in length-8 H4 loops, and the 3 anchor residues before and after the loop. IMGT residue numbers are provided at the bottom of each panel. **B.** ϕ,ψ plots for residues in length-6 L4 loops, and the 3 anchor residues before and after the loop

**Table 1.** Map between various numbering schemes within H4 and L4 loops.

| IMGT | AHo | Position in H4-6 | Position in H4-7 | Position in H4-8 | Chothia/Kabat Heavy Chain | Position in L4-6 | Position In L4-8 | Chothia/Kabat Light Chain |
| --- | --- | --- | --- | --- | --- | --- | --- | --- |
| 80 | 82 | 1 | 1 | 1 | 71 | 1 | 1 | 66 |
| 81 | 83 | 2 | 2 | 2 | 72 | 2 | 2 | 67 |
| 82 | 84 | 3 | 3 | 3 | 73 | 3 | 3 | 68 |
| 83 | 85 |  | 4 | 4 | 74 |  | 4 | 68A |
| 84 | 86 |  |  | 5 | 75 |  | 5 | 68B |
| 85 | 87 | 4 | 5 | 6 | 76 | 4 | 6 | 69 |
| 86 | 88 | 5 | 6 | 7 | 77 | 5 | 7 | 70 |
| 87 | 89 | 6 | 7 | 8 | 78 | 6 | 8 | 71 |

Mapping from various antibody numbering schemes to the numbering scheme used (1 to N where N is the length of the CDR loop considered. AHo indicates the Honegger-Plückthun numbering scheme

We clustered the structures of L4 loops with germline lengths 6 and 8 and H4 loops of length 6, 7, and 8 using a maximum dihedral angle metric described in Materials and Methods. We used a density-based clustering algorithm, DBSCAN (53) to identify and remove outliers and to identify common conformations within the data. Table 2 provides a summary of the L4 and H4 clusters, specifying their gene, consensus Ramachandran conformation, consensus sequence, number of chains in each cluster, fraction of PDB chains of that length in each cluster, number of unique sequences, and the average ϕ and ψ dihedral values for each residue in the DE loop for that cluster.

**Table 2.** DE loop canonical families.

| Gene | Cluster | Rama. string | Common sequence | # PDB chains (all) | # PDB chains ($\geq 0.75$ EDIA) | % chains (all) | % chains ($\geq 0.75$ EDIA) | Unique seqs (all) | Unique seqs ($\geq 0.75$ EDIA) | 1 | 2 | 3 | 4 | 5 | 6 | 7 | 8 |
| --- | --- | --- | --- | --- | --- | --- | --- | --- | --- | --- | --- | --- | --- | --- | --- | --- | --- |
| $\kappa$ | L4-6-1 | EBEABB | GSGTDF | 3,854 | 1,393 | 64.5 | 66.6 | 77 | 48 | 120, 170 | -164, 158 | 73, -100 | -118, -11 | -119, 122 | -132, 153 | | |
| $\lambda/\kappa^*$ | L4-6-2 | BBEABB | KSGTTA | 1,377 | 562 | 23.0 | 26.9 | 81 | 67 | -141, 136 | -144, 114 | 64, -120 | -100, 10 | -120, 136 | -115, 148 | | |
| $\kappa$ | L4-6-3 | BBAABB | GSGTDF | 95 | 48 | 1.6 | 2.3 | 7 | 4 | 163, 168 | -90, -144 | -88, -23 | -128, -13 | -125, 125 | -131, 151 | | |
| $\lambda$ | L4-6-4 | BBLLBB | LIGGKA | 31 | 14 | 0.5 | 0.7 | 6 | 5 | -110, 130 | -138, 123 | 51, 48 | 70,, 12 | -124, 156 | -86, 145 | | |
| $\lambda/\kappa$ | noise | - | - | 622 | 76 | 10.4 | 3.5 | 74 | 29 | - | - | - | - | - | - | | |
| $\lambda 5/\lambda 6$ | L4-8-1 | BBAAALBB | IDSSNSA | 90 | 40 | 100.0 | 100.0 | 8 | 6 | -128, 135 | -120, 100 | -65, -31 | -66, -34 | -101, 2 | 54, 44 | -134, 156 | -111, 151 |
| H | H4-6-1 | BBAABB | RTSTTV | 35 | 21 | 76.1 | 100.0 | 4 | 3 | -134, 152 | -119, -172 | -64, -34 | -124, 0 | -138, 158 | -127, 133 |  |  |
| H | noise | - | - | 11 | 0 | 23.9 | 0.0 | 5 | 0 | - | - | - | - | - | - | - | - |
| H | H4-7-1 | BABAABB |  | 37 | 17 | 100.0 | 100.0 | 7 | 4 | -94, 112 | -90, -22 | -160, -176 | -71, -15 | -125, 5 | -135, 137 | -124, 140 |  |
| H | H4-8-1 | BBAAALBB | RDNSKNT<br>A | 6,269 | 1,953 | 94.0 | 96.8 | 646 | 333 | -143, 149 | -122, 110 | -65, -30 | -67, -34 | -102, 2 | 53, 47 | -129, 141 | -113, 144 |
| H | H4-8-2 | BBAALABB | RDNSKSTA | 60 | 19 | 0.9 | 1.1 | 26 | 12 | -136, 155 | -80, 169 | -60, -39 | -72, -12 | 64, 28 | -104, -11 | -135, 137 | -115, 145 |
| H | noise | - | - | 347 | 37 | 5.1 | 2.1 | 121 | 13 | - | - | - | - | - | - | - | - |

Properties and frequencies of L4 and H4 structural clusters and noise structures for each length of light chain and heavy chain DE loop. The clustering was performed on the entire PDB and on a subset of structures that pass an electron density cutoff ( $EDIA \geq 0.75$ ). $\phi, \psi$ values (in degrees) are given for each residue in each cluster of L4-6, L4-8, H4-6, H4-7, and H4-8 DE loop length families. Ramachandran map regions are: A=alpha-helix region; B = beta sheet region; P = polyproline II helix region; E = epsilon region (lower right of Ramachandran map); L=alpha-left region.
\* Cluster L4-6-2 is composed of 75% $\lambda$ chains and 25% $\kappa$ chains.

Because structures of low resolution or those with dynamic loops may be solved incorrectly, we used the EDIA program (Electron Density for Individual Atoms) to evaluate the electron density of the backbone atoms in each DE loop in the PDB (15). EDIA provides an atom-level assessment of electron density by integrating the 2fo-fc density in a sphere centered on each atomic coordinate. We repeated the clustering on all structures that pass a 0.75 EDIA cutoff for the backbone carbonyl oxygen atoms of the DE loop. The number of structures and unique sequences for each cluster is also provided in Table 2 for the 0.75 EDIA data set, representing structures that are well-solved in the region of the DE loop (See Methods). The average EDIA score for atoms in 2.8 Å structures is 0.8, so most loops with resolution worse than 2.8 Å are removed from the EDIA-filtered data set. The EDIA set is about one third the size of the unfiltered data set but contains about half the number of unique sequences.

For length-6 L4 structures, we observed four different clusters. Figure 3A displays the ϕ,ψ plots for the L4 length-6 clustering, where each row represents a cluster, and each column represents a residue within L4. This figure colors the data points by framework identity (κ in blue, λ in magenta) and shows the noise data as the bottom-most row. The four clusters partition out primarily according to gene, yielding the following clusters (ordered by decreasing size): a primary κ gene cluster (L4-6-1), a mixed λ/κ cluster (L4-6-2) that contains 75% λ chains and 25% κ chains, a secondary κ cluster (L4-6-3), and a secondary λ cluster (L4-6-4). Ten percent of chains end up in noise, which likely includes many low-resolution structures that are not correctly solved. With an EDIA cutoff of 0.75, DBSCAN put only 3.6% of the chains in noise.

**Figure 3.**
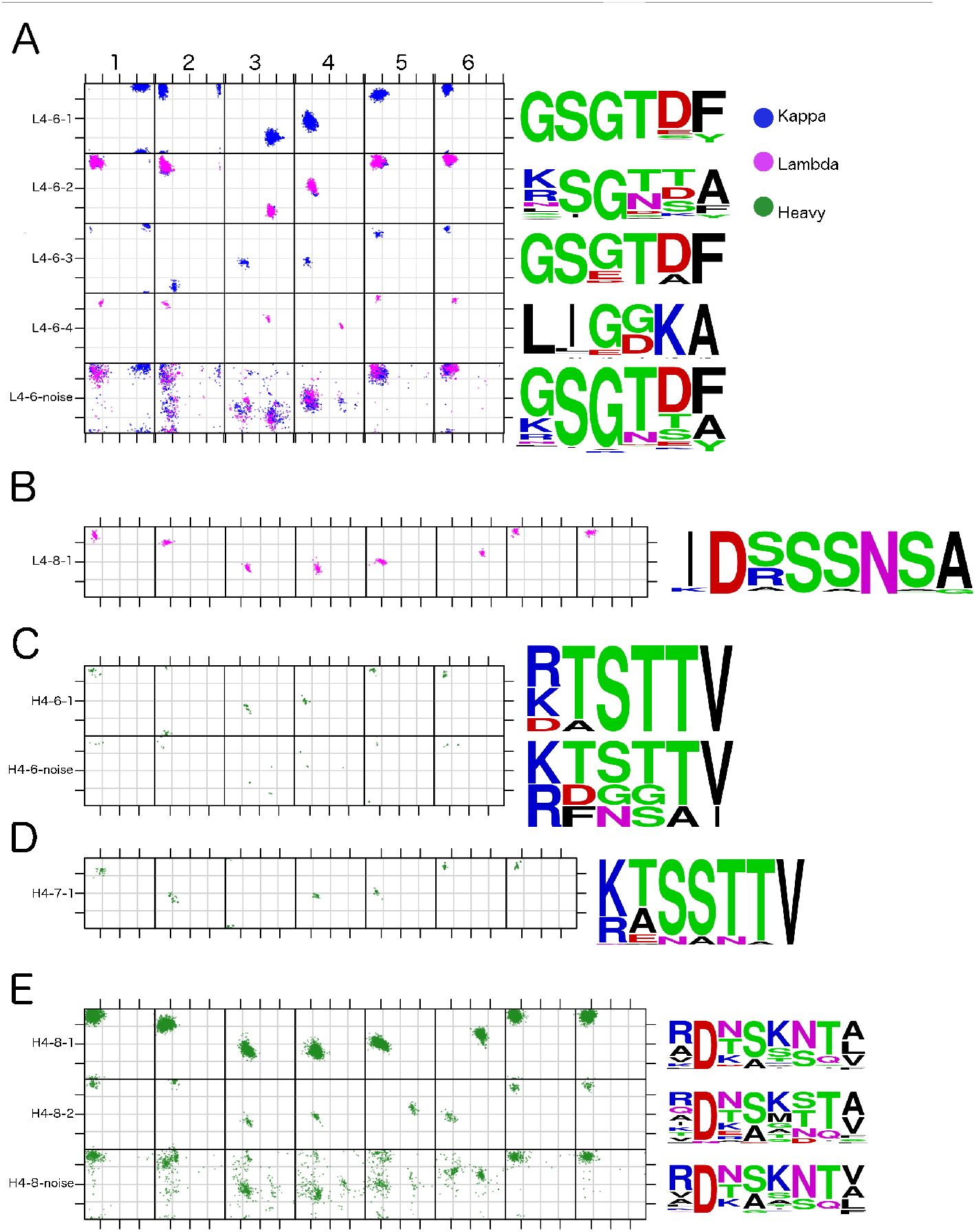
Canonical conformations of L4 and H4. ϕ,ψ plots for each residue in the DE loop for each of the L4 and H4 DE loop clusters from the full data set (no EDIA cutoff) with their respective sequence logos. **A.** L4 length-6 loops. **B.** L4 length-8 loops. **C.** H4 length-6 loops. D. H4 length-7 loops. E. H4 length-8 loops

Across the four L4-6 clusters, the changes in backbone conformation may be viewed structurally as a hinge motion away from the variable domain of the antibody, with L4-6-1 being closest to the domain, followed by L4-6-3 and L4-6-2, while L4-6-4 is the farthest away from the domain. Figure 4A shows representative structures of L4-6-1, L4-6-2, L4-6-3, and L4-6-4 DE loops superposed by alignment to the stems of the DE loop (−3 C-terminal, +3 N-terminal).

**Figure 4:**
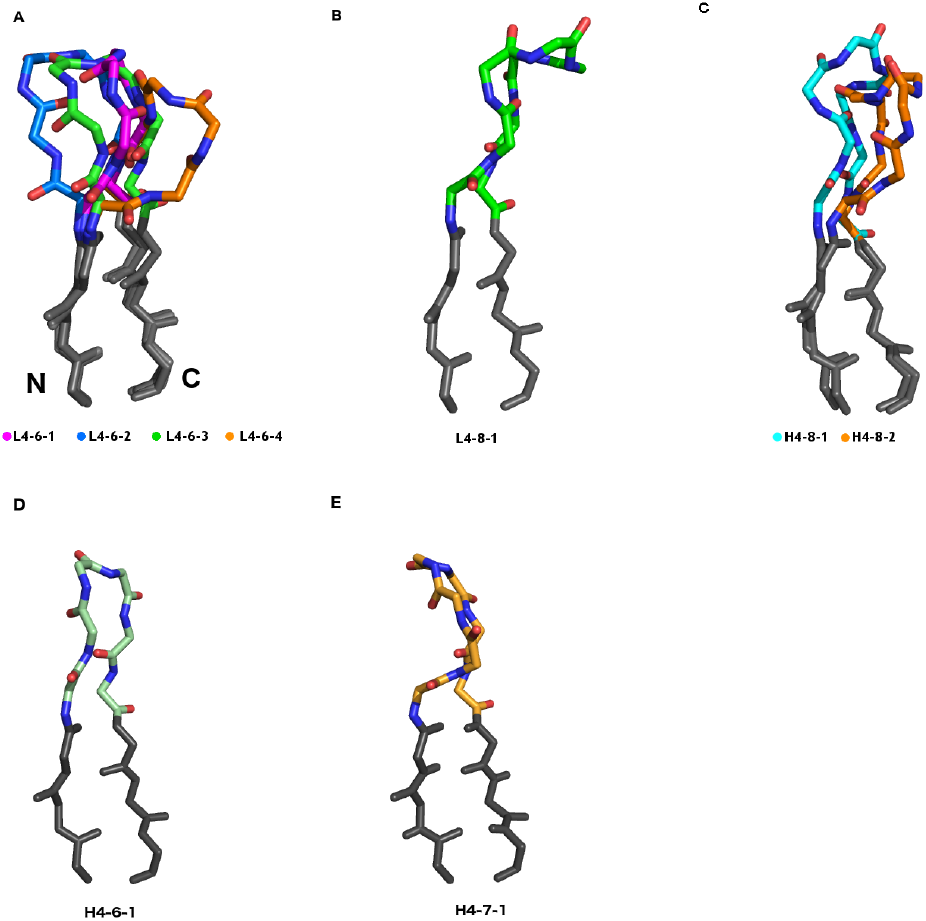
Structures of all DE loop germline-length clusters. **A.** The antibody light-chain DE loop (L4-6) backbone is shown for each cluster where L4-6-1 (PDB 3d9aL, blue) sits closest to the antibody domain, L4-6-3 (1mjuL, green) hinges slightly away and flips the second carbonyl of the DE loop backbone about 180° relative to the other clusters, L4-6-2 (4unuA, magenta) hinges further away from the domain than L4-6-1 and L4-6-3, and L4-6-4 (6frjH, orange) sits the furthest away from the domain. The stems of the DE loop are colored dark gray. **B.** Same representation as in (A), but for the sole L4-8-1 (5jpjA, green) cluster. **C.** Same representation as in (A), but for the H4-8-1 (5e7bA, cyan) and H4-8-2 (6bliJ, orange) clusters..D. Same representation as in (A), but for the H4-6-1 cluster (6i9iH, green). E. Same representation as in (A) but for the H4-7-1 cluster (6dbdD).

The primary difference between the two biggest clusters, L4-6-1 and L4-6-2, is the amino-acid identity and Ramachandran conformation of the first residue. In the germlines of all human κ light chains and nearly all mouse κ light chains, the first residue of the DE loop is glycine. In human and mouse λ germlines, the first residue is (in order of most common to least common): Lys, Ser, Ile, Asp, Leu, Arg, or Thr in 81 out of 84 human and mouse IMGT alleles and Gly in only 3 human alleles (all IGLV9-49, not represented in the PDB). In L4-6-1, the first residue is in an epsilon conformation (ϕ=119.8°, ψ=170.2°), consistent with a Gly in κ antibodies. In L4-6-2, the first residue is in a beta conformation (ϕ=-144.9°, ψ=137.2°), consistent with the λ non-Gly residues. 252 out of 333 κ structures in L4-6-2 (75%) contain somatic mutations at the first residue position from Gly to Arg, Ala, Glu, and Gln in decreasing order of frequency. In the 0.75 EDIA data set clustering, 86% of the κ structures in L4-6-2 have somatic mutations from Gly to some other amino acid at position 1. Several structures in L4-6-2 have germline-encoded hydrophobic residues (Leu, Ile) at the first two residues of the DE loop in the L4-6-2 sequences (e.g., mouse IGLV1*01, IGLV2*01; human IGLV7-43*01, IGLV7-46*01).

L4-6-3 is an all-κ cluster that differs from all-κ L4-6-1 at positions 2 and 3, such that L4-6-1 has average (ϕ_2_,ψ_2_=-164.1°,157.5°; ϕ_3_,ψ_3_=73.6°,-100.0°) and L4-6-3 has average (ϕ_2_,ψ_2_=-90.4°,-137.0°; ϕ_3_,ψ_3_=-93.6°,-26.0°). In the L4-6-3 structures, 18 of 69 (26%) chains (from 3 PDB entries) have somatic mutations at position 3 from Gly to Glu, which accounts for the change in residue 3 from an epsilon conformation to an alpha conformation (with a compensating change at residue 2). The remaining structures have germline sequences, including one structure with a germline Arg residue at residue 3.

L4-6-4 is an all-λ cluster that differs from the L4-6-2 λ/κ cluster at residue positions 3 and 4. This cluster has residues with the left-handed conformation at positions 3 and 4 (ϕ_3_,ψ_3_=51°,48°; ϕ_4_,ψ_4_=70°,12°), which is not seen in any of the other clusters. Consistent with these conformations, a majority of the loops have glycine residues at these positions. Similar to some of the sequences in L4-6-2, the sequences that are part of this cluster start with hydrophobic residues at positions 1 and 2 (Leu or Ile) that come from human IGLV7-43, llama IGLV8-3, and mouse IGVL1 and IGLV2 germlines. Out of the 31 structures of L4-6-4, 12 structures have the germline sequence. The remaining sequences are somatic mutations of the third position (G→E, 4 structures), the second position (L→R, L→I, 2 structures), and the fourth position (D→G, 13 structures).

We evaluated both L4-6-3 and L4-6-4 by their electron density by examining how many of their chains and unique sequences are produced from the clustering of the EDIA≥0.75 data set. For L4-6-3, four out of seven unique sequences remain, and for L4-6-4, five out of six unique sequences remain after the EDIA cutoff, indicating that they are robust clusters, and not related to mis-solved residues.

For the 67 length-8 L4 structures in the PDB, which are related to a small number of λ germlines in humans, mice, rats, rabbits, and macaques, we observed a single cluster (Figures 3B and 4B) representing 7 unique sequences. No structures were placed into noise by the DBSCAN algorithm, indicating a low level of structural variance. Out of 49 PDB entries containing light chains with length-8 DE loops, 17 of them are involved in Bence-Jones homodimers associated with light-chain amyloidosis (54). After enforcing the EDIA cutoff, 41 of the 67 chains remain, which contain all 6 of the unique sequences found in the clustering with no EDIA cutoff.

For canonical length-8 H4 structures, clustering with DBSCAN produced one large cluster with 6,269 chains, and several small clusters with less than 60 chains each. The sequences of the small clusters were very similar to those in the large cluster. Many of them involve peptide flips from the large cluster and may be incorrectly solved. Peptide flips involve changing the ψ dihedral of one residue by 180° and the ϕ dihedral of the next residue by 180° (55). The carbonyl residue of the first residue moves by more than 3 Å in a peptide flip, so the electron density of the oxygen atom of each residue in a loop is diagnostic of mis-solved peptide flips, which are common in protein loops. By clustering the EDIA≥0.75 data set, in addition to the large cluster, only one small cluster remained with a substantial number of structures (19), as well as unique sequences (12). The others involved peptide flips from the large cluster, and likely are due to incorrectly solved structures.

We therefore chose to name only two H4-8 clusters, H4-8-1 and H4-8-2 (Figure 3E and 4C). The H4-8-1 cluster has 646 unique sequences, exhibiting far greater sequence variation than any of the L4-6 clusters. For H4-8-2 (ϕ_2_,ψ_2_=-80°,-160°) there is a small shift in residue 2 compared to H4-8-1 (ϕ_2_,ψ_2_=-122°,-110°), and a substantial shift at residues 5 and 6, which change the conformation of these residues to LA (ϕ_5_,ψ_5_=64°,28°; ϕ_6_,ψ_6_=-104°,-11°), from AL (ϕ_5_,ψ_5_=-102°,2°; ϕ_6_,ψ_6_=-53°,47°) respectively. Any other clusters generated in the clustering step had fewer than 10 unique sequences, or they disappeared after enforcing the EDIA cutoff, leading to a suspicion that they are clusters consisting of mis-solved residues at position 5 and 6.

The conformation of length-8 H4 structures is the same conformation as length-8 L4 structures as shown in structural alignment of the two clusters by the CDR4 stem regions (Figure 5A). We also clustered a small number of H4 loops of lengths 6 (from the rabbit IGHVIS69 germline) and 7 (from rabbit IGHV1S45, IGHV1S47, and IGHV1S69 and llama IGHV1S3 germlines) (Table 2, Figure 3C and 3D), which produced one cluster for each length, designated H4-6-1 and H4-7-1. The H4-6-1 cluster is similar in conformation to L4-6-3 as shown in Figure 5B.

**Figure 5.**
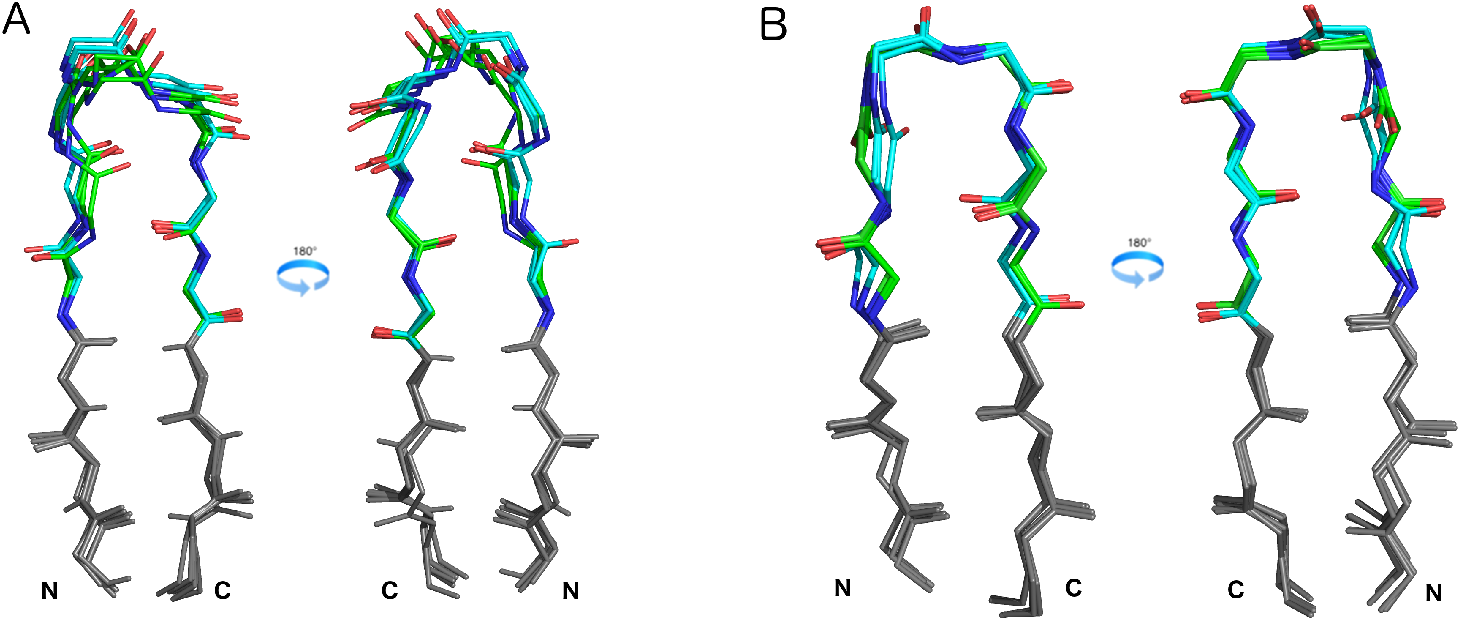
Comparison of H4-8 and L4-8 clusters with structural homology. **A.** Superposition of several high-resolution heavy chain H4-8-1 structures (cyan, PDB chains: 2×1qA, 4qyoB, 2vxvH), and L4-8-1 structures (green, PDB chains: 1cd0A, 2w01A, 3h0tA) aligned by the stem of CDR4 (colored in gray) show structural similarity between the two clusters. **B.** Same superposition as in 5A, but for L4-6-3 (green, 1mujL, 6qnkC, 6mv5L) and H4-6-1 (cyan, 6i8iH, 6cf2A, 6banH) clusters.

### Interactions between CDR4 and CDRs 1 and 2

To describe the relationship between various L4 conformations with CDR1 and CDR2 conformations, we first calculated the occurrence of each L4-6 cluster given the various common L1 clusters (Table 3). The κ L1 clusters have more than 80% L4-6-1 DE loops. For most of these the secondary cluster is L4-6-2, indicating a tendency for residue 1 in the corresponding germlines to mutate from Gly to another residue type. The L1-16-1 structures prefer L4-6-3 as a secondary cluster, probably because germlines with length 16 more frequently contain residues that are not Gly at position 3 of L4 (11 of 63 human, mouse, and rat germlines in IMGT).

**Table 3.** Occupancy of the co-occurrence of L1/L4 pairs from structures in the PDB.

| L1 cluster | Gene | # chains | L4-6-1 | L4-6-2 | L4-6-3 | L4-6-4 | L4-8-1 |
| --- | --- | --- | --- | --- | --- | --- | --- |
| L1-10-1 | κ | 185 | 99.4 | 0.6 | - | - | - |
| L1-10-2 | κ | 93 | 100.0 | - | - | - | - |
| L1-11-1 | κ | 1559 | 90.0 | 9.9 | 0.1 | - | - |
| L1-11-2 | κ | 508 | 91.6 | 8.2 | 0.2 | - | - |
| L1-12-1 | κ | 175 | 98.1 | 1.9 | - | - | - |
| L1-12-2 | κ | 110 | 98.1 | 1.9 | - | - | - |
| L1-15-1 | κ | 274 | 98.8 | - | 1.2 | - | - |
| L1-16-1 | κ | 597 | 84.1 | 1.5 | 14.4 | - | - |
| L1-17-1 | κ | 277 | 99.0 | 0.3 | 0.7 | - | - |
| L1-11-3 | λ | 149 | - | 100.0 | - | - | - |
| L1-12-3 | λ | 32 | - | 100.0 | - | - | - |
| L1-13-1 | λ | 248 | - | 100.0 | - | - | - |
| L1-13-2 | λ | 71 | - | - | - | - | 100.0 |
| L1-14-1 | λ | 193 | - | 85.0 | - | 15.0 | - |
| L1-14-2 | λ | 176 | - | 100.0 | - | - | - |
| H1 cluster | Gene | # chains | H4-8-1 | H4-8-2 | H4-6-1 | H4-7-1 |  |
| H1-13-1 | H | 4285 | 99.1 | 0.1 | 0.5 | 0.3 | - |
| H1-13-2 | H | 64 | 96.0 | 4.0 | - | - | - |
| H1-13-3 | H | 101 | 93.0 | 7.0 | - | - | - |
| H1-13-4 | H | 183 | 99.4 | 0.6 | - | - | - |
| H1-13-5 | H | 86 | 93.8 | 1.2 | - | 5.0 | - |
| H1-13-6 | H | 28 | 100.0 | - | - | - | - |
| H1-13-7 | H | 58 | 94.6 | 5.4 | - | - | - |
| H1-13-10 | H | 16 | 81.3 | 18.7 | - | - | - |
| H1-14-1 | H | 102 | 100.0 | - | - | - | - |
| H1-15-1 | H | 173 | 97.1 | 2.9 | - | - | - |

For each L1 or H1 cluster, the distribution among the L4 or H4 clusters is provided in percent (excluding the noise cluster)

For the λ germlines, all of the L1 clusters except L1-14-1 are 100% L4-6-2. L1-14-1 structures contain 85% L4-6-2 and 15% L4-6-4 DE loops. The L4-6-4 sequence logo (Figure 3A) shows that this cluster comprises sequences that begin with aliphatic amino acids at positions 1 and 2 (Leu or Ile) and have either Lys or Arg at position 5. All germlines with these sequence features have L1 CDRs of length 14 (human IGLV7-43*01, IGLV7-46*01/02, IGLV8-61*01/02, and IGLV8-OR8-1; mouse IGLV1*01/02 and IGLV2*01; rat IGLV1S1*01). The charged residue at position 5 does not take part in a hydrogen bond in any of the structures where the Lys or Arg is present with L1-14-1.

Second, we have calculated all hydrogen bonds between the DE loop and CDR1 or CDR2. Supplementary Table S1 shows all hydrogen bonds calculated between the DE loop and CDR1 or CDR2 for each CDR1 and CDR2 cluster of L1, H1, and H2 (there are no characteristic hydrogen bonds between L4 and L2 with an occupancy over 60%). In the case of hydrogen bonds involving side-chain atoms, the hydrogen bonds are grouped by structures with the same amino acid at the same position within the DE loop.

Hydrogen bonds between the DE loop and CDR1 and CDR2 partition into the following broad categories: (1) backbone-backbone hydrogen bonds shared across several CDR1 clusters (Figure 6C); (2) backbone-backbone hydrogen bonds unique to specific DE loop/CDR1 cluster pairings (Figure 6D, 6E); (3) hydrogen bonds between DE-loop side-chain atoms and backbone atoms at positions shared across several CDR1/CDR2 clusters (Figure 6F); (4) hydrogen bonds between DE loop side-chain atoms and CDR backbone atoms that are specific for some CDR1/2 clusters and lengths (Figure 6A, 6B, 6G); and (5) hydrogen bonds between DE loop backbone atoms and CDR1 side-chain atoms that occur in L1 loops longer than 14 residues (Figure 6H, 6I).

**Figure 6:**
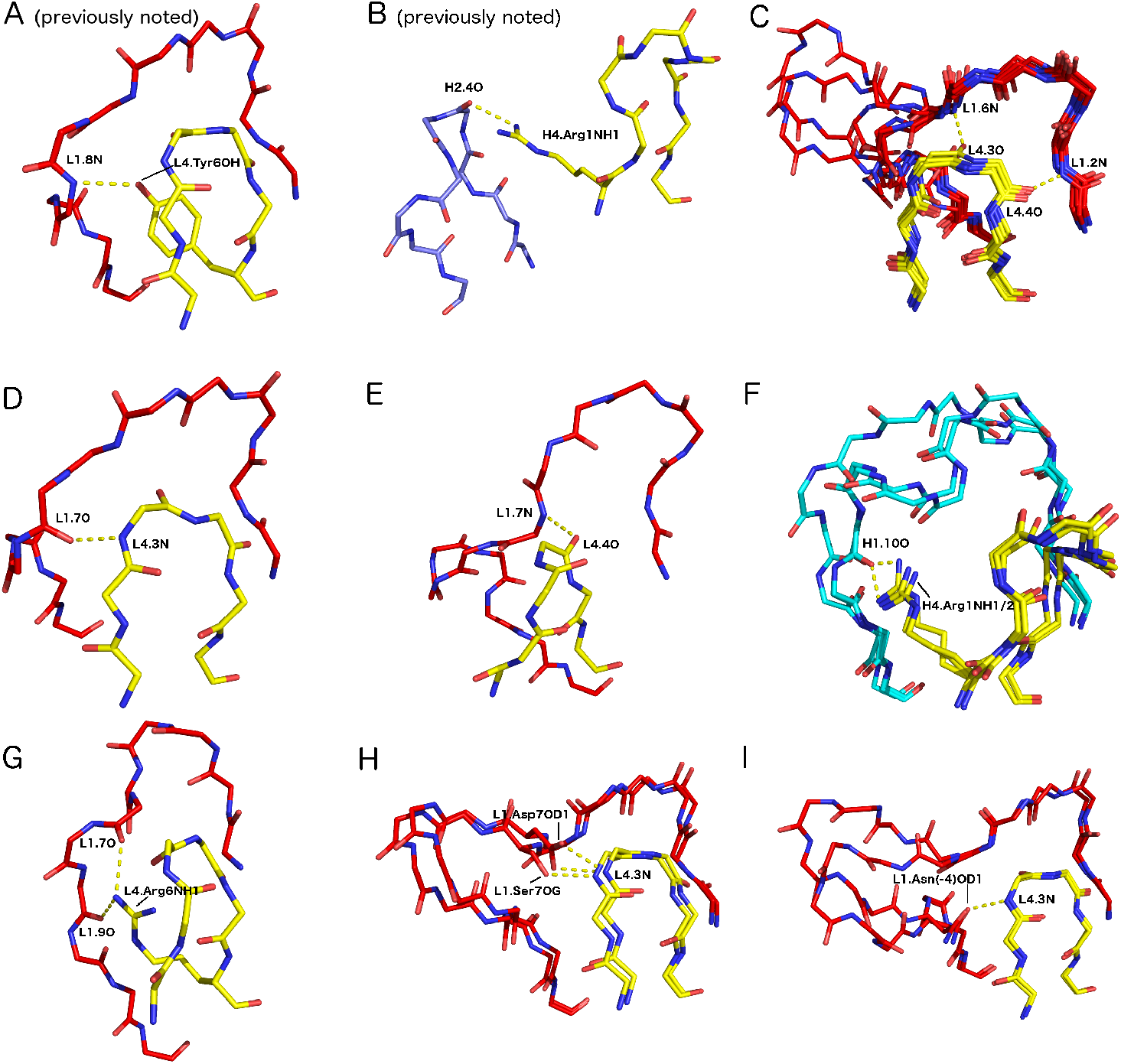
Various characteristic hydrogen bonds between the DE loop and CDR1. Hydrogen bonds are labeled CDR4.resnumAtom/CDRn-cluster.resnumAtom (e.g. H4.6O/H1.2N). If a hydrogen bond is specific to a particular cluster, that is included in the nomenclature. **A.** Side-chain/backbone hydrogen bond L4.Tyr6OH/L1.8N (yellow/red) common in L1-11-2, **B**. Side-chain/backbone hydrogen bond H4.Arg1NH1/H2.3O (yellow/purple) common in H2-10-1. **C.** Side-chain/backbone hydrogen bonds L4.3O/L1.6N and L4.4O/L1.2N (both yellow/red) common in L1-10, L1-11, L1-12, L1-13-1, L1-14-2, L1-15-1, L1-16-1, L1-17-1 clusters. **D.** Backbone/backbone hydrogen bond L4.3N/L1-11-1.7O common in L1-11-1. **E.** Backbone/backbone hydrogen bond L4-6-2.4O/L1-14-1.7N common in L1-14-1. **F**, Side-chain/backbone hydrogen bond H4.R1NH/H1-13.10O common in H1-13-1,2,3,4. **G.** Side-chain/backbone hydrogen bond L4.R6NH/L1-12-3.7O and L4.R6NH/L1-12-3.9O common in L1-12-3. **H**. backbone/side-chain hydrogen bonds L4.3N/L1.S7OG and L4.3N/L1.D7OD common in L1-15-1. **I**. L4.3N/L1.N(−4)OD common in L1-16-1 and L1-17-1.

Regardless of DE loop conformation, DE residue 4 in length-6 L4 forms a backbone-backbone hydrogen bond to the backbone nitrogen of the second residue in CDR1 (counting L1 residues immediately after the Cys of the disulfide bond; Figure 6C) for the vast majority of L1 clusters (L1-14-1 excluded). This hydrogen bond is part of the beta sheet containing the C-terminal segment of CDR4 and the N-terminal strand at the beginning of CDR1. On the light chain, most DE loop structures also have a backbone-backbone hydrogen bond between DE residue 3 and residue 6 of CDR1 (L1-12-3, L1-13-1, L1-14-1, and L1-14-2 excluded; Figure 6C). Structures that have both of these hydrogen bonds have very similar conformations between the residues that are hydrogen bonded, even amongst a diverse set of L1 lengths.

Beyond backbone-backbone hydrogen bonds correlated with the arrangement of the L4 backbone atoms, we note several particular side-chain/backbone hydrogen bonds that occur uniquely with L1 conformations. For example, residue Arg1 hydrogen bonds to the backbone carbonyl of residue 7 of L1-13-1 in 24/26 chains, whereas when the DE residue 1 is Lys, but is still paired with L1-13-1, the hydrogen bond occupancy is only 24% (213 cases).

As noted in previous studies (5,56), the OH atom of the Tyr6 side chain of the L4 loop forms a hydrogen bond to the backbone nitrogen atom of residue 8 in length-11 L1 CDRs (Figure 6A), flipping its conformation from L1-11-1 (predominantly Phe6) to L1-11-2 (predominantly Tyr6). This hydrogen bond forms in 85% of structures of L1-11-2 with a Tyr residue at position 6 of L4. When this residue is Phe6 instead, this hydrogen bond is lost, and the structure of L1 is L1-11-1, and a hydrogen bond instead forms between the backbone nitrogen of DE residue 3 to the carbonyl of L1-11 residue 7 (Figure 6D). In similar fashion, we note a new hydrogen bond of the side chain of Arg6 in L4-6-2 to the carbonyl backbone oxygen atoms of residues 7 and 9 of L1-12-3 structures, creating a hydrogen bond network. This is an example where the exclusive occurrence of an L1/L4 pair is associated with a unique contact between L1 and L4.

For L1-15-1 and L1-17-1, the carbonyl oxygen of DE residue 1 of L4 is not only hydrogen bonded to a backbone nitrogen atom in L1, but the backbone nitrogen atom of DE residue 1 is also hydrogen bonded to various side-chain oxygen atoms of residue 7 in L1-15-1 (Asp, Ser, or Thr; Figure 6H), or residue 14 (Ser or Asn; Figure 6I) in L1-17-1. Taken together, these results demonstrate that L1/L4 pairs often entail highly specific interactions, facilitated by the L4 cluster-specific arrangement of the L4 backbone and side-chain atoms, which can provide stabilizing hydrogen bonds between L4 and L1.

For H4, in addition to the conserved hydrogen bond involving residue 4 in most H1/H4 pairs for common H1 lengths and clusters (similar to the L4 residue 4 hydrogen bond in Figure 6C), there are several side-chain/backbone hydrogen bonds that are shared between several H1 clusters and various residues in H4. Most notably, the Arg1 residue in H4 hydrogen bonds with the backbone carbonyl of residue 10 in the large H1-13-1 cluster using both the NH1 and NH2 atom in the interaction (Figure 6F). For specific hydrogen bonds, the occupancy of the hydrogen bond depends highly on the H4 residue type. DE residue Asn6 uses its side-chain oxygen atom as well as its side-chain NH2 group to form side-chain/backbone hydrogen bonds between residue 2, residue 5, and residue 7 of H1, stabilizing the H1-14-1 conformation with a hydrogen bond network.

The common Arg1 side-chain of H4 often forms hydrogen bonds with H1 and H2 (Figure 6B). The most common hydrogen bond of Arg 1 is to residue 10 of over 1400 H1-13 structures (in clusters H1-13-1, 2, 3, and 4). In our clustering of the CDR loop conformations, we noted the common presence of H4-Arg1 in H2-10-2 structures and its relative absence in H2-10-1 structures (56), implicating a hydrogen bond of Arg1 with the backbone carbonyl of residue 3 of H2-10-2. While the Arg1 side chain is in contact with this carbonyl atom in many structures, the hydrogen bond geometry is poor and Rosetta does not identify it as a hydrogen bond. Its interaction with H2-10-2 is primarily hydrophobic, as noted by Tramontano et al (6).

For H2-10-1, most H4 loops have an Ala residue at position 1. However, when position 1 in H4 is Arg, 70% of structures have a hydrogen bond from the NH1/NH2/NE atom of Arg to the backbone carbonyl of the residue at position 4 in H2-10 (354 cases with Arg1 when the H2 conformation is H2-10-1). When this residue is Gln, 59% of structures (15/27 chains with H2-10-1 and H4-8-1 with Gln at position 1) contain a hydrogen bond between the NE2 atom of Gln and the backbone carbonyl oxygen of residue 4 of H2. When this residue is instead Lys1, the occupancy of this hydrogen bond is only 42% (17/40 chains with H2-10-1 and H4-8-1 with Lys at position 1). In all other H2-10-1 structures, there is no polar residue at position 1 with a sidechain available to make a hydrogen bond to H2-10. This indicates that a hydrogen bond from position 1 in H4 is not a strong association to the presence of the H2-10-1 conformation, but structures that have this hydrogen bond may have better stability, especially when that hydrogen bond is made from Arg1.

When the conformation of H2-10 is H2-10-2, the Arg1 of H4 rarely forms hydrogen bonds with the carbonyl oxygen at position 4 of H2-10-2 (less than 3% in all 1,180 structures with Arg1 and H2-10-2). Instead, the ND2 atom of Asn3 of H4, which is fully conserved in structures with H2-10-2, forms hydrogen bonds with the same backbone carbonyl oxygen at position 4 of H2 (73% of 1180 structures with Asn at position 3 in H4 and H2-10-2). This points to Asn3 in H4 as a major indicator of the H2-10-2 conformation.

### Analysis of the sequence variability in DE loops arising from somatically mutated and germline sequences

From a set of ~2.5 million sequences of naïve human antibodies (57–62), we calculated the sequence entropy in four of the most prevalent human germlines for the heavy, κ, and λ genes in the data set (Figure 7A), as well as the entropy of human germline sequences of the same length (Figure 7B). As other studies have noted (63), variability of both framework and CDR residues depends highly upon germline. We did not find any DE loops with insertions in this set, so we did not have to account for insertions in the calculation of the sequence entropy.

**Figure 7:**
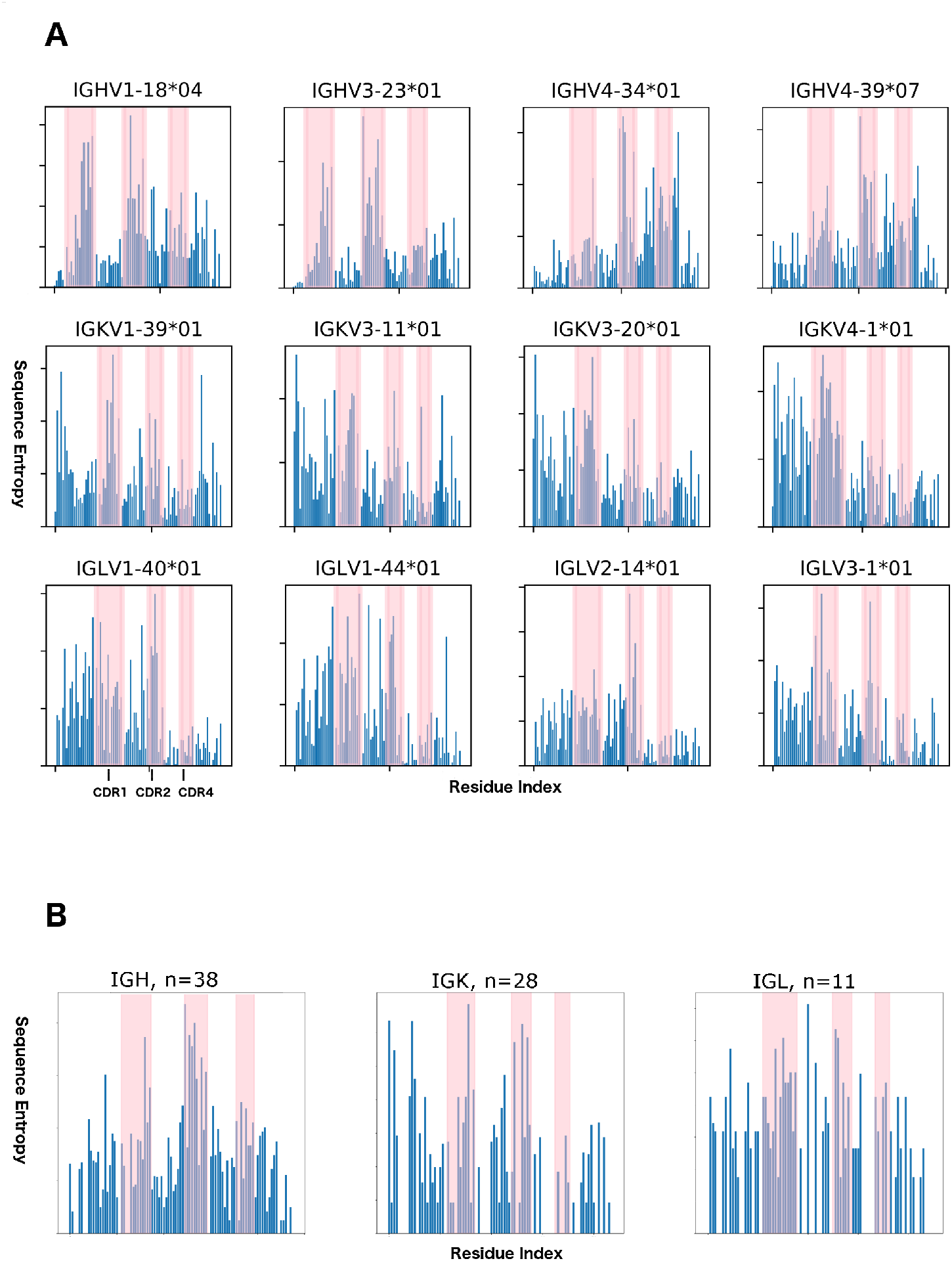
Sequence entropy in naïve human antibodies and human germlines. **A.** Sequence entropy for 12 common germlines in a naive human antibody sample (>10,000 sequences for each germline). **B.** Sequence entropy for human germlines derived from all IGKV, IGLV, and IGHV sequences from IMGT. From left to right, the pink shaded regions indicate CDR1, CDR2, and the DE loop. CDR3 is omitted due to varying lengths and different diversification mechanisms.

For H4 sequence variability, we find that in cases of somatic mutation of any one particular germline, the average sequence entropy of H4 for each of the four germlines exceeds the average sequence entropy for FR1, FR2, and FR3 of the same germline antibodies (Table 4). However, these DE loop residues are less variable than H1 or H2 residues within the same germlines.

**Table 4.** Average sequence entropies for CDR and framework regions.

| **germline** | **CDR1** | **CDR2** | **FR1** | **FR2** | **FR3** | **CDR4** |
| --- | --- | --- | --- | --- | --- | --- |
| IGHV1-18*04 | 0.38 | 0.44 | 0.04 | 0.17 | 0.21 | **0.27** |
| IGHV3-23*01 | 0.42 | 0.74 | 0.03 | 0.15 | 0.21 | **0.26** |
| IGHV4-34*01 | 0.14 | 0.38 | 0.05 | 0.14 | 0.17 | **0.29** |
| IGHV4-39*07 | 0.21 | 0.36 | 0.08 | 0.13 | 0.15 | **0.20** |
| IGKV1-39*01 | 0.29 | 0.22 | 0.20 | 0.12 | 0.13 | 0.11 |
| IGKV3-11*01 | 0.25 | 0.22 | 0.21 | 0.10 | 0.10 | 0.11 |
| IGKV3-20*01 | 0.30 | 0.21 | 0.23 | 0.10 | 0.10 | 0.07 |
| IGKV4-1*01 | 0.30 | 0.15 | 0.24 | 0.09 | 0.08 | 0.07 |
| IGLV1-40*01 | 0.25 | 0.29 | 0.22 | 0.12 | 0.05 | 0.07 |
| IGLV1-44*01 | 0.27 | 0.26 | 0.21 | 0.12 | 0.06 | 0.07 |
| IGLV2-14*01 | 0.24 | 0.30 | 0.17 | 0.13 | 0.06 | 0.08 |
| IGLV3-1*01 | 0.28 | 0.26 | 0.20 | 0.10 | 0.06 | 0.13 |
| **gene** | **CDR1** | **CDR2** | **FR1** | **FR2** | **FR3** | **CDR4** |
| IGH | 0.78 | 1.50 | 0.53 | 0.61 | 0.53 | **0.86** |
| IGK | 0.57 | 0.71 | 0.46 | 0.24 | 0.20 | 0.19 |
| IGL | 0.80 | 0.63 | 0.43 | 0.33 | 0.28 | **0.52** |

Average sequence entropies partitioned by CDR or framework region, excluding CDR3. Bolded values are those where the CDR4 average sequence entropy either compares to CDR1/CDR2, or exceeds the values for FR1, FR2, and FR3

For κ antibodies, residue 2 has a higher sequence entropy than about 80% of the other framework residues (Figure 7A). Comparing 28 germline sequences for human κ antibodies (Figure 7B), we observe three highly variable residues (DE residue 2, 5, and 6), and 3 completely conserved residues (IMGT residues Gly1, Gly3 and Thr4). In the variable residues of L4, the entropy is comparable to the most variable framework residues in germlines, the average entropy does not compare to the average entropy of L1 or L2, and does not exceed the average entropy of FR1, FR2, or FR3 (Table 4).

Within each λ germline, L4 sequences are much less somatically mutated than in κ antibody sequences (Figure 7A). The amount of sequence variability due to somatic mutation is less than even the most variable framework residues, and does not compare to sequence variability in L1 or L2. However, looking at 11 germline sequences, average sequence entropy in λ L4 sequences exceeds that of FR1, FR2, and FR3, but is less than that of L1 and L2 (Table 4). Sequence variability at positions 1, 4, and 5 is comparable to both L1 and L2 (Figure 7B). This indicates that sequence variability in λ L4 relates primarily to germline sequence differences, and not somatic mutation.

### Non-canonical L4 and H4 length in HIV-1 bNAbs

All known mammalian VH germlines have a DE-loop of 8 residues, except for a small number of rabbit VH genes with DE-loop lengths 6 and 7, some of which are represented in the PDB (Table 2). All known mammalian VL germlines have a DE-loop of either 6 or 8 residues, except for one alpaca germline (IGLV5-12*01) with a DE loop of length 3 (not represented in the PDB). There are 119 chains from 43 entries in the PDB with H4 loops longer than 8 amino acids, ranging from 10 amino acids to 16 amino acids. There are 65 chains from 50 entries that have insertions in L4 (all lambda chains), resulting in L4 loops of length 9. All of the antibodies in the PDB with non-germline insertions in L4 or H4, except one, are broadly neutralizing antibodies against HIV-1. The sole exception is an engineered nanobody against Higb2 toxin (PDB: 5MJE). Table 5 lists the various PDB structures of the bnAbs that have insertions in either the light or heavy chain DE loop as well as their sequences, germlines, and which bnAb class they belong to. The structures are shown in Figure 8 and grouped by antibody class. In these structures, L4 and H4 bind the antigen epitope better than H1 and H3 as well as any of the light chain CDRs, as demonstrated by the extent of antigen-buried surface area for each CDR including CDR4 (Figure 9).

**Table 5.**
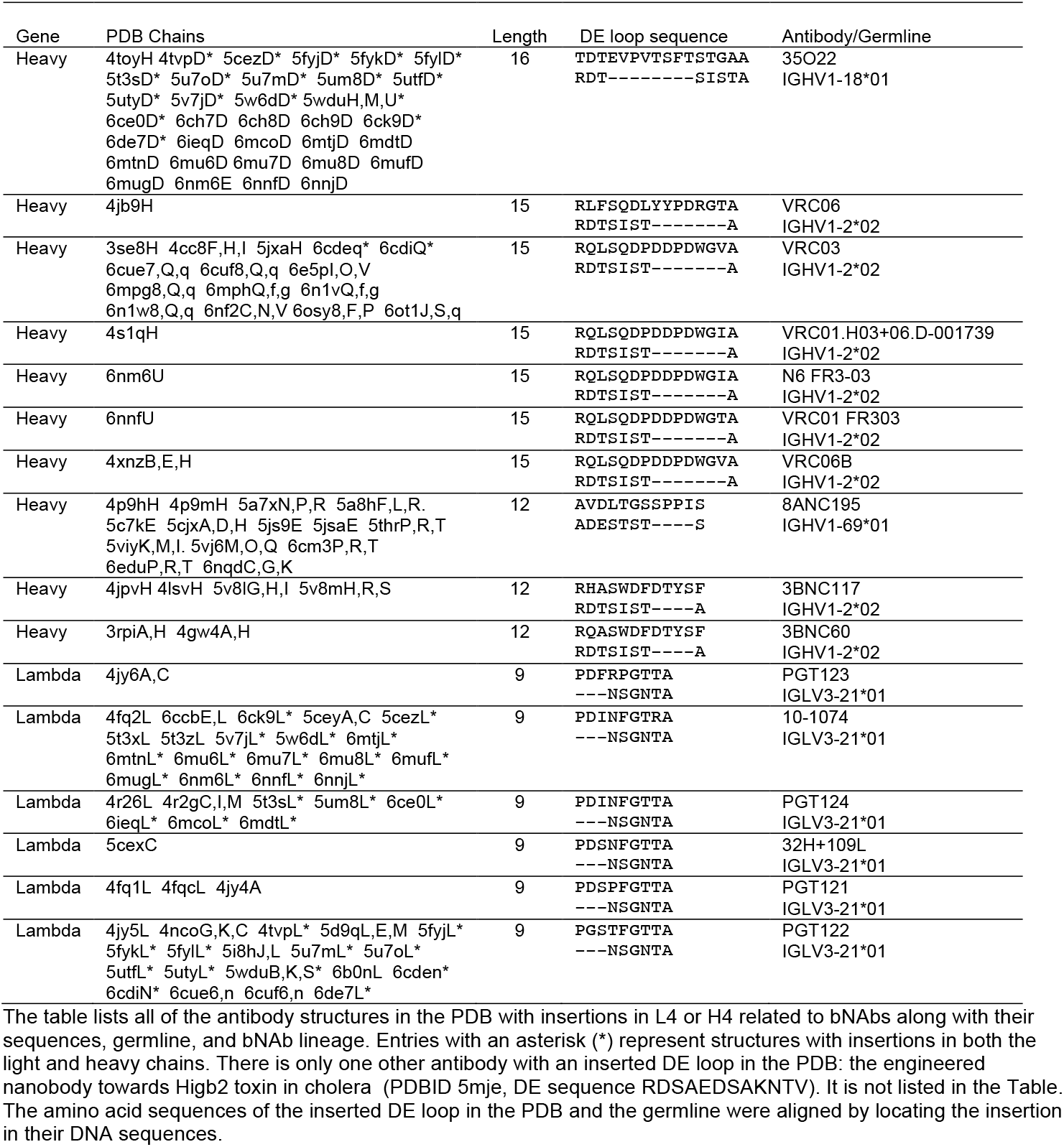
HIV-1 bNAbs with insertions in L4 and/or H4.

**Figure 8.**
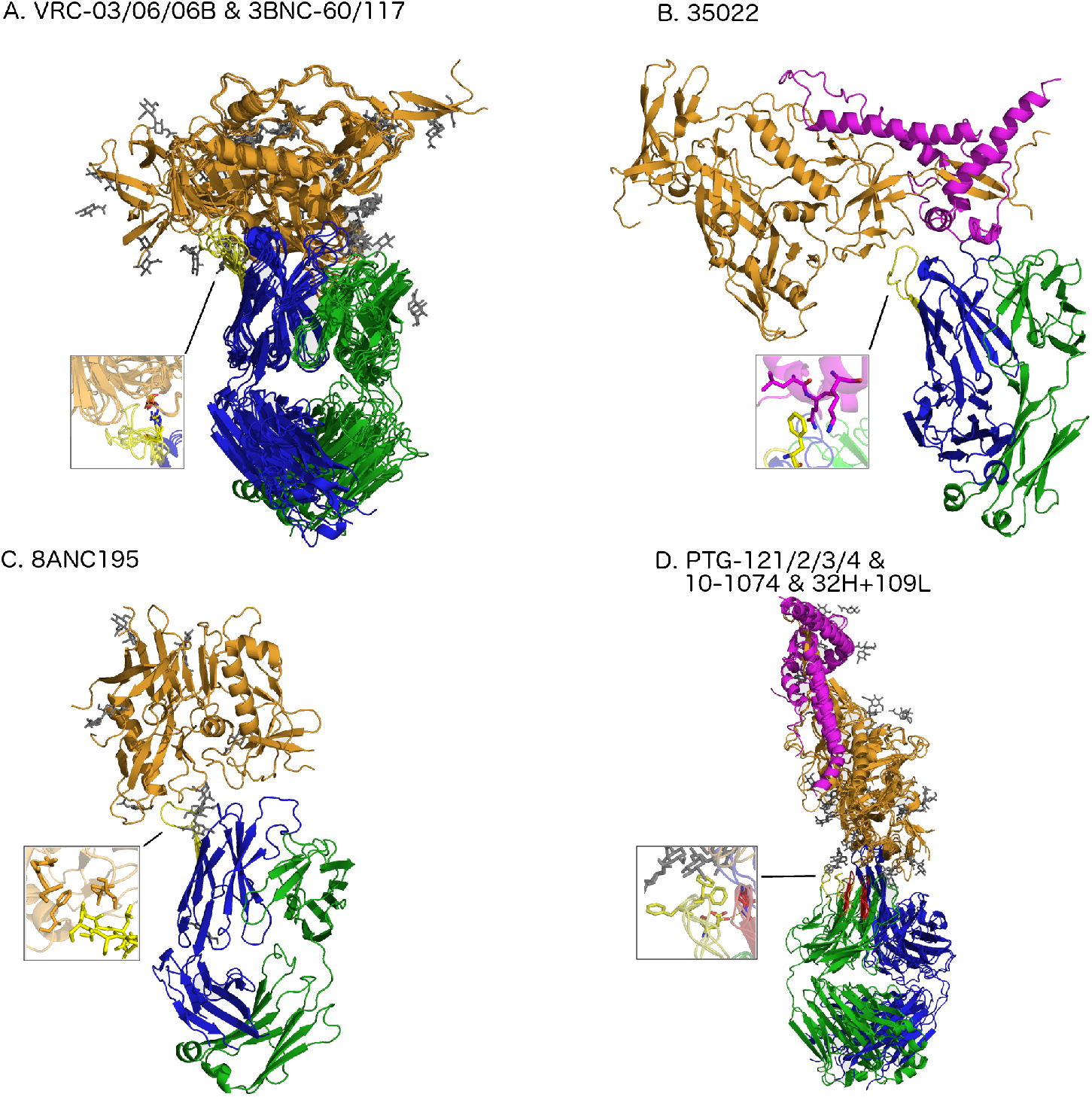
Alignment of a subset of gp120 binding HIV-1 bNAbs representing all unique DE loop sequences. **A.** Aligned structures of H4 inserted bNAbs of the VRC and 3BNC series (one structure per H4 sequence). The inset shows hydrophobic contacts between the antibody-antigen interface, as well as a salt bridge between the first Arg residue in H4 and the antigen. **B**. Structure of the 35O22 antibody found in 35 PDB entries with an H4 of length 16. **C**. Structure of the 8ANC195 antibody with an H4 of length 12 represented in 16 PDB entries. **D.** Aligned structures of L4-inserted bnAbs binding to HIV-1 gp120 (one representative per unique L4 sequence of length 9). The inset shows hydrophobic contacts between the antibody-antigen interface, as well as hydrogen bonds at the antibody-antigen interface and between L1 and L4, stabilizing a unique L1 conformation.

**Figure 9.**
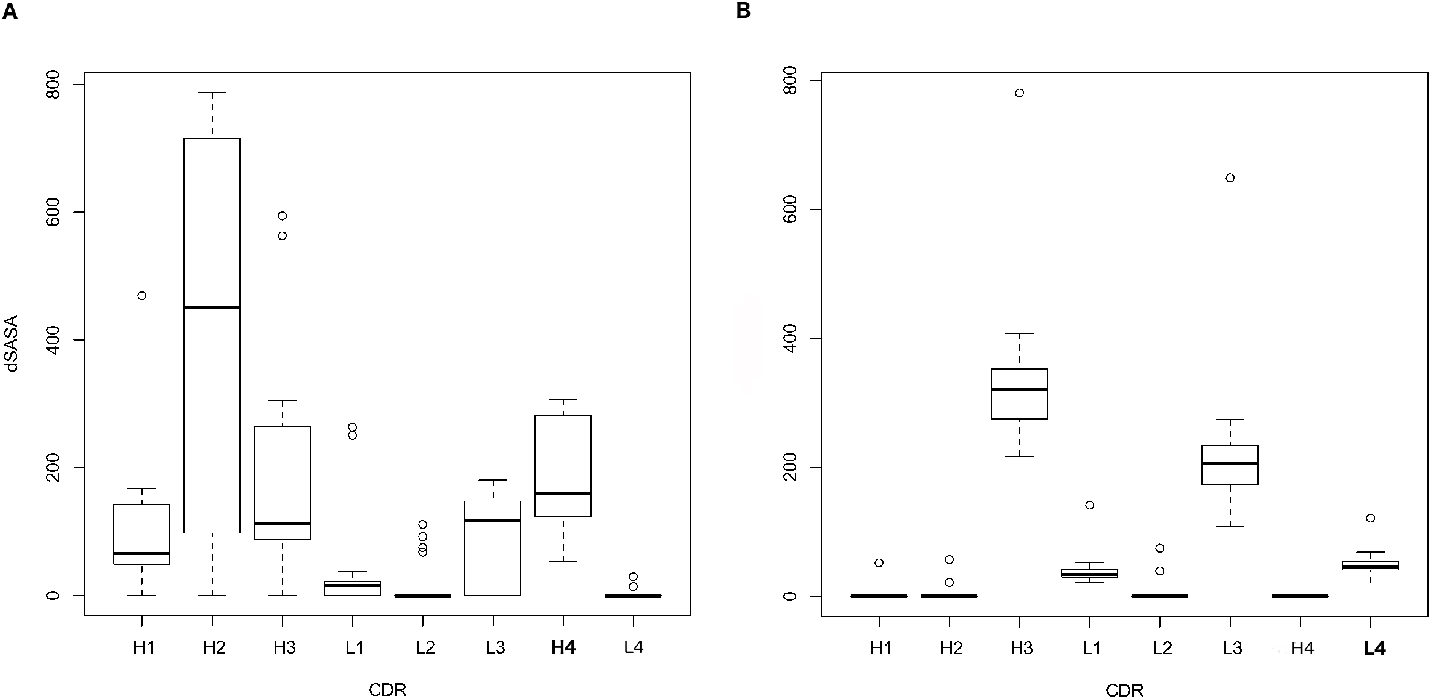
Buried surface area for each CDR at the antibody-antigen interface of HIV-1 bNAbs that bind to gp120 in the PDB. **A.** Buried surface area plot for 18 PDB structures (non-redundant by chain) with insertions in H4. **B**. Buried surface area plot for 31 PDB structures (non-redundant by chain) with insertions in L4.

In the case of elongated H4 loops that contact gp120, the antibodies containing insertions in H4 bind to three separate binding sites. Antibodies in the VRC03/06/06B and 3BNC60/117 classes that have an Arg at position 1 of H4 that makes a salt bridge with residue Asp368 of gp120 near the CD4 binding site (Figure 8A), as noted in previous studies of the VRC01 class of antibodies (16,20). Regardless of insertion length (4 residue insertion in the 3BNC antibodies, and 7 residue insertion in the VRC series of antibodies), these structures all localize to the same epitope on gp120 and share the same salt bridge as noted previously. In this way, DE residue 1 determines localization of the binding site of the VRC class of antibodies, which mimic the CD4 binding to gp120. Besides this interaction, much of the interaction between elongated H4 in the VRC antibodies and gp120 consists of buried hydrophobic contacts (Figure 8A). Figures 8B and 8C show the other structures of bnAbs with H4 loops with somatic insertions.

The non-canonical length L4 loops are related to the Hu_IGLV3_21*01 germline and feature length 9 L4 loops. These loops not only directly bind antigen with hydrophobic interactions at the apex of the loop (Figure 8D), but also stabilize the conformation of L1 through a couple of backbone/backbone and backbone/side-chain hydrogen bonds to a serine in L1, which is sandwiched between L4 and L3. This ‘L1 sandwich’ motif appears to rigidify the binding conformation of the antibody light chain that buries a tremendous amount of binding surface area while binding to gp120 even in the presence of highly glycosylated elements (Figure 8D). The L1-14 conformations associated with these antibodies are exclusive, and no other antibody structures contain these unique conformations of L1. The extended L4 loop length and conformation may stabilize the unusual conformation of L1, and enable the formation of new interactions between the antibody and antigen.

### Features of DE loop sequences from HIV-1 infected individuals

As noted earlier, we did not find any DE loop insertions in 2.5 million antibody sequences from HIV-uninfected individuals. We searched a set of ~24 million high-throughput sequences related to 13 studies of HIV-1 bnAbs to determine whether there were H4 and L4 insertions. These sequences are found across 13 HIV-1 high-throughput sequencing datasets related to the affinity maturation of VRC01, CH103, and PTG134-137 lineage antibodies, as well as co-evolution of HIV-1 bnAbs with their founder HIV-1 virus (17,46–51).

We identified potential insertions in the DE loop by examining the alignments of these sequences to hidden Markov models of the κ, λ, and heavy chain variable domain, and searching for gaps in the HMM consensus sequence within 10 amino acids before and after the DE loop. This resulted in 599 unique (637 total) heavy chain sequences, 521 unique (1,354 total) λ sequences, and 3,174 (6,352 total) κ sequences with amino acid insertions in, or around the DE loop. We used Clustal-Omega to align each group of sequences (heavy chain, κ, λ) separately and Jalview to edit the alignments and to analyze the DE-loop region. In each of the alignments, there were some sequences that contained substantial changes in the protein sequence in or near the DE loop. These changes are likely due to frameshifts caused by an insertion or deletion of one or more bases that are compensated by another insertion or deletion later in the nucleic acid sequence such that the frame is restored. In some cases, these may be due to sequencing errors. In other cases, if the frameshift covers a substantial region of the amino acid sequence, the antibody may not fold properly.

Antibody L4 sequences with κ germlines (Figure 10A) have the most insertions compared to heavy and λ DE loop sequences. By comparing the sequence of 5 amino acids before the DE loop, the DE loop itself, and 5 amino acids after the DE loop, 480 of 3174 sequences (15.6%) appear to have a frameshift mutation in or adjacent to the DE loop. Of the remaining sequences, 98% contain a length 8 DE loop, with the sequence in 85% of these containing a two-residue insertion resulting in a sequence resembling GSGSGTDF (e.g., a sequence from derived from IGKV4-4*01 inserts GS before GSGTDF in Figure 10A). The remainder are of length 7. Some sequences have short insertions, alongside somatic mutations (e.g. a human IGKV3D-11*01 sequence (germline sequence GPGTDF) has both a GS insertion before the DE loop, as well as eight-residue DE loop sequences ASAAGTEF and ASASGTDF).

**Figure 10.**
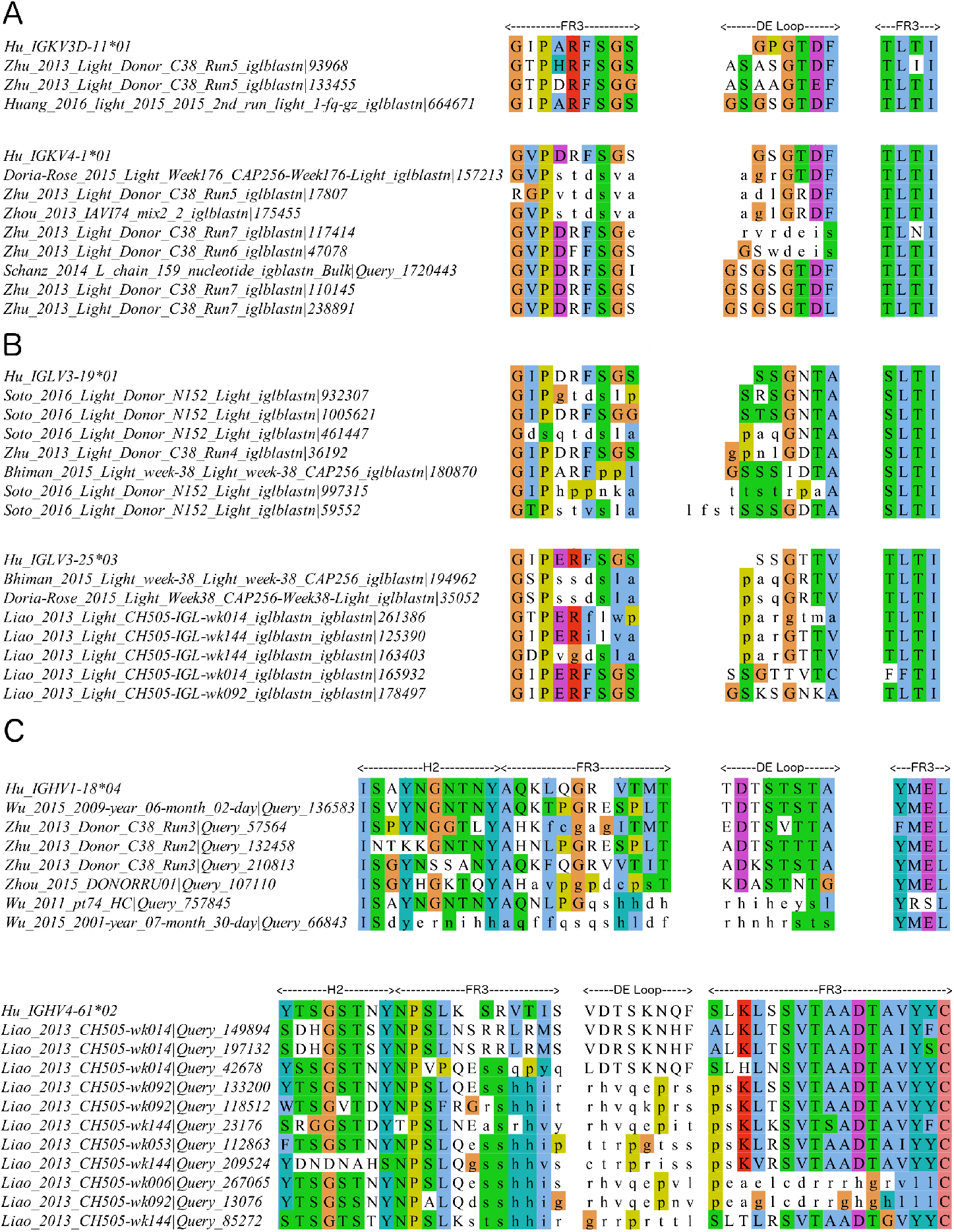
DE loop and DE loop adjacent insertions from a large antibody sequencing dataset from HIV-infected individuals. **A.** Insertions in κ gene antibodies. **B.** Insertions in λ gene antibodies. **C.** Insertions in heavy gene antibodies.

Antibody L4 sequences from λ germlines (Figure 10B) have fewer insertions than κ L4 sequences, but still have features of simple amino-acid insertions, frameshifting insertions, and somatic mutation alongside insertion. Frameshifts are associated with 21.5% of the 521 unique sequences in the λ data set. Of the remaining 409 sequences, 92.4% have DE loops of length 8 and the rest are of length 7. Similar to IGKV4-4*01, an IGLV3-12*02 sequence has an inserted GS before the DE loop and mutations of the germline DE loop from NPGNTA, resulting in the sequence GSKSGNKA. Sequences from IGLV3-19*01 and IGLV3-25*03 have both frameshifts, as well single amino-acid insertions that do not dramatically change the DE loop sequence (e.g., a single amino acid insertion changing the germline sequence from SSGNTA to STSGNTA).

In the heavy-chain alignment, we observed many sequences that are likely due to frameshifts in or adjacent to the DE loop. 108 sequences (17.9%) have an insertion that causes a frameshift prior to the DE loop that extends through the DE loop sequence. 284 sequences (47.4%) contain a frameshift that starts prior to the DE loop and extends through the beginning of H3. The remaining 207 heavy chain sequences (34.5%) appear to have an insertion of one amino acid in the loop that leads into the d strand and no insertion in the DE loop. Examples of each case are shown in Figure 10C. There are no clear examples of a simple insertion within the DE loop itself in the heavy chain sequences, and no cases that resemble the insertions in the HIV bNAbs in the PDB.

## Discussion

In this paper, we have analyzed sequence and structural features of the antibody DE loop. We have clustered DE loop conformations of the heavy and light chains, identified atomic interactions that are highly associated with various CDR1/CDR2 and DE loop pairs, and have shown features of affinity matured antibodies that utilize DE loops with somatic insertions to directly bind antigen. We identified nine clusters of the DE loop in heavy and light chains, and denoted them L4-6-1, L4-6-2, L4-6-3, L4-6-4, L4-8-1, H4-6-1, H4-7-1, H4-8-1, and H4-8-2. For each clustering of each length of L4 and H4, we have also provided support for each individual cluster by counting how many structures and unique sequences after enforcing a strict electron density fit calculation through the EDIA score (15). This allows us to find clusters that are not defined by mis-solved residues, such as peptide flips (55), and is an important step in validating clusters and backbone structures not only for clustering the DE loop, but the other CDRs as well as backbone structures from other proteins (64,65).

With this new structural classification and nomenclature to describe the DE loop of antibody structures, we argue that the DE loop structure and sequence should be analyzed when new antibody structures are determined and during development of antibody reagents and therapeutics. In some respects, the DE loop acts as a fourth CDR, since it behaves like the other CDRs: it binds antigen and it undergoes somatic mutations and insertions that directly or indirectly affect the antibody binding paratope at a rate that is higher than most framework residues (at least in the heavy chain and λ light-chain sequences).

Our analysis has the limitation that we cannot always determine causative associations between the sequence and structure of the DE loop and those of the CDRs. For example, even if we have some association between a less common DE loop conformation and CDR1 cluster, we cannot say that DE loop conformation is a determinant of the CDR1/CDR2 conformation. For example, L1-13-1 conformations are associated with L4-6-2, while L1-13-2 conformations are associated with L4-8-1. The sequence profiles of L1-13-1 and L1-13-2 are distinct. We do not know if an antibody was constructed with an L1-13-2 sequence and a typical L4-6-2 DE loop, whether the L1-13-2 conformation would be maintained. Therefore, when accounting for an impact that the DE loop may have on a neighboring CDR, it is important to note the L4 conformations available for that particular choice of L1 conformation, and also the sequence positions within a DE loop cluster that differentially impact CDR1 or CDR2 conformation. While some hydrogen bonds occur at a very high occupancy (90% or above) in specific DE loops pairings with CDR1/CDR2, others occur at a lower occupancy. Hydrogen bonds of this nature may not be a strict requirement for a particular L1 conformation, but may have an effect on stability of that L1 conformation, but without extensive additional experimental data to test this hypothesis, we cannot determine this.

Considering the DE loop as a fourth CDR suggests applications for antibody design and antibody modeling. For example, when designing antibodies using the ‘CDR grafting’ method (66), we suggest that whenever CDR1 is grafted on the light chain, or CDR1 or CDR2 on the heavy chain, L4 or H4 should be ‘co-grafted’ onto the same template structure. This method will preserve contacts between L4/L1, H4/H1, or H4/H2 that may be necessary for preserving or stabilizing the structures of CDR1 or CDR2.

When considering antibody structure prediction (67–69), a common strategy is to use CDR and framework templates based upon sequence similarity to known structures. We suggest extra attention to the relationships of L4 sequences with their structural clusters. For example, κ antibodies with a somatic mutation at the first position of the L4 from glycine to any other residue should be modeled with representative structures from cluster L4-6-2 instead of the more common L4-6-1 conformation. A similar approach can be considered when selecting templates for solving X-ray crystallographic structures by molecular replacement. Taking this information into account is more likely to recapitulate contacts observed in experimental structures. The appropriate cluster, and thus structure, for CDR1 and CDR2 often depends on the sequence and conformation of CDR4, and they should be modeled together in antibody structure prediction methods.

With high-throughput sequencing data in response to HIV-1, we have shown that the DE loop undergoes somatic mutation, alongside nucleotide insertions and deletions causing frameshift mutations in several human germline examples. Tracking useful features from the DE loop sequences that contribute to antigen binding, and ultimately neutralization of viral infections, may prove an important step in identifying functional antibodies from the human repertoire.

## Materials and Methods

### Antibody structure and sequence data

We compiled sequence and structure data for all antibodies from the Protein Data Bank (PDB). To collect the list of antibodies in the PDB, we used a lab maintained software, PyIgClassify (70). PyIgClassify compiles all antibody structures from the PDB by applying a set of hidden Markov models (HMMs) for each antibody gene (VH, Vl, and Vk) to all sequences in the PDB using HMMER3.0. PyIgClassify also renumbers antibodies according to a modified Honegger-Plückthun CDR scheme and numbering system described in North et al. (56,71) In order to identify CDRs in PyIgClassify, the software uses sequence alignment to the match states of the HMMs.

In order to identify which residues are structurally variable, we plotted ϕ and ψ for all residues in and around the solvent exposed DE loop (3 before the loop, and 3 after the loop, Figure 2). We updated PyIgClassify to recognize L4 and H4 in each antibody sequence, adding insert codes appropriately for loops longer the pre-allocated range of numbers.

We determine germline by comparing each PDB sequence to a curated set of IMGT germline protein sequences with BLAST, taking into account the author-provided species designation. However, these are often incorrect. We use a germline from a different species from the author-provided one if the sequence identity of the antibody is at least 8 percentage points higher than the author-provided species. This script also handles cases of ambiguous assignment such as humanized antibodies originating from non-human germlines. The dataset used in this study was compiled in March 2020 and includes data for 3,910 antibody structures containing 13,012 individual chains (available at http://dunbrack2.fccc.edu/PyIgClassify)

### Calculating electron density support for individual atoms (EDIA)

In order to add support for the electron density for the individual atoms for all structures within clusters, we calculated the EDIA score using the *ediascorer* standalone software from the University of Hamburg (15) (https://www.zbh.uni-hamburg.de/en/forschung/amd/software/edia.html). This software requires the structure factor file as well as the electron density map (.mtz). We downloaded both of these files from the PDB except for some older entries without deposited structure factor files. We then calculated the EDIA score for each backbone atom, and took as the EDIA score the value for the carbonyl oxygen, which is the most sensitive atom to the electron density fit. We established a second data set by eliminating any structure with one or more carbonyl oxygen atoms with an EDIA score of less than 0.75.

### Analyzing antibody-antigen complex set

For non-canonical length structures, we calculated the antibody-antigen buried surface area with the Rosetta macromolecular modeling suite (72). We calculated buried surface area as the change in antigen surface area of the CDR from the bound structure to unbound structure:

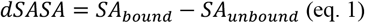

where SA represents the surface area calculated in Rosetta using the Shrake-Rupley algorithm (73) and a standard probe radius of 1.4 Å.

### Clustering loop structures

In order to group various conformations of L4 and H4 into structural families, we implemented a density based clustering method for dihedral angles based on the DBSCAN algorithm (53). This unsupervised learning method finds robust clusters by identifying dense regions in the metric space which are separated by low density. It also identifies “noise points,” which are outlier structures due to poor crystal structure determination or unusual mutations that cause uncommon structural changes. We used the implementation of DBSCAN in the sci-kit learn library (74) in python.

To compare two loops *i* and *j* with identical lengths, we first calculate the dihedral similarity between two angles θ_1_ and θ_2_ for each pair of corresponding residues, where θ represents any chosen backbone dihedral angle selected from ϕ, ψ, or ω:

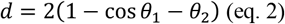

For our purposes we chose to include ϕ, ψ, and ω, which provides the maximum capability to resolve structures with both cis- and trans-peptide bonds. Next, we take as the final clustering distance the maximum value out of the set of calculations of *d* for {ϕ, ψ, ω}, which we call the *L*_∞_ norm:

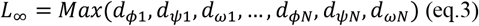

We chose the *L*_∞_ norm due to its sensitivity in separating loops which are different even at one single dihedral, giving our final clustering single dihedral resolution.

The resulting set of pairwise *L*_∞_ distances are then clustered from an *N*x*N* pairwise matrix using DBSCAN. This algorithm requires two parameters: ε and *MinPts*. The first parameter, ε, describes a distance from a given data point to search for neighboring data points. The second parameter, *MinPts*, specifies the requirement for the minimum number of neighboring data points within ε of a data point to label the data point under consideration a ‘core point’. Data points which are within ε of a core point, but do have *MinPts* data points within ε are called ‘border points’; points which do not meet either criterion are labeled as ‘noise points.’ The final clusters are the connected graphs of all of the core points, together with their border points.

Each selection of a combination of ε and *MinPts* produces a different set of clusters. Two main obstacles exist in identifying all of the interesting clusters from DBSCAN. First, at larger values of ε and smaller *MinPts*, DBSCAN may merge clusters that ought to be separated. Merged clusters are easily identified from their Ramachandran plot distributions at specific residues within a cluster (e.g., separated densities in the alpha and beta regions of the Ramachandran map). Second, valid, low-density clusters may only arise at larger values ε, while also producing undesirable merged clusters. This means that no singular selection of ε and *MinPts* will generate the entire set of valid clusters. To overcome these two issues, we developed a method to select a set of final clusters after running DBSCAN on a grid of ε and *MinPts*, by combining the results of each run of DBSCAN. First, we establish a parameter grid of ε and *MinPts* by selecting a range of both parameters, and run DBSCAN at each parameter selection. We then filter out any merged clusters by removing any clusters in which any two members of the cluster are more than 150° apart. This eliminates clusters that contain points in two or more Ramachandran regions (A, B, L, or E), the centroids of which are approximately 180° apart in ϕ or ψ. Next, the remaining clusters that pass the merge filtering criterion are treated as nodes on a graph, where the nodes have edges connected to them based on the calculation of Simpson’s similarity score (75):

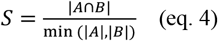

Finally, for each connected subgraph with *n* nodes, we take the final cluster of that subgraph as the union of all nodes *n* within the connected subgraph. This produces a final clustering set with clusters of varying density, without including merged clusters.

Following the determination of the final cluster set, we determined cluster representatives using angular statistical analysis. For a given cluster C consisting of N data points, for each structure *i* we calculate the average distance *d_i_* to all other points *j* in the same cluster C:

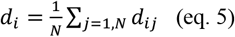

We choose the cluster representative as the structure which has the lowest *d_i_* of all of the structures.

### Identifying important hydrogen bonds between CDR4 and CDR1/CDR2

We calculated all hydrogen bonds between CDR4, CDR1, and CDR2 using Rosetta’s distance and orientation-dependent hydrogen bond energy accessed through the *report_hbonds_for_plugin.<release>* available in the public release of Rosetta3. We used the resulting contact information to find important contacts that are either frequent or unique over several CDR-lengths and germlines. We analyzed the hydrogen bonds between all CDR1-CDR4 and CDR2-CDR4 pairs for which both CDR1 and CDR4 have defined cluster membership. We then calculated the hydrogen bond occupancy for a particular hydrogen bond as the following:

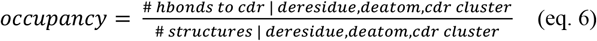

### High-throughput sequence analysis of naïve human antibodies

We accessed high-throughput sequencing data through the antibodymap.org server (www.antibodymap.org) (76). To gain an understanding of how variable L4 and H4 are compared to the other CDRs, we analyzed 12 human germlines (IGHV1-18*04, IGHV3-23*01, IGHV4-34*01, IGHV4-39*07, IGKV1-39*01, IGKV3-11*01, IGKV3-20*01, IGKV4-1*01, IGLV1-40*01, IGLV1-44*01, IGLV2-14*01, IGLV3-1*01) collected from naïve donor deep sequencing samples with thousands of sequences for each germline (download shell script included in supplementary data). Separately, we compared sequence variability between all human germlines for each heavy, λ, and κ gene compiled from IMGT for all germline sequences of the same length. We calculated sequence variability according to the Shannon entropy, denoted H, which represents a robust method to calculate antibody CDR variability (77):

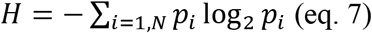

We calculated *H* only for residues up until the conserved cysteine before CDR3 on both the light and heavy chains.

### High-throughput sequence analysis of HIV-1 bnAbs

In order to search for insertions in L4 or H4 amongst HIV-1 infected patients, we collected all studies referring to HIV-1 from the antibodymap API (download shell script included in supplementary data). We identified CDRs for all of the FASTA files using the HMMER3.0 hmmsearch command, providing the profile HMMs implemented in PyIgClassify for IGHV, IGKV, IGLV, and IGLV6 genes (provided in supplementary data). We searched for sequences that had insertions compared to the profile, and examined these for features related to the long L4 or H4 structures we found in the PDB (sequences are provided in FASTA format in the supplementary data). From this set, we used Clustal-omega to align all of the sequences to the germline sequence which matched the IMGT germline assignment (provided in supplementary data). We observed frameshift mutations using the IMGT/V-QUEST tool (78), which notates nucleotide insertions that result in frameshifts.

## Acknowledgements

This work was funded by NIH grant R35 GM122517 (to R.L.D.) as well as the NIH T32 Structural Biology and Molecular Biophysics training grant awarded by the Department of Biochemistry and Molecular Biophysics at the University of Pennsylvania (GM008275).

---

**Table S1.**
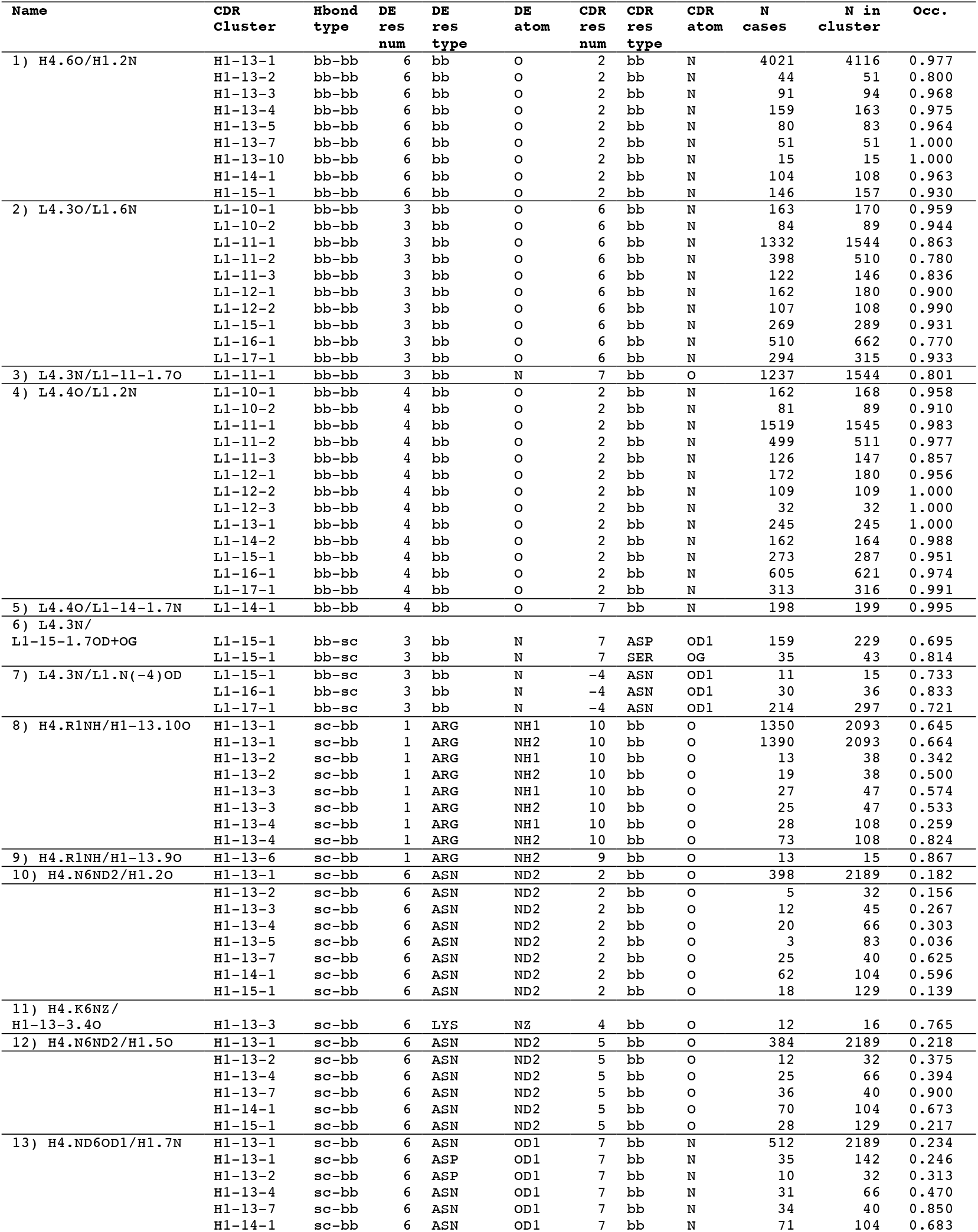
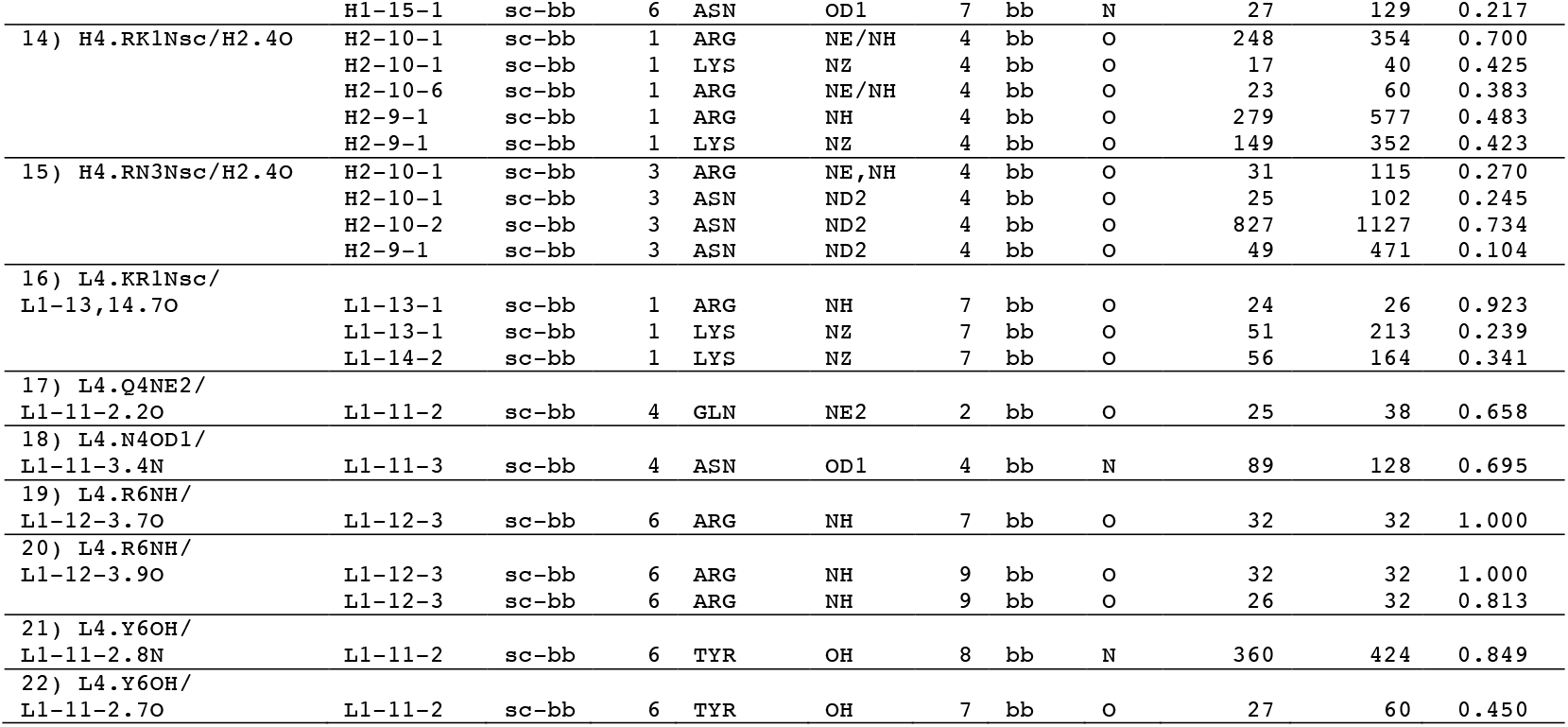
Hydrogen bonds between DE loop residues and CDRs.

