## Supplementary material for "Hiding in plain sight: structure and sequence analysis reveals the importance of the antibody DE loop for antibody-antigen binding": download_hiv_seqs.sh

```
wget http://opig.stats.ox.ac.uk/webapps/ngsdb/json/Schanz_2014/Schanz_2014_L_chain_159_wk181_Schanz_159_wk181_nucleotide_renamed_igblastn_Bulk.json.gz
wget http://opig.stats.ox.ac.uk/webapps/ngsdb/json/Schanz_2014/Schanz_2014_H_chain_159_wk181_IGHG_Schanz_159_wk181_nucleotide_renamed_igblastn_IGHG.json.gz
wget http://opig.stats.ox.ac.uk/webapps/ngsdb/json/Schanz_2014/Schanz_2014_H_chain_159_wk181_IGHE_Schanz_159_wk181_nucleotide_renamed_igblastn_IGHE.json.gz
wget http://opig.stats.ox.ac.uk/webapps/ngsdb/json/Schanz_2014/Schanz_2014_H_chain_159_wk94_IGHE_Schanz_159_wk94_nucleotide_renamed_igblastn_IGHE.json.gz
wget http://opig.stats.ox.ac.uk/webapps/ngsdb/json/Schanz_2014/Schanz_2014_H_chain_159_wk213_IGHG_Schanz_159_wk213_nucleotide_renamed_igblastn_IGHG.json.gz
wget http://opig.stats.ox.ac.uk/webapps/ngsdb/json/Schanz_2014/Schanz_2014_H_chain_159_wk213_IGHM_Schanz_159_wk213_nucleotide_renamed_igblastn_IGHM.json.gz
wget http://opig.stats.ox.ac.uk/webapps/ngsdb/json/Schanz_2014/Schanz_2014_H_chain_159_wk94_Bulk_Schanz_159_wk94_nucleotide_renamed_igblastn_Bulk.json.gz
wget http://opig.stats.ox.ac.uk/webapps/ngsdb/json/Schanz_2014/Schanz_2014_H_chain_159_wk213_IGHA_Schanz_159_wk213_nucleotide_renamed_igblastn_IGHA.json.gz
wget http://opig.stats.ox.ac.uk/webapps/ngsdb/json/Schanz_2014/Schanz_2014_H_chain_159_wk213_Bulk_Schanz_159_wk213_nucleotide_renamed_igblastn_Bulk.json.gz
wget http://opig.stats.ox.ac.uk/webapps/ngsdb/json/Schanz_2014/Schanz_2014_H_chain_159_wk94_IGHG_Schanz_159_wk94_nucleotide_renamed_igblastn_IGHG.json.gz
wget http://opig.stats.ox.ac.uk/webapps/ngsdb/json/Schanz_2014/Schanz_2014_L_chain_159_wk94_Schanz_159_wk94_nucleotide_renamed_igblastn_Bulk.json.gz
wget http://opig.stats.ox.ac.uk/webapps/ngsdb/json/Schanz_2014/Schanz_2014_H_chain_159_wk213_IGHE_Schanz_159_wk213_nucleotide_renamed_igblastn_IGHE.json.gz
wget http://opig.stats.ox.ac.uk/webapps/ngsdb/json/Schanz_2014/Schanz_2014_H_chain_159_wk181_IGHM_Schanz_159_wk181_nucleotide_renamed_igblastn_IGHM.json.gz
wget http://opig.stats.ox.ac.uk/webapps/ngsdb/json/Schanz_2014/Schanz_2014_H_chain_159_wk181_Bulk_Schanz_159_wk181_nucleotide_renamed_igblastn_Bulk.json.gz
wget http://opig.stats.ox.ac.uk/webapps/ngsdb/json/Schanz_2014/Schanz_2014_L_chain_159_wk213_Schanz_159_wk213_nucleotide_renamed_igblastn_Bulk.json.gz
wget http://opig.stats.ox.ac.uk/webapps/ngsdb/json/Schanz_2014/Schanz_2014_H_chain_159_wk94_IGHM_Schanz_159_wk94_nucleotide_renamed_igblastn_IGHM.json.gz
wget http://opig.stats.ox.ac.uk/webapps/ngsdb/json/Zhou_2013/Zhou_2013_IAVI23_VL2_2_iglblastn.json.gz
wget http://opig.stats.ox.ac.uk/webapps/ngsdb/json/Zhou_2013/Zhou_2013_RU3_VK1_2_iglblastn.json.gz
wget http://opig.stats.ox.ac.uk/webapps/ngsdb/json/Zhou_2013/Zhou_2013_NIAID45_VK3_2_iglblastn.json.gz
wget http://opig.stats.ox.ac.uk/webapps/ngsdb/json/Zhou_2013/Zhou_2013_IAVI57_VK1_2_iglblastn.json.gz
wget http://opig.stats.ox.ac.uk/webapps/ngsdb/json/Zhou_2013/Zhou_2013_IAVI74_mix2_2_iglblastn.json.gz
wget http://opig.stats.ox.ac.uk/webapps/ngsdb/json/Wu_2015/Wu_2015_Heavy_Visit_G_IGHG_H-1-chain_G-visit_2008-year_08-month_19-day_iglblastn_iglblastn_IGHG.json.gz
wget http://opig.stats.ox.ac.uk/webapps/ngsdb/json/Wu_2015/Wu_2015_Light_Visit_C_L-chain_C-visit_2006-year_07-month_14-day_iglblastn.json.gz
wget http://opig.stats.ox.ac.uk/webapps/ngsdb/json/Wu_2015/Wu_2015_Heavy_Visit_0_IGHE_H-chain_0-visit_1995-year_03-month_20-day_iglblastn_iglblastn_IGHE.json.gz
wget http://opig.stats.ox.ac.uk/webapps/ngsdb/json/Wu_2015/Wu_2015_Heavy_Visit_B_IGHM_H-chain_B-visit_2002-year_04-month_16-day_iglblastn_iglblastn_IGHM.json.gz
wget http://opig.stats.ox.ac.uk/webapps/ngsdb/json/Wu_2015/Wu_2015_Heavy_Visit_D_IGHG_H-chain_D-visit_2007-year_01-month_09-day_iglblastn_iglblastn_IGHG.json.gz
wget http://opig.stats.ox.ac.uk/webapps/ngsdb/json/Wu_2015/Wu_2015_Heavy_Visit_A_IGHE_H-4-chain_A-visit_2001-year_07-month_30-day_iglblastn_iglblastn_IGHE.json.gz
wget http://opig.stats.ox.ac.uk/webapps/ngsdb/json/Wu_2015/Wu_2015_Heavy_Visit_A_Bulk_H-3-chain_A-visit_2001-year_07-month_30-day_iglblastn_iglblastn_Bulk.json.gz
wget http://opig.stats.ox.ac.uk/webapps/ngsdb/json/Wu_2015/Wu_2015_Heavy_Visit_A_Bulk_H-4-chain_A-visit_2001-year_07-month_30-day_iglblastn_iglblastn_Bulk.json.gz
wget http://opig.stats.ox.ac.uk/webapps/ngsdb/json/Wu_2015/Wu_2015_Light_Visit_D_L-chain_D-visit_2007-year_01-month_09-day_iglblastn.json.gz
wget http://opig.stats.ox.ac.uk/webapps/ngsdb/json/Wu_2015/Wu_2015_Heavy_Visit_C_IGHE_H-1-chain_C-visit_2006-year_07-month_14-day_iglblastn_iglblastn_IGHE.json.gz
wget http://opig.stats.ox.ac.uk/webapps/ngsdb/json/Wu_2015/Wu_2015_Heavy_Visit_G_Bulk_H-chain_G-visit_2008-year_08-month_19-day_iglblastn_iglblastn_Bulk.json.gz
wget http://opig.stats.ox.ac.uk/webapps/ngsdb/json/Wu_2015/Wu_2015_Heavy_Visit_G_IGHE_H-chain_G-visit_2008-year_08-month_19-day_iglblastn_iglblastn_IGHE.json.gz
wget http://opig.stats.ox.ac.uk/webapps/ngsdb/json/Wu_2015/Wu_2015_Heavy_Visit_A_IGHG_H-1-chain_A-visit_2001-year_07-month_30-day_iglblastn_iglblastn_IGHG.json.gz
wget http://opig.stats.ox.ac.uk/webapps/ngsdb/json/Wu_2015/Wu_2015_Heavy_Visit_D_Bulk_H-chain_D-visit_2007-year_01-month_09-day_iglblastn_iglblastn_Bulk.json.gz
wget http://opig.stats.ox.ac.uk/webapps/ngsdb/json/Wu_2015/Wu_2015_Heavy_Visit_0_IGHM_H-2-chain_0-visit_1995-year_03-month_20-day_iglblastn_iglblastn_IGHM.json.gz
wget http://opig.stats.ox.ac.uk/webapps/ngsdb/json/Wu_2015/Wu_2015_Heavy_Visit_B_IGHG_H-chain_B-visit_2002-year_04-month_16-day_iglblastn_iglblastn_IGHG.json.gz
wget http://opig.stats.ox.ac.uk/webapps/ngsdb/json/Wu_2015/Wu_2015_Heavy_Visit_0_IGHD_H-2-chain_0-visit_1995-year_03-month_20-day_iglblastn_iglblastn_IGHD.json.gz
wget http://opig.stats.ox.ac.uk/webapps/ngsdb/json/Wu_2015/Wu_2015_Light_Visit_I_L-chain_I-visit_2009-year_12-month_31-day_iglblastn.json.gz
wget http://opig.stats.ox.ac.uk/webapps/ngsdb/json/Wu_2015/Wu_2015_Heavy_Visit_B_Bulk_H-chain_B-visit_2002-year_04-month_16-day_iglblastn_iglblastn_Bulk.json.gz
wget http://opig.stats.ox.ac.uk/webapps/ngsdb/json/Wu_2015/Wu_2015_Heavy_Visit_G_IGHM_H-2-chain_G-visit_2008-year_08-month_19-day_iglblastn_iglblastn_IGHM.json.gz
wget http://opig.stats.ox.ac.uk/webapps/ngsdb/json/Wu_2015/Wu_2015_Heavy_Visit_C_Bulk_H-chain_C-visit_2006-year_07-month_14-day_iglblastn_iglblastn_Bulk.json.gz
wget http://opig.stats.ox.ac.uk/webapps/ngsdb/json/Wu_2015/Wu_2015_Heavy_Visit_A_IGHM_H-chain_A-visit_2001-year_07-month_30-day_iglblastn_iglblastn_IGHM.json.gz
wget http://opig.stats.ox.ac.uk/webapps/ngsdb/json/Wu_2015/Wu_2015_Heavy_Visit_G_IGHE_H-2-chain_G-visit_2008-year_08-month_19-day_iglblastn_iglblastn_IGHE.json.gz
wget http://opig.stats.ox.ac.uk/webapps/ngsdb/json/Wu_2015/Wu_2015_Heavy_Visit_D_IGHE_H-chain_D-visit_2007-year_01-month_09-day_iglblastn_iglblastn_IGHE.json.gz
wget http://opig.stats.ox.ac.uk/webapps/ngsdb/json/Wu_2015/Wu_2015_Heavy_Visit_A_IGHM_H-4-chain_A-visit_2001-year_07-month_30-day_iglblastn_iglblastn_IGHM.json.gz
wget http://opig.stats.ox.ac.uk/webapps/ngsdb/json/Wu_2015/Wu_2015_Light_Visit_E_L-chain_E-visit_2007-year_07-month_12-day_iglblastn.json.gz
wget http://opig.stats.ox.ac.uk/webapps/ngsdb/json/Wu_2015/Wu_2015_Heavy_Visit_A_IGHM_H-2-chain_A-visit_2001-year_07-month_30-day_iglblastn_iglblastn_IGHM.json.gz
wget http://opig.stats.ox.ac.uk/webapps/ngsdb/json/Wu_2015/Wu_2015_Heavy_Visit_F_IGHA_H-RERUN-G1-chain_F-visit_2008-year_01-month_17-day_iglblastn_iglblastn_IGHA.json.gz
wget http://opig.stats.ox.ac.uk/webapps/ngsdb/json/Wu_2015/Wu_2015_Heavy_Visit_F_Bulk_H-RERUN-G1-chain_F-visit_2008-year_01-month_17-day_iglblastn_iglblastn_Bulk.json.gz
wget http://opig.stats.ox.ac.uk/webapps/ngsdb/json/Wu_2015/Wu_2015_Heavy_Visit_A_IGHM_H-1-chain_A-visit_2001-year_07-month_30-day_iglblastn_iglblastn_IGHM.json.gz
wget http://opig.stats.ox.ac.uk/webapps/ngsdb/json/Wu_2015/Wu_2015_Heavy_Visit_G_Bulk_H-2-chain_G-visit_2008-year_08-month_19-day_iglblastn_iglblastn_Bulk.json.gz
wget http://opig.stats.ox.ac.uk/webapps/ngsdb/json/Wu_2015/Wu_2015_Heavy_Visit_G_Bulk_H-1-chain_G-visit_2008-year_08-month_19-day_iglblastn_iglblastn_Bulk.json.gz
wget http://opig.stats.ox.ac.uk/webapps/ngsdb/json/Wu_2015/Wu_2015_Heavy_Visit_I_IGHD_H-chain_I-visit_2009-year_12-month_31-day_iglblastn_iglblastn_IGHD.json.gz
wget http://opig.stats.ox.ac.uk/webapps/ngsdb/json/Wu_2015/Wu_2015_Heavy_Visit_A_IGHD_H-2-chain_A-visit_2001-year_07-month_30-day_iglblastn_iglblastn_IGHD.json.gz
wget http://opig.stats.ox.ac.uk/webapps/ngsdb/json/Wu_2015/Wu_2015_Heavy_Visit_A_IGHE_H-2-chain_A-visit_2001-year_07-month_30-day_iglblastn_iglblastn_IGHE.json.gz
wget http://opig.stats.ox.ac.uk/webapps/ngsdb/json/Wu_2015/Wu_2015_Heavy_Visit_H_IGHE_H-chain_H-visit_2009-year_06-month_02-day_iglblastn_iglblastn_IGHE.json.gz
wget http://opig.stats.ox.ac.uk/webapps/ngsdb/json/Wu_2015/Wu_2015_Heavy_Visit_0_IGHD_H-chain_0-visit_1995-year_03-month_20-day_iglblastn_iglblastn_IGHD.json.gz
wget http://opig.stats.ox.ac.uk/webapps/ngsdb/json/Wu_2015/Wu_2015_Heavy_Visit_E_IGHG_H-chain_E-visit_2007-year_07-month_12-day_iglblastn_iglblastn_IGHG.json.gz
wget http://opig.stats.ox.ac.uk/webapps/ngsdb/json/Wu_2015/Wu_2015_Heavy_Visit_0_Bulk_H-chain_0-visit_1995-year_03-month_20-day_iglblastn_iglblastn_Bulk.json.gz
wget http://opig.stats.ox.ac.uk/webapps/ngsdb/json/Wu_2015/Wu_2015_Heavy_Visit_F_Bulk_H-chain_F-visit_2008-year_01-month_17-day_iglblastn_iglblastn_Bulk.json.gz
wget http://opig.stats.ox.ac.uk/webapps/ngsdb/json/Wu_2015/Wu_2015_Heavy_Visit_A_IGHE_H-chain_A-visit_2001-year_07-month_30-day_iglblastn_iglblastn_IGHE.json.gz
wget http://opig.stats.ox.ac.uk/webapps/ngsdb/json/Wu_2015/Wu_2015_Heavy_Visit_B_IGHA_H-chain_B-visit_2002-year_04-month_16-day_iglblastn_iglblastn_IGHA.json.gz
wget http://opig.stats.ox.ac.uk/webapps/ngsdb/json/Wu_2015/Wu_2015_Heavy_Visit_A_IGHG_H-3-chain_A-visit_2001-year_07-month_30-day_iglblastn_iglblastn_IGHG.json.gz
wget http://opig.stats.ox.ac.uk/webapps/ngsdb/json/Wu_2015/Wu_2015_Light_Visit_B_L-chain_B-visit_2002-year_04-month_16-day_iglblastn.json.gz
wget http://opig.stats.ox.ac.uk/webapps/ngsdb/json/Wu_2015/Wu_2015_Heavy_Visit_F_IGHG_H-chain_F-visit_2008-year_01-month_17-day_iglblastn_iglblastn_IGHG.json.gz
wget http://opig.stats.ox.ac.uk/webapps/ngsdb/json/Wu_2015/Wu_2015_Light_Visit_H_L-chain_H-visit_2009-year_06-month_02-day_iglblastn.json.gz
wget http://opig.stats.ox.ac.uk/webapps/ngsdb/json/Wu_2015/Wu_2015_Light_Visit_0_L-chain_0-visit_1995-year_03-month_20-day_iglblastn.json.gz
wget http://opig.stats.ox.ac.uk/webapps/ngsdb/json/Wu_2015/Wu_2015_Heavy_Visit_I_IGHM_H-chain_I-visit_2009-year_12-month_31-day_iglblastn_iglblastn_IGHM.json.gz
wget http://opig.stats.ox.ac.uk/webapps/ngsdb/json/Wu_2015/Wu_2015_Heavy_Visit_G_IGHG_H-2-chain_G-visit_2008-year_08-month_19-day_iglblastn_iglblastn_IGHG.json.gz
wget http://opig.stats.ox.ac.uk/webapps/ngsdb/json/Wu_2015/Wu_2015_Heavy_Visit_B_IGHE_H-chain_B-visit_2002-year_04-month_16-day_iglblastn_iglblastn_IGHE.json.gz
wget http://opig.stats.ox.ac.uk/webapps/ngsdb/json/Wu_2015/Wu_2015_Heavy_Visit_C_IGHM_H-chain_C-visit_2006-year_07-month_14-day_iglblastn_iglblastn_IGHM.json.gz
wget http://opig.stats.ox.ac.uk/webapps/ngsdb/json/Wu_2015/Wu_2015_Heavy_Visit_I_IGHG_H-chain_I-visit_2009-year_12-month_31-day_iglblastn_iglblastn_IGHG.json.gz
wget http://opig.stats.ox.ac.uk/webapps/ngsdb/json/Wu_2015/Wu_2015_Heavy_Visit_A_Bulk_H-1-chain_A-visit_2001-year_07-month_30-day_iglblastn_iglblastn_Bulk.json.gz
wget http://opig.stats.ox.ac.uk/webapps/ngsdb/json/Wu_2015/Wu_2015_Heavy_Visit_C_IGHG_H-chain_C-visit_2006-year_07-month_14-day_iglblastn_iglblastn_IGHG.json.gz
wget http://opig.stats.ox.ac.uk/webapps/ngsdb/json/Wu_2015/Wu_2015_Heavy_Visit_H_Bulk_H-chain_H-visit_2009-year_06-month_02-day_iglblastn_iglblastn_Bulk.json.gz
wget http://opig.stats.ox.ac.uk/webapps/ngsdb/json/Wu_2015/Wu_2015_Heavy_Visit_F_IGHG_H-RERUN-G1-chain_F-visit_2008-year_01-month_17-day_iglblastn_iglblastn_IGHG.json.gz
wget http://opig.stats.ox.ac.uk/webapps/ngsdb/json/Wu_2015/Wu_2015_Heavy_Visit_0_IGHE_H-2-chain_0-visit_1995-year_03-month_20-day_iglblastn_iglblastn_IGHE.json.gz
wget http://opig.stats.ox.ac.uk/webapps/ngsdb/json/Wu_2015/Wu_2015_Heavy_Visit_C_Bulk_H-1-chain_C-visit_2006-year_07-month_14-day_iglblastn_iglblastn_Bulk.json.gz
wget http://opig.stats.ox.ac.uk/webapps/ngsdb/json/Wu_2015/Wu_2015_Heavy_Visit_C_IGHM_H-1-chain_C-visit_2006-year_07-month_14-day_iglblastn_iglblastn_IGHM.json.gz
wget http://opig.stats.ox.ac.uk/webapps/ngsdb/json/Wu_2015/Wu_2015_Light_Visit_F_L-chain_F-visit_2008-year_01-month_17-day_iglblastn.json.gz
wget http://opig.stats.ox.ac.uk/webapps/ngsdb/json/Wu_2015/Wu_2015_Heavy_Visit_H_IGHM_H-chain_H-visit_2009-year_06-month_02-day_iglblastn_iglblastn_IGHM.json.gz
wget http://opig.stats.ox.ac.uk/webapps/ngsdb/json/Wu_2015/Wu_2015_Heavy_Visit_A_IGHG_H-2-chain_A-visit_2001-year_07-month_30-day_iglblastn_iglblastn_IGHG.json.gz
wget http://opig.stats.ox.ac.uk/webapps/ngsdb/json/Wu_2015/Wu_2015_Heavy_Visit_F_IGHM_H-RERUN-G1-chain_F-visit_2008-year_01-month_17-day_iglblastn_iglblastn_IGHM.json.gz
wget http://opig.stats.ox.ac.uk/webapps/ngsdb/json/Wu_2015/Wu_2015_Heavy_Visit_C_IGHG_H-1-chain_C-visit_2006-year_07-month_14-day_iglblastn_iglblastn_IGHG.json.gz
wget http://opig.stats.ox.ac.uk/webapps/ngsdb/json/Wu_2015/Wu_2015_Heavy_Visit_A_IGHG_H-4-chain_A-visit_2001-year_07-month_30-day_iglblastn_iglblastn_IGHG.json.gz
wget http://opig.stats.ox.ac.uk/webapps/ngsdb/json/Wu_2015/Wu_2015_Heavy_Visit_G_IGHA_H-chain_G-visit_2008-year_08-month_19-day_iglblastn_iglblastn_IGHA.json.gz
wget http://opig.stats.ox.ac.uk/webapps/ngsdb/json/Wu_2015/Wu_2015_Light_Visit_G_L-chain_G-visit_2008-year_08-month_19-day_iglblastn.json.gz
wget http://opig.stats.ox.ac.uk/webapps/ngsdb/json/Wu_2015/Wu_2015_Heavy_Visit_G_IGHM_H-1-chain_G-visit_2008-year_08-month_19-day_iglblastn_iglblastn_IGHM.json.gz
wget http://opig.stats.ox.ac.uk/webapps/ngsdb/json/Wu_2015/Wu_2015_Heavy_Visit_0_IGHM_H-chain_0-visit_1995-year_03-month_20-day_iglblastn_iglblastn_IGHM.json.gz
wget http://opig.stats.ox.ac.uk/webapps/ngsdb/json/Wu_2015/Wu_2015_Heavy_Visit_A_IGHD_H-3-chain_A-visit_2001-year_07-month_30-day_iglblastn_iglblastn_IGHD.json.gz
wget http://opig.stats.ox.ac.uk/webapps/ngsdb/json/Wu_2015/Wu_2015_Heavy_Visit_I_IGHE_H-chain_I-visit_2009-year_12-month_31-day_iglblastn_iglblastn_IGHE.json.gz
wget http://opig.stats.ox.ac.uk/webapps/ngsdb/json/Wu_2015/Wu_2015_Heavy_Visit_I_Bulk_H-chain_I-visit_2009-year_12-month_31-day_iglblastn_iglblastn_Bulk.json.gz
wget http://opig.stats.ox.ac.uk/webapps/ngsdb/json/Wu_2015/Wu_2015_Heavy_Visit_G_IGHM_H-chain_G-visit_2008-year_08-month_19-day_iglblastn_iglblastn_IGHM.json.gz
wget http://opig.stats.ox.ac.uk/webapps/ngsdb/json/Wu_2015/Wu_2015_Heavy_Visit_0_IGHG_H-chain_0-visit_1995-year_03-month_20-day_iglblastn_iglblastn_IGHG.json.gz
wget http://opig.stats.ox.ac.uk/webapps/ngsdb/json/Wu_2015/Wu_2015_Heavy_Visit_E_IGHM_H-chain_E-visit_2007-year_07-month_12-day_iglblastn_iglblastn_IGHM.json.gz
wget http://opig.stats.ox.ac.uk/webapps/ngsdb/json/Wu_2015/Wu_2015_Heavy_Visit_C_IGHE_H-chain_C-visit_2006-year_07-month_14-day_iglblastn_iglblastn_IGHE.json.gz
wget http://opig.stats.ox.ac.uk/webapps/ngsdb/json/Wu_2015/Wu_2015_Heavy_Visit_G_IGHD_H-chain_G-visit_2008-year_08-month_19-day_iglblastn_iglblastn_IGHD.json.gz
wget http://opig.stats.ox.ac.uk/webapps/ngsdb/json/Wu_2015/Wu_2015_Heavy_Visit_A_IGHE_H-3-chain_A-visit_2001-year_07-month_30-day_iglblastn_iglblastn_IGHE.json.gz
wget http://opig.stats.ox.ac.uk/webapps/ngsdb/json/Wu_2015/Wu_2015_Heavy_Visit_F_IGHE_H-RERUN-G1-chain_F-visit_2008-year_01-month_17-day_iglblastn_iglblastn_IGHE.json.gz
wget http://opig.stats.ox.ac.uk/webapps/ngsdb/json/Wu_2015/Wu_2015_Heavy_Visit_F_IGHM_H-chain_F-visit_2008-year_01-month_17-day_iglblastn_iglblastn_IGHM.json.gz
wget http://opig.stats.ox.ac.uk/webapps/ngsdb/json/Wu_2015/Wu_2015_Heavy_Visit_A_Bulk_H-2-chain_A-visit_2001-year_07-month_30-day_iglblastn_iglblastn_Bulk.json.gz
wget http://opig.stats.ox.ac.uk/webapps/ngsdb/json/Wu_2015/Wu_2015_Heavy_Visit_0_Bulk_H-2-chain_0-visit_1995-year_03-month_20-day_iglblastn_iglblastn_Bulk.json.gz
wget http://opig.stats.ox.ac.uk/webapps/ngsdb/json/Wu_2015/Wu_2015_Heavy_Visit_D_IGHM_H-chain_D-visit_2007-year_01-month_09-day_iglblastn_iglblastn_IGHM.json.gz
wget http://opig.stats.ox.ac.uk/webapps/ngsdb/json/Wu_2015/Wu_2015_Heavy_Visit_E_IGHE_H-chain_E-visit_2007-year_07-month_12-day_iglblastn_iglblastn_IGHE.json.gz
wget http://opig.stats.ox.ac.uk/webapps/ngsdb/json/Wu_2015/Wu_2015_Heavy_Visit_G_IGHE_H-1-chain_G-visit_2008-year_08-month_19-day_iglblastn_iglblastn_IGHE.json.gz
wget http://opig.stats.ox.ac.uk/webapps/ngsdb/json/Wu_2015/Wu_2015_Light_Visit_A_L-chain_A-visit_2001-year_07-month_30-day_iglblastn.json.gz
wget http://opig.stats.ox.ac.uk/webapps/ngsdb/json/Wu_2015/Wu_2015_Heavy_Visit_A_IGHE_H-1-chain_A-visit_2001-year_07-month_30-day_iglblastn_iglblastn_IGHE.json.gz
wget http://opig.stats.ox.ac.uk/webapps/ngsdb/json/Wu_2015/Wu_2015_Heavy_Visit_D_IGHD_H-chain_D-visit_2007-year_01-month_09-day_iglblastn_iglblastn_IGHD.json.gz
wget http://opig.stats.ox.ac.uk/webapps/ngsdb/json/Wu_2015/Wu_2015_Heavy_Visit_0_IGHA_H-2-chain_0-visit_1995-year_03-month_20-day_iglblastn_iglblastn_IGHA.json.gz
wget http://opig.stats.ox.ac.uk/webapps/ngsdb/json/Wu_2015/Wu_2015_Heavy_Visit_E_Bulk_H-chain_E-visit_2007-year_07-month_12-day_iglblastn_iglblastn_Bulk.json.gz
wget http://opig.stats.ox.ac.uk/webapps/ngsdb/json/Wu_2015/Wu_2015_Heavy_Visit_G_IGHG_H-chain_G-visit_2008-year_08-month_19-day_iglblastn_iglblastn_IGHG.json.gz
wget http://opig.stats.ox.ac.uk/webapps/ngsdb/json/Wu_2015/Wu_2015_Heavy_Visit_A_Bulk_H-chain_A-visit_2001-year_07-month_30-day_iglblastn_iglblastn_Bulk.json.gz
wget http://opig.stats.ox.ac.uk/webapps/ngsdb/json/Wu_2015/Wu_2015_Heavy_Visit_H_IGHG_H-chain_H-visit_2009-year_06-month_02-day_iglblastn_iglblastn_IGHG.json.gz
wget http://opig.stats.ox.ac.uk/webapps/ngsdb/json/Wu_2015/Wu_2015_Heavy_Visit_A_IGHD_H-4-chain_A-visit_2001-year_07-month_30-day_iglblastn_iglblastn_IGHD.json.gz
wget http://opig.stats.ox.ac.uk/webapps/ngsdb/json/Wu_2015/Wu_2015_Heavy_Visit_F_IGHG_H-1-chain_F-visit_2008-year_01-month_17-day_iglblastn_iglblastn_IGHG.json.gz
wget http://opig.stats.ox.ac.uk/webapps/ngsdb/json/Wu_2015/Wu_2015_Heavy_Visit_A_IGHG_H-chain_A-visit_2001-year_07-month_30-day_iglblastn_iglblastn_IGHG.json.gz
wget http://opig.stats.ox.ac.uk/webapps/ngsdb/json/Wu_2015/Wu_2015_Heavy_Visit_0_IGHG_H-2-chain_0-visit_1995-year_03-month_20-day_iglblastn_iglblastn_IGHG.json.gz
wget http://opig.stats.ox.ac.uk/webapps/ngsdb/json/Wu_2015/Wu_2015_Heavy_Visit_E_IGHD_H-chain_E-visit_2007-year_07-month_12-day_iglblastn_iglblastn_IGHD.json.gz
wget http://opig.stats.ox.ac.uk/webapps/ngsdb/json/Wu_2015/Wu_2015_Heavy_Visit_A_IGHM_H-3-chain_A-visit_2001-year_07-month_30-day_iglblastn_iglblastn_IGHM.json.gz
wget http://opig.stats.ox.ac.uk/webapps/ngsdb/json/Liao_2013/Liao_2013_Heavy_Bulk_CH505-IGH-wk006_iglblastn_igblastn_Bulk.json.gz
wget http://opig.stats.ox.ac.uk/webapps/ngsdb/json/Liao_2013/Liao_2013_Heavy_IGHD_CH505-IGH-wk092_iglblastn_igblastn_IGHD.json.gz
wget http://opig.stats.ox.ac.uk/webapps/ngsdb/json/Liao_2013/Liao_2013_Heavy_IGHM_CH505-IGH-wk006_iglblastn_igblastn_IGHM.json.gz
wget http://opig.stats.ox.ac.uk/webapps/ngsdb/json/Liao_2013/Liao_2013_Heavy_IGHD_CH505-IGH-wk014_iglblastn_igblastn_IGHD.json.gz
wget http://opig.stats.ox.ac.uk/webapps/ngsdb/json/Liao_2013/Liao_2013_Heavy_Bulk_CH505-IGH-wk014_iglblastn_igblastn_Bulk.json.gz
wget http://opig.stats.ox.ac.uk/webapps/ngsdb/json/Liao_2013/Liao_2013_Heavy_IGHG_CH505-IGH-wk006_iglblastn_igblastn_IGHG.json.gz
wget http://opig.stats.ox.ac.uk/webapps/ngsdb/json/Liao_2013/Liao_2013_Heavy_IGHG_CH505-IGH-wk144_iglblastn_igblastn_IGHG.json.gz
wget http://opig.stats.ox.ac.uk/webapps/ngsdb/json/Liao_2013/Liao_2013_Heavy_Bulk_CH505-IGH-wk144_iglblastn_igblastn_Bulk.json.gz
wget http://opig.stats.ox.ac.uk/webapps/ngsdb/json/Liao_2013/Liao_2013_Heavy_IGHE_CH505-IGH-wk144_iglblastn_igblastn_IGHE.json.gz
wget http://opig.stats.ox.ac.uk/webapps/ngsdb/json/Liao_2013/Liao_2013_Light_CH505-IGL-wk014_iglblastn_igblastn.json.gz
wget http://opig.stats.ox.ac.uk/webapps/ngsdb/json/Liao_2013/Liao_2013_Heavy_IGHE_CH505-IGH-wk006_iglblastn_igblastn_IGHE.json.gz
wget http://opig.stats.ox.ac.uk/webapps/ngsdb/json/Liao_2013/Liao_2013_Light_CH505-IGL-wk053_iglblastn_igblastn.json.gz
wget http://opig.stats.ox.ac.uk/webapps/ngsdb/json/Liao_2013/Liao_2013_Light_CH505-IGL-wk092_iglblastn_igblastn.json.gz
wget http://opig.stats.ox.ac.uk/webapps/ngsdb/json/Liao_2013/Liao_2013_Heavy_Bulk_CH505-IGH-wk053_iglblastn_igblastn_Bulk.json.gz
wget http://opig.stats.ox.ac.uk/webapps/ngsdb/json/Liao_2013/Liao_2013_Heavy_IGHM_CH505-IGH-wk014_iglblastn_igblastn_IGHM.json.gz
wget http://opig.stats.ox.ac.uk/webapps/ngsdb/json/Liao_2013/Liao_2013_Heavy_IGHG_CH505-IGH-wk014_iglblastn_igblastn_IGHG.json.gz
wget http://opig.stats.ox.ac.uk/webapps/ngsdb/json/Liao_2013/Liao_2013_Heavy_IGHG_CH505-IGH-wk053_iglblastn_igblastn_IGHG.json.gz
wget http://opig.stats.ox.ac.uk/webapps/ngsdb/json/Liao_2013/Liao_2013_Heavy_IGHG_CH505-IGH-wk092_iglblastn_igblastn_IGHG.json.gz
wget http://opig.stats.ox.ac.uk/webapps/ngsdb/json/Liao_2013/Liao_2013_Heavy_IGHE_CH505-IGH-wk053_iglblastn_igblastn_IGHE.json.gz
wget http://opig.stats.ox.ac.uk/webapps/ngsdb/json/Liao_2013/Liao_2013_Heavy_IGHE_CH505-IGH-wk014_iglblastn_igblastn_IGHE.json.gz
wget http://opig.stats.ox.ac.uk/webapps/ngsdb/json/Liao_2013/Liao_2013_Light_CH505-IGL-wk144_iglblastn_igblastn.json.gz
wget http://opig.stats.ox.ac.uk/webapps/ngsdb/json/Liao_2013/Liao_2013_Heavy_IGHM_CH505-IGH-wk092_iglblastn_igblastn_IGHM.json.gz
wget http://opig.stats.ox.ac.uk/webapps/ngsdb/json/Liao_2013/Liao_2013_Heavy_Bulk_CH505-IGH-wk092_iglblastn_igblastn_Bulk.json.gz
wget http://opig.stats.ox.ac.uk/webapps/ngsdb/json/Liao_2013/Liao_2013_Light_CH505-IGL-wk006_iglblastn_igblastn.json.gz
wget http://opig.stats.ox.ac.uk/webapps/ngsdb/json/Liao_2013/Liao_2013_Heavy_IGHM_CH505-IGH-wk053_iglblastn_igblastn_IGHM.json.gz
wget http://opig.stats.ox.ac.uk/webapps/ngsdb/json/Liao_2013/Liao_2013_Heavy_IGHM_CH505-IGH-wk144_iglblastn_igblastn_IGHM.json.gz
wget http://opig.stats.ox.ac.uk/webapps/ngsdb/json/Liao_2013/Liao_2013_Heavy_IGHE_CH505-IGH-wk092_iglblastn_igblastn_IGHE.json.gz
wget http://opig.stats.ox.ac.uk/webapps/ngsdb/json/Wu_2011/Wu_2011_Heavy_IGHG_pt74_HC_iglblastn_igblastn_IGHG.json.gz
wget http://opig.stats.ox.ac.uk/webapps/ngsdb/json/Wu_2011/Wu_2011_Heavy_IGHG_pt45_HC_iglblastn_igblastn_IGHG.json.gz
wget http://opig.stats.ox.ac.uk/webapps/ngsdb/json/Wu_2011/Wu_2011_Heavy_IGHE_pt45_HC_iglblastn_igblastn_IGHE.json.gz
wget http://opig.stats.ox.ac.uk/webapps/ngsdb/json/Wu_2011/Wu_2011_Heavy_IGHE_pt74_HC_iglblastn_igblastn_IGHE.json.gz
wget http://opig.stats.ox.ac.uk/webapps/ngsdb/json/Wu_2011/Wu_2011_Heavy_Bulk_pt74_HC_iglblastn_igblastn_Bulk.json.gz
wget http://opig.stats.ox.ac.uk/webapps/ngsdb/json/Wu_2011/Wu_2011_Heavy_IGHM_pt74_HC_iglblastn_igblastn_IGHM.json.gz
wget http://opig.stats.ox.ac.uk/webapps/ngsdb/json/Wu_2011/Wu_2011_Light_pt45_kappa_iglblastn.json.gz
wget http://opig.stats.ox.ac.uk/webapps/ngsdb/json/Wu_2011/Wu_2011_Heavy_IGHM_pt45_HC_iglblastn_igblastn_IGHM.json.gz
wget http://opig.stats.ox.ac.uk/webapps/ngsdb/json/Wu_2011/Wu_2011_Heavy_Bulk_pt45_HC_iglblastn_igblastn_Bulk.json.gz
wget http://opig.stats.ox.ac.uk/webapps/ngsdb/json/Huang_2016/Huang_2016_heavy_2014_Bulk_2014_1st_run_heavy_2-fq-gz_iglblastn_Bulk.json.gz
wget http://opig.stats.ox.ac.uk/webapps/ngsdb/json/Huang_2016/Huang_2016_heavy_2015_Bulk_2015_3rd_run_heavy_2-fq-gz_iglblastn_Bulk.json.gz
wget http://opig.stats.ox.ac.uk/webapps/ngsdb/json/Huang_2016/Huang_2016_heavy_2015_Bulk_2015_1st_run_heavy_1-fq-gz_iglblastn_Bulk.json.gz
wget http://opig.stats.ox.ac.uk/webapps/ngsdb/json/Huang_2016/Huang_2016_light_2014_2014_2nd_run_light_1-fq-gz_iglblastn.json.gz
wget http://opig.stats.ox.ac.uk/webapps/ngsdb/json/Huang_2016/Huang_2016_heavy_2012_Bulk_2012_1st_run_heavy_1-fq-gz_iglblastn_Bulk.json.gz
wget http://opig.stats.ox.ac.uk/webapps/ngsdb/json/Huang_2016/Huang_2016_heavy_2014_Bulk_2014_2nd_run_heavy_2-fq-gz_iglblastn_Bulk.json.gz
wget http://opig.stats.ox.ac.uk/webapps/ngsdb/json/Huang_2016/Huang_2016_light_2012_2012_1st_run_light_1-fq-gz_iglblastn.json.gz
wget http://opig.stats.ox.ac.uk/webapps/ngsdb/json/Huang_2016/Huang_2016_light_2015_2015_3rd_run_light_2-fq-gz_iglblastn.json.gz
wget http://opig.stats.ox.ac.uk/webapps/ngsdb/json/Huang_2016/Huang_2016_heavy_2012_Bulk_2012_3rd_run_heavy_2-fq-gz_iglblastn_Bulk.json.gz
wget http://opig.stats.ox.ac.uk/webapps/ngsdb/json/Huang_2016/Huang_2016_light_2015_2015_1st_run_light_1-fq-gz_iglblastn.json.gz
wget http://opig.stats.ox.ac.uk/webapps/ngsdb/json/Huang_2016/Huang_2016_heavy_2012_Bulk_2012_2nd_run_heavy_1-fq-gz_iglblastn_Bulk.json.gz
wget http://opig.stats.ox.ac.uk/webapps/ngsdb/json/Huang_2016/Huang_2016_heavy_2014_Bulk_2014_3rd_run_heavy_2-fq-gz_iglblastn_Bulk.json.gz
wget http://opig.stats.ox.ac.uk/webapps/ngsdb/json/Huang_2016/Huang_2016_heavy_2015_Bulk_2015_2nd_run_heavy_2-fq-gz_iglblastn_Bulk.json.gz
wget http://opig.stats.ox.ac.uk/webapps/ngsdb/json/Huang_2016/Huang_2016_light_2012_2012_3rd_run_light_2-fq-gz_iglblastn.json.gz
wget http://opig.stats.ox.ac.uk/webapps/ngsdb/json/Huang_2016/Huang_2016_light_2012_2012_2nd_run_light_1-fq-gz_iglblastn.json.gz
wget http://opig.stats.ox.ac.uk/webapps/ngsdb/json/Huang_2016/Huang_2016_light_2015_2015_2nd_run_light_1-fq-gz_iglblastn.json.gz
wget http://opig.stats.ox.ac.uk/webapps/ngsdb/json/Huang_2016/Huang_2016_light_2014_2014_1st_run_light_1-fq-gz_iglblastn.json.gz
wget http://opig.stats.ox.ac.uk/webapps/ngsdb/json/Huang_2016/Huang_2016_light_2014_2014_3rd_run_light_2-fq-gz_iglblastn.json.gz
wget http://opig.stats.ox.ac.uk/webapps/ngsdb/json/Setliff_2018a/Setliff_2018a_3-year_CAP301_3-year_CAP301_No_igblastn.json.gz
wget http://opig.stats.ox.ac.uk/webapps/ngsdb/json/Setliff_2018a/Setliff_2018a_6-month_CAP322_6-month_CAP322_Replicate_igblastn.json.gz
wget http://opig.stats.ox.ac.uk/webapps/ngsdb/json/Setliff_2018a/Setliff_2018a_6-month_CAP322_6-month_CAP322_No_igblastn.json.gz
wget http://opig.stats.ox.ac.uk/webapps/ngsdb/json/Setliff_2018a/Setliff_2018a_3-year_CAP351_3-year_CAP351_Replicate_igblastn.json.gz
wget http://opig.stats.ox.ac.uk/webapps/ngsdb/json/Setliff_2018a/Setliff_2018a_3-year_CAP335_3-year_CAP335_No_igblastn.json.gz
wget http://opig.stats.ox.ac.uk/webapps/ngsdb/json/Setliff_2018a/Setliff_2018a_6-month_CAP351_6-month_CAP351_No_igblastn.json.gz
wget http://opig.stats.ox.ac.uk/webapps/ngsdb/json/Setliff_2018a/Setliff_2018a_3-year_CAP312_3-year_CAP312_No_igblastn.json.gz
wget http://opig.stats.ox.ac.uk/webapps/ngsdb/json/Setliff_2018a/Setliff_2018a_3-year_CAP351_3-year_CAP351_No_igblastn.json.gz
wget http://opig.stats.ox.ac.uk/webapps/ngsdb/json/Setliff_2018a/Setliff_2018a_6-month_CAP287_6-month_CAP287_No_igblastn.json.gz
wget http://opig.stats.ox.ac.uk/webapps/ngsdb/json/Setliff_2018a/Setliff_2018a_6-month_CAP301_6-month_CAP301_No_igblastn.json.gz
wget http://opig.stats.ox.ac.uk/webapps/ngsdb/json/Setliff_2018a/Setliff_2018a_6-month_CAP287_6-month_CAP287_Replicate_igblastn.json.gz
wget http://opig.stats.ox.ac.uk/webapps/ngsdb/json/Setliff_2018a/Setliff_2018a_3-year_CAP287_3-year_CAP287_Replicate_igblastn.json.gz
wget http://opig.stats.ox.ac.uk/webapps/ngsdb/json/Setliff_2018a/Setliff_2018a_3-year_CAP335_3-year_CAP335_Replicate_igblastn.json.gz
wget http://opig.stats.ox.ac.uk/webapps/ngsdb/json/Setliff_2018a/Setliff_2018a_6-month_CAP335_6-month_CAP335_No_igblastn.json.gz
wget http://opig.stats.ox.ac.uk/webapps/ngsdb/json/Setliff_2018a/Setliff_2018a_6-month_CAP312_6-month_CAP312_No_igblastn.json.gz
wget http://opig.stats.ox.ac.uk/webapps/ngsdb/json/Setliff_2018a/Setliff_2018a_3-year_CAP312_3-year_CAP312_Replicate_igblastn.json.gz
wget http://opig.stats.ox.ac.uk/webapps/ngsdb/json/Setliff_2018a/Setliff_2018a_6-month_CAP301_6-month_CAP301_Replicate_igblastn.json.gz
wget http://opig.stats.ox.ac.uk/webapps/ngsdb/json/Setliff_2018a/Setliff_2018a_3-year_CAP322_3-year_CAP322_No_igblastn.json.gz
wget http://opig.stats.ox.ac.uk/webapps/ngsdb/json/Setliff_2018a/Setliff_2018a_6-month_CAP312_6-month_CAP312_Replicate_igblastn.json.gz
wget http://opig.stats.ox.ac.uk/webapps/ngsdb/json/Setliff_2018a/Setliff_2018a_3-year_CAP322_3-year_CAP322_Replicate_igblastn.json.gz
wget http://opig.stats.ox.ac.uk/webapps/ngsdb/json/Setliff_2018a/Setliff_2018a_6-month_CAP351_6-month_CAP351_Replicate_igblastn.json.gz
wget http://opig.stats.ox.ac.uk/webapps/ngsdb/json/Setliff_2018a/Setliff_2018a_3-year_CAP301_3-year_CAP301_Replicate_igblastn.json.gz
wget http://opig.stats.ox.ac.uk/webapps/ngsdb/json/Setliff_2018a/Setliff_2018a_6-month_CAP335_6-month_CAP335_Replicate_igblastn.json.gz
wget http://opig.stats.ox.ac.uk/webapps/ngsdb/json/Setliff_2018a/Setliff_2018a_3-year_CAP287_3-year_CAP287_No_igblastn.json.gz
wget http://opig.stats.ox.ac.uk/webapps/ngsdb/json/Bhiman_2015/Bhiman_2015_Heavy_week-38_IGHA_Heavy_week-38_CAP256_iglblastn_igblastn_IGHA.json.gz
wget http://opig.stats.ox.ac.uk/webapps/ngsdb/json/Bhiman_2015/Bhiman_2015_Heavy_week-38_Bulk_Heavy_week-38_CAP256_iglblastn_igblastn_Bulk.json.gz
wget http://opig.stats.ox.ac.uk/webapps/ngsdb/json/Bhiman_2015/Bhiman_2015_Light_week-38_Light_week-38_CAP256_iglblastn.json.gz
wget http://opig.stats.ox.ac.uk/webapps/ngsdb/json/Bhiman_2015/Bhiman_2015_Heavy_week-38_IGHM_Heavy_week-38_CAP256_iglblastn_igblastn_IGHM.json.gz
wget http://opig.stats.ox.ac.uk/webapps/ngsdb/json/Bhiman_2015/Bhiman_2015_Heavy_week-38_IGHG_Heavy_week-38_CAP256_iglblastn_igblastn_IGHG.json.gz
wget http://opig.stats.ox.ac.uk/webapps/ngsdb/json/Bhiman_2015/Bhiman_2015_Heavy_week-38_IGHE_Heavy_week-38_CAP256_iglblastn_igblastn_IGHE.json.gz
wget http://opig.stats.ox.ac.uk/webapps/ngsdb/json/Zhu_2013/Zhu_2013_Light_Donor_C38_Run9_iglblastn.json.gz
wget http://opig.stats.ox.ac.uk/webapps/ngsdb/json/Zhu_2013/Zhu_2013_Heavy_Bulk_Donor_C38_Run9_iglblastn_igblastn_Bulk.json.gz
wget http://opig.stats.ox.ac.uk/webapps/ngsdb/json/Zhu_2013/Zhu_2013_Light_Donor_C38_Run6_iglblastn.json.gz
wget http://opig.stats.ox.ac.uk/webapps/ngsdb/json/Zhu_2013/Zhu_2013_Heavy_IGHG_Donor_C38_Run9_iglblastn_igblastn_IGHG.json.gz
wget http://opig.stats.ox.ac.uk/webapps/ngsdb/json/Zhu_2013/Zhu_2013_Heavy_Bulk_Donor_C38_Run8_iglblastn_igblastn_Bulk.json.gz
wget http://opig.stats.ox.ac.uk/webapps/ngsdb/json/Zhu_2013/Zhu_2013_Heavy_IGHA_Donor_C38_Run1_iglblastn_igblastn_IGHA.json.gz
wget http://opig.stats.ox.ac.uk/webapps/ngsdb/json/Zhu_2013/Zhu_2013_Heavy_IGHM_Donor_C38_Run4_iglblastn_igblastn_IGHM.json.gz
wget http://opig.stats.ox.ac.uk/webapps/ngsdb/json/Zhu_2013/Zhu_2013_Heavy_IGHG_Donor_C38_Run3_iglblastn_igblastn_IGHG.json.gz
wget http://opig.stats.ox.ac.uk/webapps/ngsdb/json/Zhu_2013/Zhu_2013_Heavy_IGHM_Donor_C38_Run7_iglblastn_igblastn_IGHM.json.gz
wget http://opig.stats.ox.ac.uk/webapps/ngsdb/json/Zhu_2013/Zhu_2013_Heavy_IGHM_Donor_C38_Run1_iglblastn_igblastn_IGHM.json.gz
wget http://opig.stats.ox.ac.uk/webapps/ngsdb/json/Zhu_2013/Zhu_2013_Heavy_IGHM_Donor_C38_Run8_iglblastn_igblastn_IGHM.json.gz
wget http://opig.stats.ox.ac.uk/webapps/ngsdb/json/Zhu_2013/Zhu_2013_Heavy_IGHM_Donor_C38_Run6_iglblastn_igblastn_IGHM.json.gz
wget http://opig.stats.ox.ac.uk/webapps/ngsdb/json/Zhu_2013/Zhu_2013_Heavy_IGHE_Donor_C38_Run2_iglblastn_igblastn_IGHE.json.gz
wget http://opig.stats.ox.ac.uk/webapps/ngsdb/json/Zhu_2013/Zhu_2013_Heavy_IGHE_Donor_C38_Run1_iglblastn_igblastn_IGHE.json.gz
wget http://opig.stats.ox.ac.uk/webapps/ngsdb/json/Zhu_2013/Zhu_2013_Heavy_IGHG_Donor_C38_Run2_iglblastn_igblastn_IGHG.json.gz
wget http://opig.stats.ox.ac.uk/webapps/ngsdb/json/Zhu_2013/Zhu_2013_Heavy_IGHG_Donor_C38_Run4_iglblastn_igblastn_IGHG.json.gz
wget http://opig.stats.ox.ac.uk/webapps/ngsdb/json/Zhu_2013/Zhu_2013_Heavy_IGHG_Donor_C38_Run7_iglblastn_igblastn_IGHG.json.gz
wget http://opig.stats.ox.ac.uk/webapps/ngsdb/json/Zhu_2013/Zhu_2013_Heavy_IGHE_Donor_C38_Run7_iglblastn_igblastn_IGHE.json.gz
wget http://opig.stats.ox.ac.uk/webapps/ngsdb/json/Zhu_2013/Zhu_2013_Heavy_Bulk_Donor_C38_Run1_iglblastn_igblastn_Bulk.json.gz
wget http://opig.stats.ox.ac.uk/webapps/ngsdb/json/Zhu_2013/Zhu_2013_Heavy_Bulk_Donor_C38_Run7_iglblastn_igblastn_Bulk.json.gz
wget http://opig.stats.ox.ac.uk/webapps/ngsdb/json/Zhu_2013/Zhu_2013_Heavy_IGHM_Donor_C38_Run9_iglblastn_igblastn_IGHM.json.gz
wget http://opig.stats.ox.ac.uk/webapps/ngsdb/json/Zhu_2013/Zhu_2013_Light_Donor_C38_Run4_iglblastn.json.gz
wget http://opig.stats.ox.ac.uk/webapps/ngsdb/json/Zhu_2013/Zhu_2013_Light_Donor_C38_Run7_iglblastn.json.gz
wget http://opig.stats.ox.ac.uk/webapps/ngsdb/json/Zhu_2013/Zhu_2013_Heavy_IGHE_Donor_C38_Run8_iglblastn_igblastn_IGHE.json.gz
wget http://opig.stats.ox.ac.uk/webapps/ngsdb/json/Zhu_2013/Zhu_2013_Heavy_IGHG_Donor_C38_Run1_iglblastn_igblastn_IGHG.json.gz
wget http://opig.stats.ox.ac.uk/webapps/ngsdb/json/Zhu_2013/Zhu_2013_Heavy_IGHM_Donor_C38_Run2_iglblastn_igblastn_IGHM.json.gz
wget http://opig.stats.ox.ac.uk/webapps/ngsdb/json/Zhu_2013/Zhu_2013_Light_Donor_C38_Run5_iglblastn.json.gz
wget http://opig.stats.ox.ac.uk/webapps/ngsdb/json/Zhu_2013/Zhu_2013_Heavy_IGHM_Donor_C38_Run3_iglblastn_igblastn_IGHM.json.gz
wget http://opig.stats.ox.ac.uk/webapps/ngsdb/json/Zhu_2013/Zhu_2013_Heavy_IGHA_Donor_C38_Run3_iglblastn_igblastn_IGHA.json.gz
wget http://opig.stats.ox.ac.uk/webapps/ngsdb/json/Zhu_2013/Zhu_2013_Heavy_IGHG_Donor_C38_Run8_iglblastn_igblastn_IGHG.json.gz
wget http://opig.stats.ox.ac.uk/webapps/ngsdb/json/Zhu_2013/Zhu_2013_Heavy_IGHE_Donor_C38_Run3_iglblastn_igblastn_IGHE.json.gz
wget http://opig.stats.ox.ac.uk/webapps/ngsdb/json/Zhu_2013/Zhu_2013_Heavy_Bulk_Donor_C38_Run2_iglblastn_igblastn_Bulk.json.gz
wget http://opig.stats.ox.ac.uk/webapps/ngsdb/json/Zhu_2013/Zhu_2013_Heavy_Bulk_Donor_C38_Run3_iglblastn_igblastn_Bulk.json.gz
wget http://opig.stats.ox.ac.uk/webapps/ngsdb/json/Zhu_2013/Zhu_2013_Heavy_IGHG_Donor_C38_Run6_iglblastn_igblastn_IGHG.json.gz
wget http://opig.stats.ox.ac.uk/webapps/ngsdb/json/Zhu_2012/Zhu_2012_Heavy_IGHE_Donor_39_heavy_chain_iglblastn_IGHE.json.gz
wget http://opig.stats.ox.ac.uk/webapps/ngsdb/json/Zhu_2012/Zhu_2012_Heavy_Bulk_Donor_39_heavy_chain_iglblastn_Bulk.json.gz
wget http://opig.stats.ox.ac.uk/webapps/ngsdb/json/Zhu_2012/Zhu_2012_Heavy_IGHM_Donor_39_heavy_chain_iglblastn_IGHM.json.gz
wget http://opig.stats.ox.ac.uk/webapps/ngsdb/json/Zhu_2012/Zhu_2012_Light_Donor_39_light_chain_iglblastn.json.gz
wget http://opig.stats.ox.ac.uk/webapps/ngsdb/json/Zhu_2012/Zhu_2012_Heavy_IGHG_Donor_39_heavy_chain_iglblastn_IGHG.json.gz
wget http://opig.stats.ox.ac.uk/webapps/ngsdb/json/Zhu_2012/Zhu_2012_Heavy_IGHD_Donor_39_heavy_chain_iglblastn_IGHD.json.gz
wget http://opig.stats.ox.ac.uk/webapps/ngsdb/json/Doria-Rose_2015/Doria-Rose_2015_Heavy_Week38_IGHG_CAP256-Week38-Heavy_iglblastn_IGHG.json.gz
wget http://opig.stats.ox.ac.uk/webapps/ngsdb/json/Doria-Rose_2015/Doria-Rose_2015_Heavy_Week176_IGHE_CAP256-Week176-Heavy_iglblastn_IGHE.json.gz
wget http://opig.stats.ox.ac.uk/webapps/ngsdb/json/Doria-Rose_2015/Doria-Rose_2015_Heavy_Week38_IGHM_CAP256-Week38-Heavy_iglblastn_IGHM.json.gz
wget http://opig.stats.ox.ac.uk/webapps/ngsdb/json/Doria-Rose_2015/Doria-Rose_2015_Heavy_Week176_Bulk_CAP256-Week176-Heavy_iglblastn_Bulk.json.gz
wget http://opig.stats.ox.ac.uk/webapps/ngsdb/json/Doria-Rose_2015/Doria-Rose_2015_Heavy_Week38_Bulk_CAP256-Week38-Heavy_iglblastn_Bulk.json.gz
wget http://opig.stats.ox.ac.uk/webapps/ngsdb/json/Doria-Rose_2015/Doria-Rose_2015_Heavy_Week176_IGHG_CAP256-Week176-Heavy_iglblastn_IGHG.json.gz
wget http://opig.stats.ox.ac.uk/webapps/ngsdb/json/Doria-Rose_2015/Doria-Rose_2015_Heavy_Week206_IGHA_CAP256-Week206-Heavy_iglblastn_IGHA.json.gz
wget http://opig.stats.ox.ac.uk/webapps/ngsdb/json/Doria-Rose_2015/Doria-Rose_2015_Heavy_Week206_IGHG_CAP256-Week206-Heavy_iglblastn_IGHG.json.gz
wget http://opig.stats.ox.ac.uk/webapps/ngsdb/json/Doria-Rose_2015/Doria-Rose_2015_Heavy_Week30_Bulk_CAP256-Week30-Heavy_iglblastn_Bulk.json.gz
wget http://opig.stats.ox.ac.uk/webapps/ngsdb/json/Doria-Rose_2015/Doria-Rose_2015_Heavy_Week119_IGHM_CAP256-Week119-Heavy_iglblastn_IGHM.json.gz
wget http://opig.stats.ox.ac.uk/webapps/ngsdb/json/Doria-Rose_2015/Doria-Rose_2015_Heavy_Week119_IGHG_CAP256-Week119-Heavy_iglblastn_IGHG.json.gz
wget http://opig.stats.ox.ac.uk/webapps/ngsdb/json/Doria-Rose_2015/Doria-Rose_2015_Light_Week16_CAP256-Week16-Light_iglblastn.json.gz
wget http://opig.stats.ox.ac.uk/webapps/ngsdb/json/Doria-Rose_2015/Doria-Rose_2015_Heavy_Week16_IGHG_CAP256-Week16-Heavy_iglblastn_IGHG.json.gz
wget http://opig.stats.ox.ac.uk/webapps/ngsdb/json/Doria-Rose_2015/Doria-Rose_2015_Heavy_Week30_IGHM_CAP256-Week30-Heavy_iglblastn_IGHM.json.gz
wget http://opig.stats.ox.ac.uk/webapps/ngsdb/json/Doria-Rose_2015/Doria-Rose_2015_Heavy_Week176_IGHD_CAP256-Week176-Heavy_iglblastn_IGHD.json.gz
wget http://opig.stats.ox.ac.uk/webapps/ngsdb/json/Doria-Rose_2015/Doria-Rose_2015_Heavy_Week59_IGHM_CAP256-Week59-Heavy_iglblastn_IGHM.json.gz
wget http://opig.stats.ox.ac.uk/webapps/ngsdb/json/Doria-Rose_2015/Doria-Rose_2015_Heavy_Week38_IGHE_CAP256-Week38-Heavy_iglblastn_IGHE.json.gz
wget http://opig.stats.ox.ac.uk/webapps/ngsdb/json/Doria-Rose_2015/Doria-Rose_2015_Heavy_Week48_IGHE_CAP256-Week48-Heavy_iglblastn_IGHE.json.gz
wget http://opig.stats.ox.ac.uk/webapps/ngsdb/json/Doria-Rose_2015/Doria-Rose_2015_Light_Week176_CAP256-Week176-Light_iglblastn.json.gz
wget http://opig.stats.ox.ac.uk/webapps/ngsdb/json/Doria-Rose_2015/Doria-Rose_2015_Heavy_Week16_IGHE_CAP256-Week16-Heavy_iglblastn_IGHE.json.gz
wget http://opig.stats.ox.ac.uk/webapps/ngsdb/json/Doria-Rose_2015/Doria-Rose_2015_Heavy_Week59_Bulk_CAP256-Week59-Heavy_iglblastn_Bulk.json.gz
wget http://opig.stats.ox.ac.uk/webapps/ngsdb/json/Doria-Rose_2015/Doria-Rose_2015_Heavy_Week48_IGHM_CAP256-Week48-Heavy_iglblastn_IGHM.json.gz
wget http://opig.stats.ox.ac.uk/webapps/ngsdb/json/Doria-Rose_2015/Doria-Rose_2015_Heavy_Week206_Bulk_CAP256-Week206-Heavy_iglblastn_Bulk.json.gz
wget http://opig.stats.ox.ac.uk/webapps/ngsdb/json/Doria-Rose_2015/Doria-Rose_2015_Light_Week38_CAP256-Week38-Light_iglblastn.json.gz
wget http://opig.stats.ox.ac.uk/webapps/ngsdb/json/Doria-Rose_2015/Doria-Rose_2015_Heavy_Week59_IGHE_CAP256-Week59-Heavy_iglblastn_IGHE.json.gz
wget http://opig.stats.ox.ac.uk/webapps/ngsdb/json/Doria-Rose_2015/Doria-Rose_2015_Light_Week59_CAP256-Week59-Light_iglblastn.json.gz
wget http://opig.stats.ox.ac.uk/webapps/ngsdb/json/Doria-Rose_2015/Doria-Rose_2015_Heavy_Week48_Bulk_CAP256-Week48-Heavy_iglblastn_Bulk.json.gz
wget http://opig.stats.ox.ac.uk/webapps/ngsdb/json/Doria-Rose_2015/Doria-Rose_2015_Heavy_Week30_IGHG_CAP256-Week30-Heavy_iglblastn_IGHG.json.gz
wget http://opig.stats.ox.ac.uk/webapps/ngsdb/json/Doria-Rose_2015/Doria-Rose_2015_Light_Week30_CAP256-Week30-Light_iglblastn.json.gz
wget http://opig.stats.ox.ac.uk/webapps/ngsdb/json/Doria-Rose_2015/Doria-Rose_2015_Heavy_Week119_IGHE_CAP256-Week119-Heavy_iglblastn_IGHE.json.gz
wget http://opig.stats.ox.ac.uk/webapps/ngsdb/json/Doria-Rose_2015/Doria-Rose_2015_Light_Week48_CAP256-Week48-Light_iglblastn.json.gz
wget http://opig.stats.ox.ac.uk/webapps/ngsdb/json/Doria-Rose_2015/Doria-Rose_2015_Heavy_Week59_IGHA_CAP256-Week59-Heavy_iglblastn_IGHA.json.gz
wget http://opig.stats.ox.ac.uk/webapps/ngsdb/json/Doria-Rose_2015/Doria-Rose_2015_Heavy_Week48_IGHA_CAP256-Week48-Heavy_iglblastn_IGHA.json.gz
wget http://opig.stats.ox.ac.uk/webapps/ngsdb/json/Doria-Rose_2015/Doria-Rose_2015_Heavy_Week30_IGHA_CAP256-Week30-Heavy_iglblastn_IGHA.json.gz
wget http://opig.stats.ox.ac.uk/webapps/ngsdb/json/Doria-Rose_2015/Doria-Rose_2015_Heavy_Week38_IGHA_CAP256-Week38-Heavy_iglblastn_IGHA.json.gz
wget http://opig.stats.ox.ac.uk/webapps/ngsdb/json/Doria-Rose_2015/Doria-Rose_2015_Heavy_Week16_Bulk_CAP256-Week16-Heavy_iglblastn_Bulk.json.gz
wget http://opig.stats.ox.ac.uk/webapps/ngsdb/json/Doria-Rose_2015/Doria-Rose_2015_Heavy_Week176_IGHM_CAP256-Week176-Heavy_iglblastn_IGHM.json.gz
wget http://opig.stats.ox.ac.uk/webapps/ngsdb/json/Doria-Rose_2015/Doria-Rose_2015_Heavy_Week206_IGHE_CAP256-Week206-Heavy_iglblastn_IGHE.json.gz
wget http://opig.stats.ox.ac.uk/webapps/ngsdb/json/Doria-Rose_2015/Doria-Rose_2015_Heavy_Week16_IGHA_CAP256-Week16-Heavy_iglblastn_IGHA.json.gz
wget http://opig.stats.ox.ac.uk/webapps/ngsdb/json/Doria-Rose_2015/Doria-Rose_2015_Heavy_Week206_IGHM_CAP256-Week206-Heavy_iglblastn_IGHM.json.gz
wget http://opig.stats.ox.ac.uk/webapps/ngsdb/json/Doria-Rose_2015/Doria-Rose_2015_Heavy_Week119_Bulk_CAP256-Week119-Heavy_iglblastn_Bulk.json.gz
wget http://opig.stats.ox.ac.uk/webapps/ngsdb/json/Doria-Rose_2015/Doria-Rose_2015_Light_Week206_CAP256-Week206-Light_iglblastn.json.gz
wget http://opig.stats.ox.ac.uk/webapps/ngsdb/json/Doria-Rose_2015/Doria-Rose_2015_Heavy_Week30_IGHE_CAP256-Week30-Heavy_iglblastn_IGHE.json.gz
wget http://opig.stats.ox.ac.uk/webapps/ngsdb/json/Doria-Rose_2015/Doria-Rose_2015_Heavy_Week16_IGHM_CAP256-Week16-Heavy_iglblastn_IGHM.json.gz
wget http://opig.stats.ox.ac.uk/webapps/ngsdb/json/Doria-Rose_2015/Doria-Rose_2015_Heavy_Week48_IGHG_CAP256-Week48-Heavy_iglblastn_IGHG.json.gz
wget http://opig.stats.ox.ac.uk/webapps/ngsdb/json/Doria-Rose_2015/Doria-Rose_2015_Light_Week119_CAP256-Week119-Light_iglblastn.json.gz
wget http://opig.stats.ox.ac.uk/webapps/ngsdb/json/Doria-Rose_2015/Doria-Rose_2015_Heavy_Week59_IGHG_CAP256-Week59-Heavy_iglblastn_IGHG.json.gz
wget http://opig.stats.ox.ac.uk/webapps/ngsdb/json/Bonsignori_2016/Bonsignori_2016_Hiv_infected_week-100_IGHE_HIV_infected_week-100_week_repeat-6_PBMC_iglblastn_igblastn_IGHE.json.gz
wget http://opig.stats.ox.ac.uk/webapps/ngsdb/json/Bonsignori_2016/Bonsignori_2016_Hiv_infected_week-30_Bulk_HIV_infected_week-30_week_repeat-2_PBMC_iglblastn_igblastn_Bulk.json.gz
wget http://opig.stats.ox.ac.uk/webapps/ngsdb/json/Bonsignori_2016/Bonsignori_2016_Hiv_infected_week-9_Bulk_HIV_infected_week-9_week_repeat-1_PBMC_iglblastn_igblastn_Bulk.json.gz
wget http://opig.stats.ox.ac.uk/webapps/ngsdb/json/Bonsignori_2016/Bonsignori_2016_Hiv_infected_week-22_Bulk_HIV_infected_week-22_week_repeat-2_PBMC_iglblastn_igblastn_Bulk.json.gz
wget http://opig.stats.ox.ac.uk/webapps/ngsdb/json/Bonsignori_2016/Bonsignori_2016_Hiv_infected_week-7_Bulk_HIV_infected_week-7_week_repeat-1_PBMC_iglblastn_igblastn_Bulk.json.gz
wget http://opig.stats.ox.ac.uk/webapps/ngsdb/json/Bonsignori_2016/Bonsignori_2016_Hiv_infected_week-9_Bulk_HIV_infected_week-9_week_repeat-5_PBMC_iglblastn_igblastn_Bulk.json.gz
wget http://opig.stats.ox.ac.uk/webapps/ngsdb/json/Bonsignori_2016/Bonsignori_2016_Hiv_infected_week-100_IGHG_HIV_infected_week-100_week_repeat-6_PBMC_iglblastn_igblastn_IGHG.json.gz
wget http://opig.stats.ox.ac.uk/webapps/ngsdb/json/Bonsignori_2016/Bonsignori_2016_Hiv_infected_week-41_IGHG_HIV_infected_week-41_week_repeat-6_PBMC_iglblastn_igblastn_IGHG.json.gz
wget http://opig.stats.ox.ac.uk/webapps/ngsdb/json/Bonsignori_2016/Bonsignori_2016_Hiv_infected_week-323_Bulk_HIV_infected_week-323_week_repeat-6_PBMC_iglblastn_igblastn_Bulk.json.gz
wget http://opig.stats.ox.ac.uk/webapps/ngsdb/json/Bonsignori_2016/Bonsignori_2016_Hiv_infected_week-20_IGHG_HIV_infected_week-20_week_repeat-6_PBMC_iglblastn_igblastn_IGHG.json.gz
wget http://opig.stats.ox.ac.uk/webapps/ngsdb/json/Bonsignori_2016/Bonsignori_2016_Hiv_infected_week-53_IGHE_HIV_infected_week-53_week_repeat-5_PBMC_iglblastn_igblastn_IGHE.json.gz
wget http://opig.stats.ox.ac.uk/webapps/ngsdb/json/Bonsignori_2016/Bonsignori_2016_Hiv_infected_week-8_Bulk_HIV_infected_week-8_week_repeat-5_PBMC_iglblastn_igblastn_Bulk.json.gz
wget http://opig.stats.ox.ac.uk/webapps/ngsdb/json/Bonsignori_2016/Bonsignori_2016_Hiv_infected_week-264_IGHE_HIV_infected_week-264_week_repeat-5_PBMC_iglblastn_igblastn_IGHE.json.gz
wget http://opig.stats.ox.ac.uk/webapps/ngsdb/json/Bonsignori_2016/Bonsignori_2016_Hiv_infected_week-9_IGHG_HIV_infected_week-9_week_repeat-5_PBMC_iglblastn_igblastn_IGHG.json.gz
wget http://opig.stats.ox.ac.uk/webapps/ngsdb/json/Bonsignori_2016/Bonsignori_2016_Hiv_infected_week-22_Bulk_HIV_infected_week-22_week_repeat-5_PBMC_iglblastn_igblastn_Bulk.json.gz
wget http://opig.stats.ox.ac.uk/webapps/ngsdb/json/Bonsignori_2016/Bonsignori_2016_Hiv_infected_week-6_Bulk_HIV_infected_week-6_week_repeat-1_PBMC_iglblastn_igblastn_Bulk.json.gz
wget http://opig.stats.ox.ac.uk/webapps/ngsdb/json/Bonsignori_2016/Bonsignori_2016_Hiv_infected_week-6_Bulk_HIV_infected_week-6_week_repeat-6_PBMC_iglblastn_igblastn_Bulk.json.gz
wget http://opig.stats.ox.ac.uk/webapps/ngsdb/json/Bonsignori_2016/Bonsignori_2016_Hiv_infected_week-6_Bulk_HIV_infected_week-6_week_repeat-5_PBMC_iglblastn_igblastn_Bulk.json.gz
wget http://opig.stats.ox.ac.uk/webapps/ngsdb/json/Bonsignori_2016/Bonsignori_2016_Hiv_infected_week-66_Bulk_HIV_infected_week-66_week_repeat-5_PBMC_iglblastn_igblastn_Bulk.json.gz
wget http://opig.stats.ox.ac.uk/webapps/ngsdb/json/Bonsignori_2016/Bonsignori_2016_Hiv_infected_week-152_IGHG_HIV_infected_week-152_week_repeat-6_PBMC_iglblastn_igblastn_IGHG.json.gz
wget http://opig.stats.ox.ac.uk/webapps/ngsdb/json/Bonsignori_2016/Bonsignori_2016_Hiv_infected_week-144_Bulk_HIV_infected_week-144_week_repeat-5_PBMC_iglblastn_igblastn_Bulk.json.gz
wget http://opig.stats.ox.ac.uk/webapps/ngsdb/json/Bonsignori_2016/Bonsignori_2016_Hiv_infected_week-323_Bulk_HIV_infected_week-323_week_repeat-2_PBMC_iglblastn_igblastn_Bulk.json.gz
wget http://opig.stats.ox.ac.uk/webapps/ngsdb/json/Bonsignori_2016/Bonsignori_2016_Hiv_infected_week-92_IGHG_HIV_infected_week-92_week_repeat-5_PBMC_iglblastn_igblastn_IGHG.json.gz
wget http://opig.stats.ox.ac.uk/webapps/ngsdb/json/Bonsignori_2016/Bonsignori_2016_Hiv_infected_week-264_Bulk_HIV_infected_week-264_week_repeat-6_PBMC_iglblastn_igblastn_Bulk.json.gz
wget http://opig.stats.ox.ac.uk/webapps/ngsdb/json/Bonsignori_2016/Bonsignori_2016_Hiv_infected_week-152_Bulk_HIV_infected_week-152_week_repeat-5_PBMC_iglblastn_igblastn_Bulk.json.gz
wget http://opig.stats.ox.ac.uk/webapps/ngsdb/json/Bonsignori_2016/Bonsignori_2016_Hiv_infected_week-144_Bulk_HIV_infected_week-144_week_repeat-1_PBMC_iglblastn_igblastn_Bulk.json.gz
wget http://opig.stats.ox.ac.uk/webapps/ngsdb/json/Bonsignori_2016/Bonsignori_2016_Hiv_infected_week-323_IGHG_HIV_infected_week-323_week_repeat-7_PBMC_iglblastn_igblastn_IGHG.json.gz
wget http://opig.stats.ox.ac.uk/webapps/ngsdb/json/Bonsignori_2016/Bonsignori_2016_Hiv_infected_week-100_Bulk_HIV_infected_week-100_week_repeat-2_PBMC_iglblastn_igblastn_Bulk.json.gz
wget http://opig.stats.ox.ac.uk/webapps/ngsdb/json/Bonsignori_2016/Bonsignori_2016_Hiv_infected_week-264_Bulk_HIV_infected_week-264_week_repeat-5_PBMC_iglblastn_igblastn_Bulk.json.gz
wget http://opig.stats.ox.ac.uk/webapps/ngsdb/json/Bonsignori_2016/Bonsignori_2016_Hiv_infected_week-30_IGHE_HIV_infected_week-30_week_repeat-5_PBMC_iglblastn_igblastn_IGHE.json.gz
wget http://opig.stats.ox.ac.uk/webapps/ngsdb/json/Bonsignori_2016/Bonsignori_2016_Hiv_infected_week-66_Bulk_HIV_infected_week-66_week_repeat-2_PBMC_iglblastn_igblastn_Bulk.json.gz
wget http://opig.stats.ox.ac.uk/webapps/ngsdb/json/Bonsignori_2016/Bonsignori_2016_Hiv_infected_week-8_IGHE_HIV_infected_week-8_week_repeat-6_PBMC_iglblastn_igblastn_IGHE.json.gz
wget http://opig.stats.ox.ac.uk/webapps/ngsdb/json/Bonsignori_2016/Bonsignori_2016_Hiv_infected_week-8_IGHE_HIV_infected_week-8_week_repeat-5_PBMC_iglblastn_igblastn_IGHE.json.gz
wget http://opig.stats.ox.ac.uk/webapps/ngsdb/json/Bonsignori_2016/Bonsignori_2016_Hiv_infected_week-323_IGHE_HIV_infected_week-323_week_repeat-6_PBMC_iglblastn_igblastn_IGHE.json.gz
wget http://opig.stats.ox.ac.uk/webapps/ngsdb/json/Bonsignori_2016/Bonsignori_2016_Hiv_infected_week-152_IGHE_HIV_infected_week-152_week_repeat-5_PBMC_iglblastn_igblastn_IGHE.json.gz
wget http://opig.stats.ox.ac.uk/webapps/ngsdb/json/Bonsignori_2016/Bonsignori_2016_Hiv_infected_week-22_Bulk_HIV_infected_week-22_week_repeat-1_PBMC_iglblastn_igblastn_Bulk.json.gz
wget http://opig.stats.ox.ac.uk/webapps/ngsdb/json/Bonsignori_2016/Bonsignori_2016_Hiv_infected_week-7_Bulk_HIV_infected_week-7_week_repeat-2_PBMC_iglblastn_igblastn_Bulk.json.gz
wget http://opig.stats.ox.ac.uk/webapps/ngsdb/json/Bonsignori_2016/Bonsignori_2016_Hiv_infected_week-66_Bulk_HIV_infected_week-66_week_repeat-6_PBMC_iglblastn_igblastn_Bulk.json.gz
wget http://opig.stats.ox.ac.uk/webapps/ngsdb/json/Bonsignori_2016/Bonsignori_2016_Hiv_infected_week-30_Bulk_HIV_infected_week-30_week_repeat-1_PBMC_iglblastn_igblastn_Bulk.json.gz
wget http://opig.stats.ox.ac.uk/webapps/ngsdb/json/Bonsignori_2016/Bonsignori_2016_Hiv_infected_week-22_Bulk_HIV_infected_week-22_week_repeat-6_PBMC_iglblastn_igblastn_Bulk.json.gz
wget http://opig.stats.ox.ac.uk/webapps/ngsdb/json/Bonsignori_2016/Bonsignori_2016_Hiv_infected_week-100_Bulk_HIV_infected_week-100_week_repeat-6_PBMC_iglblastn_igblastn_Bulk.json.gz
wget http://opig.stats.ox.ac.uk/webapps/ngsdb/json/Bonsignori_2016/Bonsignori_2016_Hiv_infected_week-152_Bulk_HIV_infected_week-152_week_repeat-1_PBMC_iglblastn_igblastn_Bulk.json.gz
wget http://opig.stats.ox.ac.uk/webapps/ngsdb/json/Bonsignori_2016/Bonsignori_2016_Hiv_infected_week-323_Bulk_HIV_infected_week-323_week_repeat-3_PBMC_iglblastn_igblastn_Bulk.json.gz
wget http://opig.stats.ox.ac.uk/webapps/ngsdb/json/Bonsignori_2016/Bonsignori_2016_Hiv_infected_week-144_IGHE_HIV_infected_week-144_week_repeat-6_PBMC_iglblastn_igblastn_IGHE.json.gz
wget http://opig.stats.ox.ac.uk/webapps/ngsdb/json/Bonsignori_2016/Bonsignori_2016_Hiv_infected_week-8_IGHG_HIV_infected_week-8_week_repeat-5_PBMC_iglblastn_igblastn_IGHG.json.gz
wget http://opig.stats.ox.ac.uk/webapps/ngsdb/json/Bonsignori_2016/Bonsignori_2016_Hiv_infected_week-264_Bulk_HIV_infected_week-264_week_repeat-2_PBMC_iglblastn_igblastn_Bulk.json.gz
wget http://opig.stats.ox.ac.uk/webapps/ngsdb/json/Bonsignori_2016/Bonsignori_2016_Hiv_infected_week-9_Bulk_HIV_infected_week-9_week_repeat-6_PBMC_iglblastn_igblastn_Bulk.json.gz
wget http://opig.stats.ox.ac.uk/webapps/ngsdb/json/Bonsignori_2016/Bonsignori_2016_Hiv_infected_week-144_Bulk_HIV_infected_week-144_week_repeat-6_PBMC_iglblastn_igblastn_Bulk.json.gz
wget http://opig.stats.ox.ac.uk/webapps/ngsdb/json/Bonsignori_2016/Bonsignori_2016_Hiv_infected_week-30_IGHG_HIV_infected_week-30_week_repeat-5_PBMC_iglblastn_igblastn_IGHG.json.gz
wget http://opig.stats.ox.ac.uk/webapps/ngsdb/json/Bonsignori_2016/Bonsignori_2016_Hiv_infected_week-264_Bulk_HIV_infected_week-264_week_repeat-1_PBMC_iglblastn_igblastn_Bulk.json.gz
wget http://opig.stats.ox.ac.uk/webapps/ngsdb/json/Bonsignori_2016/Bonsignori_2016_Hiv_infected_week-323_IGHG_HIV_infected_week-323_week_repeat-6_PBMC_iglblastn_igblastn_IGHG.json.gz
wget http://opig.stats.ox.ac.uk/webapps/ngsdb/json/Bonsignori_2016/Bonsignori_2016_Hiv_infected_week-92_Bulk_HIV_infected_week-92_week_repeat-1_PBMC_iglblastn_igblastn_Bulk.json.gz
wget http://opig.stats.ox.ac.uk/webapps/ngsdb/json/Bonsignori_2016/Bonsignori_2016_Hiv_infected_week-9_IGHE_HIV_infected_week-9_week_repeat-5_PBMC_iglblastn_igblastn_IGHE.json.gz
wget http://opig.stats.ox.ac.uk/webapps/ngsdb/json/Bonsignori_2016/Bonsignori_2016_Hiv_infected_week-41_IGHE_HIV_infected_week-41_week_repeat-5_PBMC_iglblastn_igblastn_IGHE.json.gz
wget http://opig.stats.ox.ac.uk/webapps/ngsdb/json/Bonsignori_2016/Bonsignori_2016_Hiv_infected_week-8_IGHG_HIV_infected_week-8_week_repeat-6_PBMC_iglblastn_igblastn_IGHG.json.gz
wget http://opig.stats.ox.ac.uk/webapps/ngsdb/json/Bonsignori_2016/Bonsignori_2016_Hiv_infected_week-53_Bulk_HIV_infected_week-53_week_repeat-2_PBMC_iglblastn_igblastn_Bulk.json.gz
wget http://opig.stats.ox.ac.uk/webapps/ngsdb/json/Bonsignori_2016/Bonsignori_2016_Hiv_infected_week-100_IGHE_HIV_infected_week-100_week_repeat-5_PBMC_iglblastn_igblastn_IGHE.json.gz
wget http://opig.stats.ox.ac.uk/webapps/ngsdb/json/Bonsignori_2016/Bonsignori_2016_Hiv_infected_week-53_Bulk_HIV_infected_week-53_week_repeat-6_PBMC_iglblastn_igblastn_Bulk.json.gz
wget http://opig.stats.ox.ac.uk/webapps/ngsdb/json/Bonsignori_2016/Bonsignori_2016_Hiv_infected_week-144_Bulk_HIV_infected_week-144_week_repeat-2_PBMC_iglblastn_igblastn_Bulk.json.gz
wget http://opig.stats.ox.ac.uk/webapps/ngsdb/json/Bonsignori_2016/Bonsignori_2016_Hiv_infected_week-7_Bulk_HIV_infected_week-7_week_repeat-6_PBMC_iglblastn_igblastn_Bulk.json.gz
wget http://opig.stats.ox.ac.uk/webapps/ngsdb/json/Bonsignori_2016/Bonsignori_2016_Hiv_infected_week-7_Bulk_HIV_infected_week-7_week_repeat-5_PBMC_iglblastn_igblastn_Bulk.json.gz
wget http://opig.stats.ox.ac.uk/webapps/ngsdb/json/Bonsignori_2016/Bonsignori_2016_Hiv_infected_week-41_Bulk_HIV_infected_week-41_week_repeat-6_PBMC_iglblastn_igblastn_Bulk.json.gz
wget http://opig.stats.ox.ac.uk/webapps/ngsdb/json/Bonsignori_2016/Bonsignori_2016_Hiv_infected_week-53_Bulk_HIV_infected_week-53_week_repeat-1_PBMC_iglblastn_igblastn_Bulk.json.gz
wget http://opig.stats.ox.ac.uk/webapps/ngsdb/json/Bonsignori_2016/Bonsignori_2016_Hiv_infected_week-100_Bulk_HIV_infected_week-100_week_repeat-1_PBMC_iglblastn_igblastn_Bulk.json.gz
wget http://opig.stats.ox.ac.uk/webapps/ngsdb/json/Bonsignori_2016/Bonsignori_2016_Hiv_infected_week-66_Bulk_HIV_infected_week-66_week_repeat-1_PBMC_iglblastn_igblastn_Bulk.json.gz
wget http://opig.stats.ox.ac.uk/webapps/ngsdb/json/Bonsignori_2016/Bonsignori_2016_Hiv_infected_week-20_IGHG_HIV_infected_week-20_week_repeat-5_PBMC_iglblastn_igblastn_IGHG.json.gz
wget http://opig.stats.ox.ac.uk/webapps/ngsdb/json/Bonsignori_2016/Bonsignori_2016_Hiv_infected_week-100_Bulk_HIV_infected_week-100_week_repeat-5_PBMC_iglblastn_igblastn_Bulk.json.gz
wget http://opig.stats.ox.ac.uk/webapps/ngsdb/json/Bonsignori_2016/Bonsignori_2016_Hiv_infected_week-152_Bulk_HIV_infected_week-152_week_repeat-6_PBMC_iglblastn_igblastn_Bulk.json.gz
wget http://opig.stats.ox.ac.uk/webapps/ngsdb/json/Bonsignori_2016/Bonsignori_2016_Hiv_infected_week-9_IGHG_HIV_infected_week-9_week_repeat-6_PBMC_iglblastn_igblastn_IGHG.json.gz
wget http://opig.stats.ox.ac.uk/webapps/ngsdb/json/Bonsignori_2016/Bonsignori_2016_Hiv_infected_week-92_Bulk_HIV_infected_week-92_week_repeat-5_PBMC_iglblastn_igblastn_Bulk.json.gz
wget http://opig.stats.ox.ac.uk/webapps/ngsdb/json/Bonsignori_2016/Bonsignori_2016_Hiv_infected_week-9_Bulk_HIV_infected_week-9_week_repeat-2_PBMC_iglblastn_igblastn_Bulk.json.gz
wget http://opig.stats.ox.ac.uk/webapps/ngsdb/json/Bonsignori_2016/Bonsignori_2016_Hiv_infected_week-53_Bulk_HIV_infected_week-53_week_repeat-5_PBMC_iglblastn_igblastn_Bulk.json.gz
wget http://opig.stats.ox.ac.uk/webapps/ngsdb/json/Bonsignori_2016/Bonsignori_2016_Hiv_infected_week-41_Bulk_HIV_infected_week-41_week_repeat-2_PBMC_iglblastn_igblastn_Bulk.json.gz
wget http://opig.stats.ox.ac.uk/webapps/ngsdb/json/Bonsignori_2016/Bonsignori_2016_Hiv_infected_week-41_Bulk_HIV_infected_week-41_week_repeat-5_PBMC_iglblastn_igblastn_Bulk.json.gz
wget http://opig.stats.ox.ac.uk/webapps/ngsdb/json/Bonsignori_2016/Bonsignori_2016_Hiv_infected_week-8_Bulk_HIV_infected_week-8_week_repeat-1_PBMC_iglblastn_igblastn_Bulk.json.gz
wget http://opig.stats.ox.ac.uk/webapps/ngsdb/json/Bonsignori_2016/Bonsignori_2016_Hiv_infected_week-6_Bulk_HIV_infected_week-6_week_repeat-2_PBMC_iglblastn_igblastn_Bulk.json.gz
wget http://opig.stats.ox.ac.uk/webapps/ngsdb/json/Bonsignori_2016/Bonsignori_2016_Hiv_infected_week-53_IGHE_HIV_infected_week-53_week_repeat-6_PBMC_iglblastn_igblastn_IGHE.json.gz
wget http://opig.stats.ox.ac.uk/webapps/ngsdb/json/Bonsignori_2016/Bonsignori_2016_Hiv_infected_week-30_Bulk_HIV_infected_week-30_week_repeat-6_PBMC_iglblastn_igblastn_Bulk.json.gz
wget http://opig.stats.ox.ac.uk/webapps/ngsdb/json/Bonsignori_2016/Bonsignori_2016_Hiv_infected_week-20_Bulk_HIV_infected_week-20_week_repeat-5_PBMC_iglblastn_igblastn_Bulk.json.gz
wget http://opig.stats.ox.ac.uk/webapps/ngsdb/json/Bonsignori_2016/Bonsignori_2016_Hiv_infected_week-152_IGHE_HIV_infected_week-152_week_repeat-6_PBMC_iglblastn_igblastn_IGHE.json.gz
wget http://opig.stats.ox.ac.uk/webapps/ngsdb/json/Bonsignori_2016/Bonsignori_2016_Hiv_infected_week-30_IGHG_HIV_infected_week-30_week_repeat-6_PBMC_iglblastn_igblastn_IGHG.json.gz
wget http://opig.stats.ox.ac.uk/webapps/ngsdb/json/Bonsignori_2016/Bonsignori_2016_Hiv_infected_week-92_Bulk_HIV_infected_week-92_week_repeat-2_PBMC_iglblastn_igblastn_Bulk.json.gz
wget http://opig.stats.ox.ac.uk/webapps/ngsdb/json/Bonsignori_2016/Bonsignori_2016_Hiv_infected_week-30_IGHE_HIV_infected_week-30_week_repeat-6_PBMC_iglblastn_igblastn_IGHE.json.gz
wget http://opig.stats.ox.ac.uk/webapps/ngsdb/json/Bonsignori_2016/Bonsignori_2016_Hiv_infected_week-20_IGHE_HIV_infected_week-20_week_repeat-5_PBMC_iglblastn_igblastn_IGHE.json.gz
wget http://opig.stats.ox.ac.uk/webapps/ngsdb/json/Bonsignori_2016/Bonsignori_2016_Hiv_infected_week-152_Bulk_HIV_infected_week-152_week_repeat-2_PBMC_iglblastn_igblastn_Bulk.json.gz
wget http://opig.stats.ox.ac.uk/webapps/ngsdb/json/Bonsignori_2016/Bonsignori_2016_Hiv_infected_week-20_Bulk_HIV_infected_week-20_week_repeat-2_PBMC_iglblastn_igblastn_Bulk.json.gz
wget http://opig.stats.ox.ac.uk/webapps/ngsdb/json/Bonsignori_2016/Bonsignori_2016_Hiv_infected_week-53_IGHG_HIV_infected_week-53_week_repeat-5_PBMC_iglblastn_igblastn_IGHG.json.gz
wget http://opig.stats.ox.ac.uk/webapps/ngsdb/json/Bonsignori_2016/Bonsignori_2016_Hiv_infected_week-20_Bulk_HIV_infected_week-20_week_repeat-6_PBMC_iglblastn_igblastn_Bulk.json.gz
wget http://opig.stats.ox.ac.uk/webapps/ngsdb/json/Bonsignori_2016/Bonsignori_2016_Hiv_infected_week-152_IGHG_HIV_infected_week-152_week_repeat-5_PBMC_iglblastn_igblastn_IGHG.json.gz
wget http://opig.stats.ox.ac.uk/webapps/ngsdb/json/Bonsignori_2016/Bonsignori_2016_Hiv_infected_week-30_Bulk_HIV_infected_week-30_week_repeat-5_PBMC_iglblastn_igblastn_Bulk.json.gz
wget http://opig.stats.ox.ac.uk/webapps/ngsdb/json/Bonsignori_2016/Bonsignori_2016_Hiv_infected_week-8_Bulk_HIV_infected_week-8_week_repeat-6_PBMC_iglblastn_igblastn_Bulk.json.gz
wget http://opig.stats.ox.ac.uk/webapps/ngsdb/json/Bonsignori_2016/Bonsignori_2016_Hiv_infected_week-7_IGHE_HIV_infected_week-7_week_repeat-5_PBMC_iglblastn_igblastn_IGHE.json.gz
wget http://opig.stats.ox.ac.uk/webapps/ngsdb/json/Bonsignori_2016/Bonsignori_2016_Hiv_infected_week-9_IGHE_HIV_infected_week-9_week_repeat-6_PBMC_iglblastn_igblastn_IGHE.json.gz
wget http://opig.stats.ox.ac.uk/webapps/ngsdb/json/Bonsignori_2016/Bonsignori_2016_Hiv_infected_week-20_IGHE_HIV_infected_week-20_week_repeat-6_PBMC_iglblastn_igblastn_IGHE.json.gz
wget http://opig.stats.ox.ac.uk/webapps/ngsdb/json/Bonsignori_2016/Bonsignori_2016_Hiv_infected_week-8_Bulk_HIV_infected_week-8_week_repeat-2_PBMC_iglblastn_igblastn_Bulk.json.gz
wget http://opig.stats.ox.ac.uk/webapps/ngsdb/json/Bonsignori_2016/Bonsignori_2016_Hiv_infected_week-6_IGHE_HIV_infected_week-6_week_repeat-6_PBMC_iglblastn_igblastn_IGHE.json.gz
wget http://opig.stats.ox.ac.uk/webapps/ngsdb/json/Bonsignori_2016/Bonsignori_2016_Hiv_infected_week-6_IGHG_HIV_infected_week-6_week_repeat-6_PBMC_iglblastn_igblastn_IGHG.json.gz
wget http://opig.stats.ox.ac.uk/webapps/ngsdb/json/Bonsignori_2016/Bonsignori_2016_Hiv_infected_week-53_IGHG_HIV_infected_week-53_week_repeat-6_PBMC_iglblastn_igblastn_IGHG.json.gz
wget http://opig.stats.ox.ac.uk/webapps/ngsdb/json/Bonsignori_2016/Bonsignori_2016_Hiv_infected_week-323_Bulk_HIV_infected_week-323_week_repeat-7_PBMC_iglblastn_igblastn_Bulk.json.gz
wget http://opig.stats.ox.ac.uk/webapps/ngsdb/json/Bonsignori_2016/Bonsignori_2016_Hiv_infected_week-41_Bulk_HIV_infected_week-41_week_repeat-1_PBMC_iglblastn_igblastn_Bulk.json.gz
wget http://opig.stats.ox.ac.uk/webapps/ngsdb/json/Bonsignori_2016/Bonsignori_2016_Hiv_infected_week-20_Bulk_HIV_infected_week-20_week_repeat-1_PBMC_iglblastn_igblastn_Bulk.json.gz
wget http://opig.stats.ox.ac.uk/webapps/ngsdb/json/Soto_2016/Soto_2016_Heavy_IGHA_IAVI84_Heavy_iglblastn_IGHA.json.gz
wget http://opig.stats.ox.ac.uk/webapps/ngsdb/json/Soto_2016/Soto_2016_Heavy_IGHA_Donor_N152_2_iglblastn_IGHA.json.gz
wget http://opig.stats.ox.ac.uk/webapps/ngsdb/json/Soto_2016/Soto_2016_Heavy_IGHD_IAVI84_Heavy_iglblastn_IGHD.json.gz
wget http://opig.stats.ox.ac.uk/webapps/ngsdb/json/Soto_2016/Soto_2016_Heavy_IGHG_Donor_N152_2_iglblastn_IGHG.json.gz
wget http://opig.stats.ox.ac.uk/webapps/ngsdb/json/Soto_2016/Soto_2016_Heavy_IGHD_Donor_N152_2_iglblastn_IGHD.json.gz
wget http://opig.stats.ox.ac.uk/webapps/ngsdb/json/Soto_2016/Soto_2016_Heavy_Bulk_Donor_N152_2_iglblastn_Bulk.json.gz
wget http://opig.stats.ox.ac.uk/webapps/ngsdb/json/Soto_2016/Soto_2016_Heavy_IGHG_Donor_N152_1_iglblastn_IGHG.json.gz
wget http://opig.stats.ox.ac.uk/webapps/ngsdb/json/Soto_2016/Soto_2016_Light_Donor_N152_Light_iglblastn.json.gz
wget http://opig.stats.ox.ac.uk/webapps/ngsdb/json/Soto_2016/Soto_2016_Heavy_IGHE_Donor_N152_1_iglblastn_IGHE.json.gz
wget http://opig.stats.ox.ac.uk/webapps/ngsdb/json/Soto_2016/Soto_2016_Heavy_IGHM_Donor_N152_1_iglblastn_IGHM.json.gz
wget http://opig.stats.ox.ac.uk/webapps/ngsdb/json/Soto_2016/Soto_2016_Heavy_IGHA_Donor_N152_1_iglblastn_IGHA.json.gz
wget http://opig.stats.ox.ac.uk/webapps/ngsdb/json/Soto_2016/Soto_2016_Heavy_IGHM_IAVI84_Heavy_iglblastn_IGHM.json.gz
wget http://opig.stats.ox.ac.uk/webapps/ngsdb/json/Soto_2016/Soto_2016_Heavy_Bulk_Donor_N152_1_iglblastn_Bulk.json.gz
wget http://opig.stats.ox.ac.uk/webapps/ngsdb/json/Soto_2016/Soto_2016_Heavy_IGHE_IAVI84_Heavy_iglblastn_IGHE.json.gz
wget http://opig.stats.ox.ac.uk/webapps/ngsdb/json/Soto_2016/Soto_2016_Heavy_IGHD_Donor_N152_1_iglblastn_IGHD.json.gz
wget http://opig.stats.ox.ac.uk/webapps/ngsdb/json/Soto_2016/Soto_2016_Light_IAVI84_Light_iglblastn.json.gz
wget http://opig.stats.ox.ac.uk/webapps/ngsdb/json/Soto_2016/Soto_2016_Heavy_IGHE_Donor_N152_2_iglblastn_IGHE.json.gz
wget http://opig.stats.ox.ac.uk/webapps/ngsdb/json/Soto_2016/Soto_2016_Heavy_IGHM_Donor_N152_2_iglblastn_IGHM.json.gz
wget http://opig.stats.ox.ac.uk/webapps/ngsdb/json/Soto_2016/Soto_2016_Heavy_Bulk_IAVI84_Heavy_iglblastn_Bulk.json.gz
wget http://opig.stats.ox.ac.uk/webapps/ngsdb/json/Soto_2016/Soto_2016_Heavy_IGHG_IAVI84_Heavy_iglblastn_IGHG.json.gz
wget http://opig.stats.ox.ac.uk/webapps/ngsdb/json/Zhou_2015/Zhou_2015_IGHM_DONORC38-HeavyChain-VRC16_iglblastn_igblastn_IGHM.json.gz
wget http://opig.stats.ox.ac.uk/webapps/ngsdb/json/Zhou_2015/Zhou_2015_IGHG_DONORC38-HeavyChain-VRC16_iglblastn_igblastn_IGHG.json.gz
wget http://opig.stats.ox.ac.uk/webapps/ngsdb/json/Zhou_2015/Zhou_2015_IGHM_DONORRU08-HeavyChain_iglblastn_igblastn_IGHM.json.gz
wget http://opig.stats.ox.ac.uk/webapps/ngsdb/json/Zhou_2015/Zhou_2015_IGHD_DONORRU01-HeavyChain_iglblastn_igblastn_IGHD.json.gz
wget http://opig.stats.ox.ac.uk/webapps/ngsdb/json/Zhou_2015/Zhou_2015_IGHG_DONOR44-HeavyChain_iglblastn_igblastn_IGHG.json.gz
wget http://opig.stats.ox.ac.uk/webapps/ngsdb/json/Zhou_2015/Zhou_2015_Bulk_DONORRU08-HeavyChain_iglblastn_igblastn_Bulk.json.gz
wget http://opig.stats.ox.ac.uk/webapps/ngsdb/json/Zhou_2015/Zhou_2015_IGHE_DONORRU08-HeavyChain_iglblastn_igblastn_IGHE.json.gz
wget http://opig.stats.ox.ac.uk/webapps/ngsdb/json/Zhou_2015/Zhou_2015_IGHE_DONOR44-HeavyChain_iglblastn_igblastn_IGHE.json.gz
wget http://opig.stats.ox.ac.uk/webapps/ngsdb/json/Zhou_2015/Zhou_2015_IGHM_DONOR44-HeavyChain_iglblastn_igblastn_IGHM.json.gz
wget http://opig.stats.ox.ac.uk/webapps/ngsdb/json/Zhou_2015/Zhou_2015_IGHD_DONOR44-HeavyChain_iglblastn_igblastn_IGHD.json.gz
wget http://opig.stats.ox.ac.uk/webapps/ngsdb/json/Zhou_2015/Zhou_2015_IGHG_DONORRU01-HeavyChain_iglblastn_igblastn_IGHG.json.gz
wget http://opig.stats.ox.ac.uk/webapps/ngsdb/json/Zhou_2015/Zhou_2015_IGHE_DONORC38-HeavyChain-VRC16_iglblastn_igblastn_IGHE.json.gz
wget http://opig.stats.ox.ac.uk/webapps/ngsdb/json/Zhou_2015/Zhou_2015_IGHG_DONORRU08-HeavyChain_iglblastn_igblastn_IGHG.json.gz
wget http://opig.stats.ox.ac.uk/webapps/ngsdb/json/Zhou_2015/Zhou_2015_Bulk_DONORRU01-HeavyChain_iglblastn_igblastn_Bulk.json.gz
wget http://opig.stats.ox.ac.uk/webapps/ngsdb/json/Zhou_2015/Zhou_2015_Bulk_DONORC38-HeavyChain-VRC16_iglblastn_igblastn_Bulk.json.gz
wget http://opig.stats.ox.ac.uk/webapps/ngsdb/json/Zhou_2015/Zhou_2015_Bulk_DONOR44-HeavyChain_iglblastn_igblastn_Bulk.json.gz
wget http://opig.stats.ox.ac.uk/webapps/ngsdb/json/Zhou_2015/Zhou_2015_IGHE_DONORRU01-HeavyChain_iglblastn_igblastn_IGHE.json.gz
wget http://opig.stats.ox.ac.uk/webapps/ngsdb/json/Zhou_2015/Zhou_2015_IGHA_DONORC38-HeavyChain-VRC16_iglblastn_igblastn_IGHA.json.gz
wget http://opig.stats.ox.ac.uk/webapps/ngsdb/json/Zhou_2015/Zhou_2015_IGHM_DONORRU01-HeavyChain_iglblastn_igblastn_IGHM.json.gz
```
