## Supplementary material for "Hiding in plain sight: structure and sequence analysis reveals the importance of the antibody DE loop for antibody-antigen binding": download_naive_seqs.sh

```
wget http://opig.stats.ox.ac.uk/webapps/ngsdb/json/Vander_Heiden_2017/Vander_Heiden_2017_Heavy_HD07_IGHG_HD07_Naive_Bcells_age33_healthy_iglblastn_igblastn_IGHG.json.gz
wget http://opig.stats.ox.ac.uk/webapps/ngsdb/json/Vander_Heiden_2017/Vander_Heiden_2017_Heavy_HD10_IGHD_HD10_Naive_Bcells_age31_healthy_iglblastn_igblastn_IGHD.json.gz
wget http://opig.stats.ox.ac.uk/webapps/ngsdb/json/Vander_Heiden_2017/Vander_Heiden_2017_Heavy_HD09_Bulk_HD09_Naive_Bcells_age31_healthy_iglblastn_igblastn_Bulk.json.gz
wget http://opig.stats.ox.ac.uk/webapps/ngsdb/json/Vander_Heiden_2017/Vander_Heiden_2017_Heavy_HD10_IGHE_HD10_Naive_Bcells_age31_healthy_iglblastn_igblastn_IGHE.json.gz
wget http://opig.stats.ox.ac.uk/webapps/ngsdb/json/Vander_Heiden_2017/Vander_Heiden_2017_Light_HD10_HD10_Naive_Bcells_age31_healthy_iglblastn.json.gz
wget http://opig.stats.ox.ac.uk/webapps/ngsdb/json/Vander_Heiden_2017/Vander_Heiden_2017_Heavy_HD13_IGHG_HD13_Naive_Bcells_age51_healty_iglblastn_igblastn_IGHG.json.gz
wget http://opig.stats.ox.ac.uk/webapps/ngsdb/json/Vander_Heiden_2017/Vander_Heiden_2017_Light_HD13_HD13_Naive_Bcells_age51_healty_iglblastn.json.gz
wget http://opig.stats.ox.ac.uk/webapps/ngsdb/json/Vander_Heiden_2017/Vander_Heiden_2017_Heavy_HD13_IGHE_HD13_Naive_Bcells_age51_healty_iglblastn_igblastn_IGHE.json.gz
wget http://opig.stats.ox.ac.uk/webapps/ngsdb/json/Vander_Heiden_2017/Vander_Heiden_2017_Light_HD09_HD09_Naive_Bcells_age31_healthy_iglblastn.json.gz
wget http://opig.stats.ox.ac.uk/webapps/ngsdb/json/Vander_Heiden_2017/Vander_Heiden_2017_Heavy_HD07_IGHM_HD07_Naive_Bcells_age33_healthy_iglblastn_igblastn_IGHM.json.gz
wget http://opig.stats.ox.ac.uk/webapps/ngsdb/json/Vander_Heiden_2017/Vander_Heiden_2017_Heavy_HD10_IGHM_HD10_Naive_Bcells_age31_healthy_iglblastn_igblastn_IGHM.json.gz
wget http://opig.stats.ox.ac.uk/webapps/ngsdb/json/Vander_Heiden_2017/Vander_Heiden_2017_Heavy_HD09_IGHD_HD09_Naive_Bcells_age31_healthy_iglblastn_igblastn_IGHD.json.gz
wget http://opig.stats.ox.ac.uk/webapps/ngsdb/json/Vander_Heiden_2017/Vander_Heiden_2017_Heavy_HD09_IGHA_HD09_Naive_Bcells_age31_healthy_iglblastn_igblastn_IGHA.json.gz
wget http://opig.stats.ox.ac.uk/webapps/ngsdb/json/Vander_Heiden_2017/Vander_Heiden_2017_Heavy_HD13_IGHA_HD13_Naive_Bcells_age51_healty_iglblastn_igblastn_IGHA.json.gz
wget http://opig.stats.ox.ac.uk/webapps/ngsdb/json/Vander_Heiden_2017/Vander_Heiden_2017_Heavy_HD07_Bulk_HD07_Naive_Bcells_age33_healthy_iglblastn_igblastn_Bulk.json.gz
wget http://opig.stats.ox.ac.uk/webapps/ngsdb/json/Vander_Heiden_2017/Vander_Heiden_2017_Heavy_HD13_IGHD_HD13_Naive_Bcells_age51_healty_iglblastn_igblastn_IGHD.json.gz
wget http://opig.stats.ox.ac.uk/webapps/ngsdb/json/Vander_Heiden_2017/Vander_Heiden_2017_Heavy_HD13_IGHM_HD13_Naive_Bcells_age51_healty_iglblastn_igblastn_IGHM.json.gz
wget http://opig.stats.ox.ac.uk/webapps/ngsdb/json/Vander_Heiden_2017/Vander_Heiden_2017_Heavy_HD09_IGHE_HD09_Naive_Bcells_age31_healthy_iglblastn_igblastn_IGHE.json.gz
wget http://opig.stats.ox.ac.uk/webapps/ngsdb/json/Vander_Heiden_2017/Vander_Heiden_2017_Heavy_HD09_IGHM_HD09_Naive_Bcells_age31_healthy_iglblastn_igblastn_IGHM.json.gz
wget http://opig.stats.ox.ac.uk/webapps/ngsdb/json/Vander_Heiden_2017/Vander_Heiden_2017_Heavy_HD09_IGHG_HD09_Naive_Bcells_age31_healthy_iglblastn_igblastn_IGHG.json.gz
wget http://opig.stats.ox.ac.uk/webapps/ngsdb/json/Vander_Heiden_2017/Vander_Heiden_2017_Heavy_HD13_Bulk_HD13_Naive_Bcells_age51_healty_iglblastn_igblastn_Bulk.json.gz
wget http://opig.stats.ox.ac.uk/webapps/ngsdb/json/Vander_Heiden_2017/Vander_Heiden_2017_Heavy_HD10_Bulk_HD10_Naive_Bcells_age31_healthy_iglblastn_igblastn_Bulk.json.gz
wget http://opig.stats.ox.ac.uk/webapps/ngsdb/json/Vander_Heiden_2017/Vander_Heiden_2017_Light_HD07_HD07_Naive_Bcells_age33_healthy_iglblastn.json.gz
wget http://opig.stats.ox.ac.uk/webapps/ngsdb/json/Vander_Heiden_2017/Vander_Heiden_2017_Heavy_HD07_IGHE_HD07_Naive_Bcells_age33_healthy_iglblastn_igblastn_IGHE.json.gz
wget http://opig.stats.ox.ac.uk/webapps/ngsdb/json/Vander_Heiden_2017/Vander_Heiden_2017_Heavy_HD10_IGHA_HD10_Naive_Bcells_age31_healthy_iglblastn_igblastn_IGHA.json.gz
wget http://opig.stats.ox.ac.uk/webapps/ngsdb/json/Vander_Heiden_2017/Vander_Heiden_2017_Heavy_HD07_IGHA_HD07_Naive_Bcells_age33_healthy_iglblastn_igblastn_IGHA.json.gz
wget http://opig.stats.ox.ac.uk/webapps/ngsdb/json/Vander_Heiden_2017/Vander_Heiden_2017_Heavy_HD07_IGHD_HD07_Naive_Bcells_age33_healthy_iglblastn_igblastn_IGHD.json.gz
wget http://opig.stats.ox.ac.uk/webapps/ngsdb/json/Vander_Heiden_2017/Vander_Heiden_2017_Heavy_HD10_IGHG_HD10_Naive_Bcells_age31_healthy_iglblastn_igblastn_IGHG.json.gz
wget http://opig.stats.ox.ac.uk/webapps/ngsdb/json/Galson_2015/Galson_2015_Visit-4_Subject-1001_IgM_Subject-1001_Visit-4_treatment-MenACWY-conjugate_Naive-B-cells_IgM_iglblastn.json.gz
wget http://opig.stats.ox.ac.uk/webapps/ngsdb/json/Galson_2015/Galson_2015_Visit-4_Subject-1020_IgM_Subject-1020_Visit-4_treatment-MenACWY-polysaccharide_Naive-B-cells_IgM_iglblastn.json.gz
wget http://opig.stats.ox.ac.uk/webapps/ngsdb/json/Galson_2015/Galson_2015_Visit-1_Subject-1017_IgM_Subject-1017_Visit-1_treatment-MenACWY-polysaccharide_Naive-B-cells_IgM_iglblastn.json.gz
wget http://opig.stats.ox.ac.uk/webapps/ngsdb/json/Galson_2015/Galson_2015_Visit-1_Subject-1011_IgM_Subject-1011_Visit-1_treatment-MenACWY-polysaccharide_Naive-B-cells_IgM_iglblastn.json.gz
wget http://opig.stats.ox.ac.uk/webapps/ngsdb/json/Galson_2015/Galson_2015_Visit-4_Subject-1018_IgM_Subject-1018_Visit-4_treatment-MenACWY-conjugate_Naive-B-cells_IgM_iglblastn.json.gz
wget http://opig.stats.ox.ac.uk/webapps/ngsdb/json/Galson_2015/Galson_2015_Visit-1_Subject-1009_IgM_Subject-1009_Visit-1_treatment-MenACWY-conjugate_Naive-B-cells_IgM_iglblastn.json.gz
wget http://opig.stats.ox.ac.uk/webapps/ngsdb/json/Galson_2015/Galson_2015_Visit-1_Subject-1010_IgM_Subject-1010_Visit-1_treatment-MenACWY-conjugate_Naive-B-cells_IgM_iglblastn.json.gz
wget http://opig.stats.ox.ac.uk/webapps/ngsdb/json/Galson_2015/Galson_2015_Visit-1_Subject-1015_IgM_Subject-1015_Visit-1_treatment-MenACWY-polysaccharide_Naive-B-cells_IgM_iglblastn.json.gz
wget http://opig.stats.ox.ac.uk/webapps/ngsdb/json/Galson_2015/Galson_2015_Visit-4_Subject-1014_IgM_Subject-1014_Visit-4_treatment-MenACWY-polysaccharide_Naive-B-cells_IgM_iglblastn.json.gz
wget http://opig.stats.ox.ac.uk/webapps/ngsdb/json/Johnson_2018/Johnson_2018_Heavy_Healthy_Healthy_Heavy_Healthy_Healthy_Naive_Part-1.json.gz
wget http://opig.stats.ox.ac.uk/webapps/ngsdb/json/Ellebedy_2016/Ellebedy_2016_Day-7_naive-B-cell_IGHG_naive-B-cell_Day-7_Donor-163_igblastn_IGHG.json.gz
wget http://opig.stats.ox.ac.uk/webapps/ngsdb/json/Ellebedy_2016/Ellebedy_2016_Day-0_naive-B-cell_IGHG_naive-B-cell_Day-0_Donor-4_igblastn_IGHG.json.gz
wget http://opig.stats.ox.ac.uk/webapps/ngsdb/json/Ellebedy_2016/Ellebedy_2016_Day-0_naive-B-cell_IGHD_naive-B-cell_Day-0_Donor-5_igblastn_IGHD.json.gz
wget http://opig.stats.ox.ac.uk/webapps/ngsdb/json/Ellebedy_2016/Ellebedy_2016_Day-7_naive-B-cell_IGHG_naive-B-cell_Day-7_Donor-155_igblastn_IGHG.json.gz
wget http://opig.stats.ox.ac.uk/webapps/ngsdb/json/Ellebedy_2016/Ellebedy_2016_Day-0_naive-B-cell_IGHG_naive-B-cell_Day-0_Donor-6_igblastn_IGHG.json.gz
wget http://opig.stats.ox.ac.uk/webapps/ngsdb/json/Ellebedy_2016/Ellebedy_2016_Day-0_naive-B-cell_Bulk_naive-B-cell_Day-0_Donor-5_igblastn_Bulk.json.gz
wget http://opig.stats.ox.ac.uk/webapps/ngsdb/json/Ellebedy_2016/Ellebedy_2016_Day-0_naive-B-cell_IGHM_naive-B-cell_Day-0_Donor-8_igblastn_IGHM.json.gz
wget http://opig.stats.ox.ac.uk/webapps/ngsdb/json/Ellebedy_2016/Ellebedy_2016_Day-0_naive-B-cell_IGHA_naive-B-cell_Day-0_Donor-8_igblastn_IGHA.json.gz
wget http://opig.stats.ox.ac.uk/webapps/ngsdb/json/Ellebedy_2016/Ellebedy_2016_Day-7_naive-B-cell_IGHM_naive-B-cell_Day-7_Donor-162_igblastn_IGHM.json.gz
wget http://opig.stats.ox.ac.uk/webapps/ngsdb/json/Ellebedy_2016/Ellebedy_2016_Day-0_naive-B-cell_IGHM_naive-B-cell_Day-0_Donor-5_igblastn_IGHM.json.gz
wget http://opig.stats.ox.ac.uk/webapps/ngsdb/json/Ellebedy_2016/Ellebedy_2016_Day-0_naive-B-cell_Bulk_naive-B-cell_Day-0_Donor-6_igblastn_Bulk.json.gz
wget http://opig.stats.ox.ac.uk/webapps/ngsdb/json/Ellebedy_2016/Ellebedy_2016_Day-0_naive-B-cell_IGHD_naive-B-cell_Day-0_Donor-6_igblastn_IGHD.json.gz
wget http://opig.stats.ox.ac.uk/webapps/ngsdb/json/Ellebedy_2016/Ellebedy_2016_Day-0_naive-B-cell_IGHM_naive-B-cell_Day-0_Donor-6_igblastn_IGHM.json.gz
wget http://opig.stats.ox.ac.uk/webapps/ngsdb/json/Ellebedy_2016/Ellebedy_2016_Day-7_naive-B-cell_IGHD_naive-B-cell_Day-7_Donor-157_igblastn_IGHD.json.gz
wget http://opig.stats.ox.ac.uk/webapps/ngsdb/json/Ellebedy_2016/Ellebedy_2016_Day-7_naive-B-cell_IGHG_naive-B-cell_Day-7_Donor-162_igblastn_IGHG.json.gz
wget http://opig.stats.ox.ac.uk/webapps/ngsdb/json/Ellebedy_2016/Ellebedy_2016_Day-0_naive-B-cell_IGHA_naive-B-cell_Day-0_Donor-5_igblastn_IGHA.json.gz
wget http://opig.stats.ox.ac.uk/webapps/ngsdb/json/Ellebedy_2016/Ellebedy_2016_Day-7_naive-B-cell_IGHM_naive-B-cell_Day-7_Donor-157_igblastn_IGHM.json.gz
wget http://opig.stats.ox.ac.uk/webapps/ngsdb/json/Ellebedy_2016/Ellebedy_2016_Day-0_naive-B-cell_IGHG_naive-B-cell_Day-0_Donor-8_igblastn_IGHG.json.gz
wget http://opig.stats.ox.ac.uk/webapps/ngsdb/json/Ellebedy_2016/Ellebedy_2016_Day-0_naive-B-cell_IGHG_naive-B-cell_Day-0_Donor-5_igblastn_IGHG.json.gz
wget http://opig.stats.ox.ac.uk/webapps/ngsdb/json/Ellebedy_2016/Ellebedy_2016_Day-0_naive-B-cell_IGHA_naive-B-cell_Day-0_Donor-6_igblastn_IGHA.json.gz
wget http://opig.stats.ox.ac.uk/webapps/ngsdb/json/Ellebedy_2016/Ellebedy_2016_Day-7_naive-B-cell_Bulk_naive-B-cell_Day-7_Donor-163_igblastn_Bulk.json.gz
wget http://opig.stats.ox.ac.uk/webapps/ngsdb/json/Ellebedy_2016/Ellebedy_2016_Day-7_naive-B-cell_IGHM_naive-B-cell_Day-7_Donor-163_igblastn_IGHM.json.gz
wget http://opig.stats.ox.ac.uk/webapps/ngsdb/json/Ellebedy_2016/Ellebedy_2016_Day-0_naive-B-cell_IGHA_naive-B-cell_Day-0_Donor-4_igblastn_IGHA.json.gz
wget http://opig.stats.ox.ac.uk/webapps/ngsdb/json/Ellebedy_2016/Ellebedy_2016_Day-7_naive-B-cell_IGHG_naive-B-cell_Day-7_Donor-157_igblastn_IGHG.json.gz
wget http://opig.stats.ox.ac.uk/webapps/ngsdb/json/Ellebedy_2016/Ellebedy_2016_Day-7_naive-B-cell_IGHM_naive-B-cell_Day-7_Donor-155_igblastn_IGHM.json.gz
wget http://opig.stats.ox.ac.uk/webapps/ngsdb/json/Ellebedy_2016/Ellebedy_2016_Day-0_naive-B-cell_IGHM_naive-B-cell_Day-0_Donor-4_igblastn_IGHM.json.gz
wget http://opig.stats.ox.ac.uk/webapps/ngsdb/json/Turchaninova_2016/Turchaninova_2016_IGHG_B_naive_donor_3_iglblastn_igblastn_IGHG.json.gz
wget http://opig.stats.ox.ac.uk/webapps/ngsdb/json/Turchaninova_2016/Turchaninova_2016_IGHM_B_naive_donor_3_iglblastn_igblastn_IGHM.json.gz
wget http://opig.stats.ox.ac.uk/webapps/ngsdb/json/Turchaninova_2016/Turchaninova_2016_Bulk_B_naive_donor_3_iglblastn_igblastn_Bulk.json.gz
wget http://opig.stats.ox.ac.uk/webapps/ngsdb/json/Turchaninova_2016/Turchaninova_2016_IGHD_B_naive_donor_3_iglblastn_igblastn_IGHD.json.gz
wget http://opig.stats.ox.ac.uk/webapps/ngsdb/json/Turchaninova_2016/Turchaninova_2016_IGHA_B_naive_donor_3_iglblastn_igblastn_IGHA.json.gz
wget http://opig.stats.ox.ac.uk/webapps/ngsdb/json/Rubelt_2016/Rubelt_2016_TWINs_4_TW04B_IGHG_TW04B_B_naive_iglblastn_IGHG.json.gz
wget http://opig.stats.ox.ac.uk/webapps/ngsdb/json/Rubelt_2016/Rubelt_2016_TWINs_4_TW04B_IGHE_TW04B_B_naive_iglblastn_IGHE.json.gz
wget http://opig.stats.ox.ac.uk/webapps/ngsdb/json/Rubelt_2016/Rubelt_2016_TWINs_4_TW04B_IGHA_TW04B_B_naive_iglblastn_IGHA.json.gz
wget http://opig.stats.ox.ac.uk/webapps/ngsdb/json/Rubelt_2016/Rubelt_2016_TWINs_3_TW03B_IGHA_TW03B_B_naive_iglblastn_IGHA.json.gz
wget http://opig.stats.ox.ac.uk/webapps/ngsdb/json/Rubelt_2016/Rubelt_2016_TWINs_3_TW03B_Bulk_TW03B_B_naive_iglblastn_Bulk.json.gz
wget http://opig.stats.ox.ac.uk/webapps/ngsdb/json/Rubelt_2016/Rubelt_2016_TWINs_1_TW01A_IGHA_TW01A_B_naive_iglblastn_IGHA.json.gz
wget http://opig.stats.ox.ac.uk/webapps/ngsdb/json/Rubelt_2016/Rubelt_2016_TWINs_2_TW02B_Bulk_TW02B_B_naive_iglblastn_Bulk.json.gz
wget http://opig.stats.ox.ac.uk/webapps/ngsdb/json/Rubelt_2016/Rubelt_2016_TWINs_2_TW02B_IGHG_TW02B_B_naive_iglblastn_IGHG.json.gz
wget http://opig.stats.ox.ac.uk/webapps/ngsdb/json/Rubelt_2016/Rubelt_2016_TWINs_2_TW02A_IGHG_TW02A_B_naive_iglblastn_IGHG.json.gz
wget http://opig.stats.ox.ac.uk/webapps/ngsdb/json/Rubelt_2016/Rubelt_2016_TWINs_1_TW01A_IGHM_TW01A_B_naive_iglblastn_IGHM.json.gz
wget http://opig.stats.ox.ac.uk/webapps/ngsdb/json/Rubelt_2016/Rubelt_2016_TWINs_3_TW03A_IGHE_TW03A_B_naive_iglblastn_IGHE.json.gz
wget http://opig.stats.ox.ac.uk/webapps/ngsdb/json/Rubelt_2016/Rubelt_2016_TWINs_3_TW03B_IGHE_TW03B_B_naive_iglblastn_IGHE.json.gz
wget http://opig.stats.ox.ac.uk/webapps/ngsdb/json/Rubelt_2016/Rubelt_2016_TWINs_5_TW05A_IGHD_TW05A_B_naive_iglblastn_IGHD.json.gz
wget http://opig.stats.ox.ac.uk/webapps/ngsdb/json/Rubelt_2016/Rubelt_2016_TWINs_2_TW02A_IGHM_TW02A_B_naive_iglblastn_IGHM.json.gz
wget http://opig.stats.ox.ac.uk/webapps/ngsdb/json/Rubelt_2016/Rubelt_2016_TWINs_1_TW01B_IGHA_TW01B_B_naive_iglblastn_IGHA.json.gz
wget http://opig.stats.ox.ac.uk/webapps/ngsdb/json/Rubelt_2016/Rubelt_2016_TWINs_4_TW04A_IGHD_TW04A_B_naive_iglblastn_IGHD.json.gz
wget http://opig.stats.ox.ac.uk/webapps/ngsdb/json/Rubelt_2016/Rubelt_2016_TWINs_5_TW05B_IGHD_TW05B_B_naive_iglblastn_IGHD.json.gz
wget http://opig.stats.ox.ac.uk/webapps/ngsdb/json/Rubelt_2016/Rubelt_2016_TWINs_2_TW02B_IGHA_TW02B_B_naive_iglblastn_IGHA.json.gz
wget http://opig.stats.ox.ac.uk/webapps/ngsdb/json/Rubelt_2016/Rubelt_2016_TWINs_5_TW05A_IGHM_TW05A_B_naive_iglblastn_IGHM.json.gz
wget http://opig.stats.ox.ac.uk/webapps/ngsdb/json/Rubelt_2016/Rubelt_2016_TWINs_2_TW02A_IGHE_TW02A_B_naive_iglblastn_IGHE.json.gz
wget http://opig.stats.ox.ac.uk/webapps/ngsdb/json/Rubelt_2016/Rubelt_2016_TWINs_1_TW01B_Bulk_TW01B_B_naive_iglblastn_Bulk.json.gz
wget http://opig.stats.ox.ac.uk/webapps/ngsdb/json/Rubelt_2016/Rubelt_2016_TWINs_2_TW02A_Bulk_TW02A_B_naive_iglblastn_Bulk.json.gz
wget http://opig.stats.ox.ac.uk/webapps/ngsdb/json/Rubelt_2016/Rubelt_2016_TWINs_1_TW01B_IGHE_TW01B_B_naive_iglblastn_IGHE.json.gz
wget http://opig.stats.ox.ac.uk/webapps/ngsdb/json/Rubelt_2016/Rubelt_2016_TWINs_4_TW04A_IGHM_TW04A_B_naive_iglblastn_IGHM.json.gz
wget http://opig.stats.ox.ac.uk/webapps/ngsdb/json/Rubelt_2016/Rubelt_2016_TWINs_4_TW04A_IGHE_TW04A_B_naive_iglblastn_IGHE.json.gz
wget http://opig.stats.ox.ac.uk/webapps/ngsdb/json/Rubelt_2016/Rubelt_2016_TWINs_4_TW04B_Bulk_TW04B_B_naive_iglblastn_Bulk.json.gz
wget http://opig.stats.ox.ac.uk/webapps/ngsdb/json/Rubelt_2016/Rubelt_2016_TWINs_2_TW02B_IGHM_TW02B_B_naive_iglblastn_IGHM.json.gz
wget http://opig.stats.ox.ac.uk/webapps/ngsdb/json/Rubelt_2016/Rubelt_2016_TWINs_1_TW01B_IGHM_TW01B_B_naive_iglblastn_IGHM.json.gz
wget http://opig.stats.ox.ac.uk/webapps/ngsdb/json/Rubelt_2016/Rubelt_2016_TWINs_4_TW04A_IGHG_TW04A_B_naive_iglblastn_IGHG.json.gz
wget http://opig.stats.ox.ac.uk/webapps/ngsdb/json/Rubelt_2016/Rubelt_2016_TWINs_3_TW03A_IGHA_TW03A_B_naive_iglblastn_IGHA.json.gz
wget http://opig.stats.ox.ac.uk/webapps/ngsdb/json/Rubelt_2016/Rubelt_2016_TWINs_2_TW02B_IGHD_TW02B_B_naive_iglblastn_IGHD.json.gz
wget http://opig.stats.ox.ac.uk/webapps/ngsdb/json/Rubelt_2016/Rubelt_2016_TWINs_5_TW05A_IGHE_TW05A_B_naive_iglblastn_IGHE.json.gz
wget http://opig.stats.ox.ac.uk/webapps/ngsdb/json/Rubelt_2016/Rubelt_2016_TWINs_4_TW04A_Bulk_TW04A_B_naive_iglblastn_Bulk.json.gz
wget http://opig.stats.ox.ac.uk/webapps/ngsdb/json/Rubelt_2016/Rubelt_2016_TWINs_2_TW02A_IGHD_TW02A_B_naive_iglblastn_IGHD.json.gz
wget http://opig.stats.ox.ac.uk/webapps/ngsdb/json/Rubelt_2016/Rubelt_2016_TWINs_1_TW01A_Bulk_TW01A_B_naive_iglblastn_Bulk.json.gz
wget http://opig.stats.ox.ac.uk/webapps/ngsdb/json/Rubelt_2016/Rubelt_2016_TWINs_3_TW03B_IGHD_TW03B_B_naive_iglblastn_IGHD.json.gz
wget http://opig.stats.ox.ac.uk/webapps/ngsdb/json/Rubelt_2016/Rubelt_2016_TWINs_1_TW01A_IGHG_TW01A_B_naive_iglblastn_IGHG.json.gz
wget http://opig.stats.ox.ac.uk/webapps/ngsdb/json/Rubelt_2016/Rubelt_2016_TWINs_5_TW05B_IGHA_TW05B_B_naive_iglblastn_IGHA.json.gz
wget http://opig.stats.ox.ac.uk/webapps/ngsdb/json/Rubelt_2016/Rubelt_2016_TWINs_3_TW03A_Bulk_TW03A_B_naive_iglblastn_Bulk.json.gz
wget http://opig.stats.ox.ac.uk/webapps/ngsdb/json/Rubelt_2016/Rubelt_2016_TWINs_5_TW05B_IGHG_TW05B_B_naive_iglblastn_IGHG.json.gz
wget http://opig.stats.ox.ac.uk/webapps/ngsdb/json/Rubelt_2016/Rubelt_2016_TWINs_1_TW01B_IGHG_TW01B_B_naive_iglblastn_IGHG.json.gz
wget http://opig.stats.ox.ac.uk/webapps/ngsdb/json/Rubelt_2016/Rubelt_2016_TWINs_5_TW05B_IGHE_TW05B_B_naive_iglblastn_IGHE.json.gz
wget http://opig.stats.ox.ac.uk/webapps/ngsdb/json/Rubelt_2016/Rubelt_2016_TWINs_5_TW05A_Bulk_TW05A_B_naive_iglblastn_Bulk.json.gz
wget http://opig.stats.ox.ac.uk/webapps/ngsdb/json/Rubelt_2016/Rubelt_2016_TWINs_3_TW03B_IGHG_TW03B_B_naive_iglblastn_IGHG.json.gz
wget http://opig.stats.ox.ac.uk/webapps/ngsdb/json/Rubelt_2016/Rubelt_2016_TWINs_2_TW02A_IGHA_TW02A_B_naive_iglblastn_IGHA.json.gz
wget http://opig.stats.ox.ac.uk/webapps/ngsdb/json/Rubelt_2016/Rubelt_2016_TWINs_5_TW05B_Bulk_TW05B_B_naive_iglblastn_Bulk.json.gz
wget http://opig.stats.ox.ac.uk/webapps/ngsdb/json/Rubelt_2016/Rubelt_2016_TWINs_5_TW05A_IGHG_TW05A_B_naive_iglblastn_IGHG.json.gz
wget http://opig.stats.ox.ac.uk/webapps/ngsdb/json/Rubelt_2016/Rubelt_2016_TWINs_5_TW05B_IGHM_TW05B_B_naive_iglblastn_IGHM.json.gz
wget http://opig.stats.ox.ac.uk/webapps/ngsdb/json/Rubelt_2016/Rubelt_2016_TWINs_3_TW03A_IGHD_TW03A_B_naive_iglblastn_IGHD.json.gz
wget http://opig.stats.ox.ac.uk/webapps/ngsdb/json/Rubelt_2016/Rubelt_2016_TWINs_3_TW03B_IGHM_TW03B_B_naive_iglblastn_IGHM.json.gz
wget http://opig.stats.ox.ac.uk/webapps/ngsdb/json/Rubelt_2016/Rubelt_2016_TWINs_1_TW01B_IGHD_TW01B_B_naive_iglblastn_IGHD.json.gz
wget http://opig.stats.ox.ac.uk/webapps/ngsdb/json/Rubelt_2016/Rubelt_2016_TWINs_3_TW03A_IGHM_TW03A_B_naive_iglblastn_IGHM.json.gz
wget http://opig.stats.ox.ac.uk/webapps/ngsdb/json/Rubelt_2016/Rubelt_2016_TWINs_4_TW04B_IGHD_TW04B_B_naive_iglblastn_IGHD.json.gz
wget http://opig.stats.ox.ac.uk/webapps/ngsdb/json/Rubelt_2016/Rubelt_2016_TWINs_1_TW01A_IGHD_TW01A_B_naive_iglblastn_IGHD.json.gz
wget http://opig.stats.ox.ac.uk/webapps/ngsdb/json/Rubelt_2016/Rubelt_2016_TWINs_3_TW03A_IGHG_TW03A_B_naive_iglblastn_IGHG.json.gz
wget http://opig.stats.ox.ac.uk/webapps/ngsdb/json/Rubelt_2016/Rubelt_2016_TWINs_4_TW04B_IGHM_TW04B_B_naive_iglblastn_IGHM.json.gz
wget http://opig.stats.ox.ac.uk/webapps/ngsdb/json/Rubelt_2016/Rubelt_2016_TWINs_5_TW05A_IGHA_TW05A_B_naive_iglblastn_IGHA.json.gz
wget http://opig.stats.ox.ac.uk/webapps/ngsdb/json/Rubelt_2016/Rubelt_2016_TWINs_4_TW04A_IGHA_TW04A_B_naive_iglblastn_IGHA.json.gz
```
